# Brain-wide associations between white matter and age highlight the role of fornix microstructure in brain ageing

**DOI:** 10.1101/2022.09.29.510029

**Authors:** Max Korbmacher, Ann Marie de Lange, Dennis van der Meer, Dani Beck, Eli Eikefjord, Arvid Lundervold, Ole A. Andreassen, Lars T. Westlye, Ivan I. Maximov

## Abstract

Unveiling the details of white matter (WM) maturation throughout ageing is a fundamental question for understanding the ageing brain. In an extensive comparison of brain age predictions and age-associations of WM features from different diffusion approaches, we analysed UK Biobank diffusion Magnetic Resonance Imaging (dMRI) data across midlife and older age (*N* = 35,749, 44.6 to 82.8 years of age). Conventional and advanced dMRI approaches were consistent in predicting brain age. WM-age associations indicate a steady microstructure degeneration with increasing age from midlife to older ages. Brain age was estimated best when combining diffusion approaches, showing different aspects of WM contributing to brain age. Fornix was found as the central region for brain age predictions across diffusion approaches in complement to forceps minor as another important region. These regions exhibited a general pattern of positive associations with age for intra axonal water fractions, axial, radial diffusivities and negative relationships with age for mean diffusivities, fractional anisotropy, kurtosis. We encourage the application of multiple dMRI approaches for detailed insights into WM, and the further investigation of fornix and forceps as potential biomarkers of brain age and ageing.

## Introduction

Neuroscientific research over the past decades has increased our understanding of the biological mechanisms associated with brain tissue maturation and ageing effects^1^. In that endevour, magnetic resonance imaging (MRI) has proven itself as a useful source of data, revealing information about structural and functional brain architecture in vivo^2^. Different MRI modalities, such as diffusion- weighted MRI (dMRI) or T_1_-weighted MRI, allow for the estimation of a variety of quantitative measures, which can then be linked to behaviour, cognitive and health scores^3, 4^. However, intra- subject variability in ageing, for example influenced by covariates from the genetic to environmental level^4^ and limitations in sample size, can limit the findings’ interpretability. The use of large-scale MRI databases, such as UK Biobank (UKB)^5^ or the Human Connectome Project^6^, supports the detecting and localising of important brain patterns while providing larger generalisability^7^. Simultaneously, large-scale data provides sufficient power for the application of advanced multivariate statistical models, and machine learning (ML) techniques.

Brain age prediction is an example of such a technique, helping translate large amounts of complex multidimensional data into practically interpretable outputs. Brain age prediction involves training a ML model to determine trajectories of brain ageing from a series of brain MRI features. Once the model is trained, it can predict the age of brains not included in the training data. The disparity between chronological age and predicted age, the so-called brain age gap (BAG), can be used as an indicator for neurological, neuropsychiatric and neurodegenerative disorders^10, 11^. For example, BAG has been associated with stroke history, diabetes, smoking, alcohol intake, several cognitive measures^12, 13^, mortality risk, different brain and psychiatric disorders^14, 15^, cardiovascular risk factors^19^, stroke risk^16^, and loneliness^17^. However, besides Alzheimer’s disease or schizophrenia, the evidence is mixed for the relationship of BAG and different health outcomes and a smaller BAG is not necessarily indicative of good health^4^. Moreover, recent longitudinal evidence shows early-life factors and genetics to have stronger effects on brain maturation than T_1_-weighted grey matter (GM) BAG^18^. However, BAG is a promising heritable indicator of general health status^10, 13, 19, 20^.

BAG and age trajectories offer paths towards a better understanding of the ageing brain. There are various detectable age-related brain changes, such as GM and WM atrophy^8^, WM de- differentiation^9^, and functional connectivity changes^4^ which have hence informed the choice of brain-age modelling-parameters^12, 16, 19, 25, 27–30^. In that context, many ML approaches have been used to make robust and clinically relevant brain age predictions from different MRI modalities^10, 21–23;^ yet, particularly the eXtreme Gradient Boosting^24^ regressor model, using a decision tree approach, being increasingly used for brain age predictions from large-scale data due to its precision and speed ^10, 25, 26^. Especially dMRI and structural MRI have been shown useful for brain age predictions^12, 16, 19, 25, 27–29, 30^. However, further systematic, sufficiently powered assessments of dMRI-derived brain age and how diffusion metrics map onto age are needed.

DMRI-derived measures consist of unique parameters allowing both to reveal WM changes at micrometer scale and to provide the basis for a prediction of macroscopic outcomes, such as age. Conventionally, WM brain architecture is described using diffusion tensor imaging (DTI)^31^.

However, recent advances offer more biophysically meaningful approaches^32^, and sensible foundation for cross-validation and better comparability^25^. DTI-derived measures, namely fractional anisotropy (FA), and axial (AD), mean (MD), and radial (RD) diffusivity have all been shown to be highly age sensitive^9, 25, 33^. Nevertheless, the DTI approach is limited by the Gaussian diffusion assumption and is unable to take into account entangled WM microstructure features^25^. In the present work, we consider 1) the Bayesian rotationally invariant approach (BRIA)^34^, 2) diffusion kurtosis imaging (DKI)^35^; 3) kurtosis derived supplement, known as white matter tract integrity (WMTI)^36^; 4) spherical mean technique (SMT)^37^, and 5) multi-compartment spherical mean technique (mcSMT)^38^ in addition to DTI. Only a few studies have compared dMRI models directly as original brain age predictors^25, 39, 40^. Yet, brain age and age curve assessments of DTI, BRIA, DKI, WMTI, SMT, mcSMT (**ST10**) in a representative sample present a great interest, as well as most influential WM regions for brain ageing. Our assessments focus on the process of ageing (from midlife to late adulthood), starting by associating BAG across diffusion approaches and compare - predicted vs chronological-age correlations in order to assess predictors’ consistency. As fornix was identified as most contributing feature in these predictions, and forceps minor as another influential region, post-hoc analyses focussed on both fornix, forceps minor and whole-brain relationships with age. Fornix was the strongest correlate of age, and fornix and forceps minor features were highly correlated across approaches. Finally, we created fornix, forceps minor, and whole-brain-age curves expecting curvilinear relationships reflecting brain-tissue-composition at different ageing _stages25,33,52._

## Methods

### Sample characteristics

The original UKB^5^ diffusion MRI data consisted of N = 42,208 participants. After exclusions, based on later withdrawn consent and an ICD-10 diagnosis from categories F, G, I, and stroke (excluded: N = 3,521), and data sets not meeting quality control standards (N = 2,938) using the YTTRIUM method^39^, we obtained a final sample consisting of 35,749 healthy adults (age range 44.57 to 82.75, M_age_ = 64.46, SD_age_ = 7.62, Md_age_ = 64.97; 52.96% females, 47.04% males). In brief, YTTRIUM converts diffusion scalar metric into 2D format using a structural similarity extension^94^ of each scalar map to their mean image in order to create a 2D distribution of image and diffusion parameters. The quality check is based on a 2-step clustering algorithm applied to identify subjects located out of the main distribution. We define healthy here as the absence of mental and behavioural disorder (ICD-10 category F), disease of the nervous system (ICD-10 category G), and disease of the circulatory system (ICD-10 category I). Included participants showed generally higher cognitive test performance and took less medication than excluded subjects (**Table 1**).Participants were recruited and scanned at four different sites: 57.62% in Cheadle, 26.30% in Newcastle, 15.96% in Reading, and 0.12% in Bristol (**Fig.1**). Imbalances in age distributions in the Bristol sample can be attributed to the small number of participants sampled (N = 43).

**Fig. 1:**
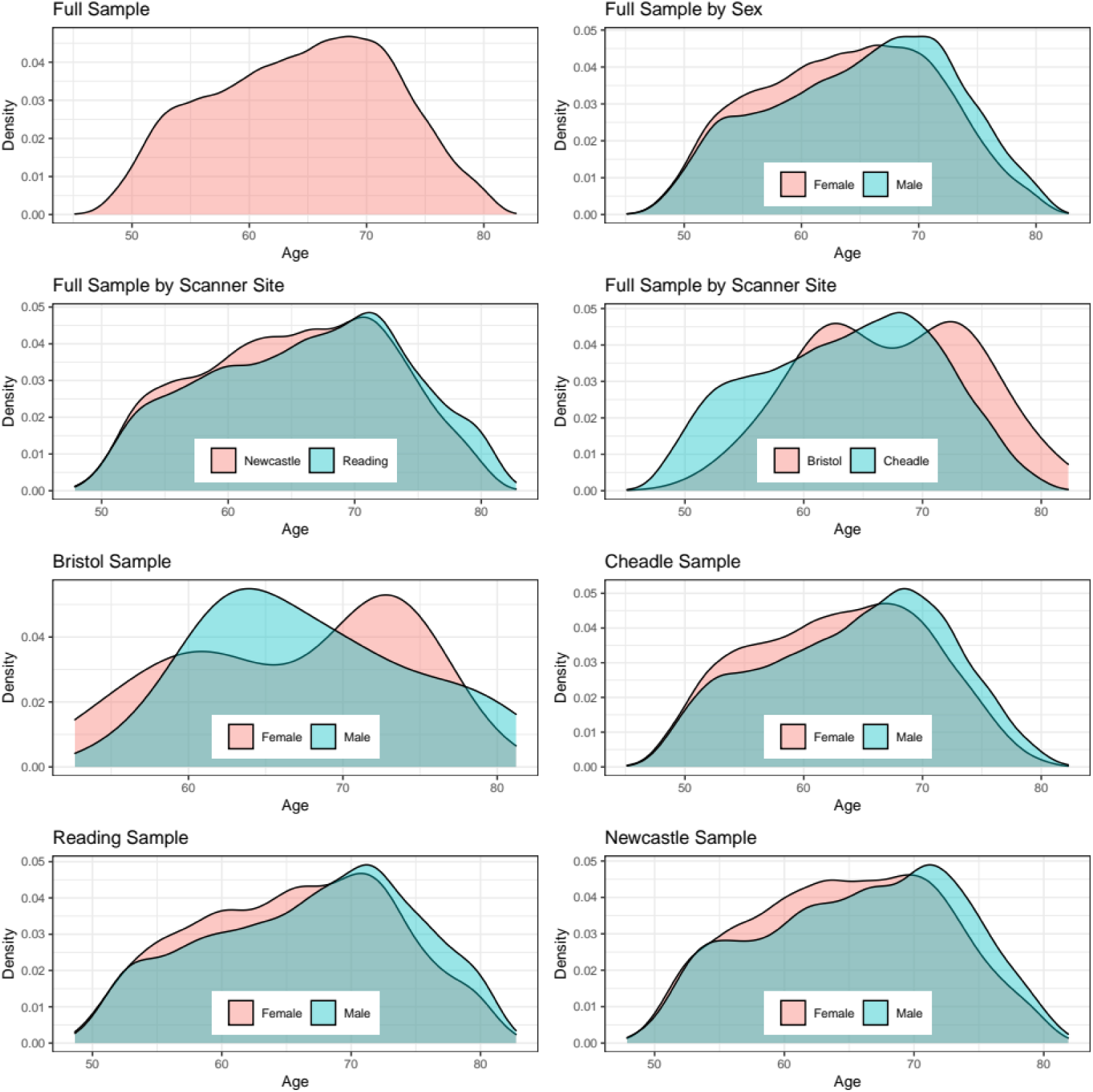
Density plots for the sample’s age by sex and scanner site. The y-axis indicates the probability of age scaled to 1.

**Table 1:** Included and Excluded Sample Characteristics.

| Variable | Excluded<br>(N = 6459) | Included<br>(N = 35,749) | P-value | Cohens *d* |
| --- | --- | --- | --- | --- |
| Number of Medications | 2.812 (2.782) | 1.784 (2.034) | <.001 | 0.474 |
| Self-Rated Health | 2.204 (0.764) | 1.965 (0.644) | <.001 | 0.360 |
| Number of Correctly Solved Matrix Puzzles | 7.671 (2.191) | 8.012 (2.126) | <.001 | -0.159 |
| Number of Correctly Solved Tower Puzzles | 9.650 (3.318) | 9.917 (3.224) | <.001 | -0.083 |
| Number of Correct Symbol Digit Matches | 17.808 (5.414) | 18.998 (5.246) | <.001 | -0.226 |
| Number of Incorrectly Matched Pairs | 2.239 (1.282) | 2.215 (1.274) | 0.250 | 0.019 |
| Matrix Puzzle Response Time in Seconds | 81.116 (16.605) | 83.011 (15.873) | <.001 | -0.119 |
| Maximum Number of Remembered Digits | 6.497 (1.642) | 6.678 (1.538) | <.001 | -0.117 |
| Fluid Intelligence | 6.429 (2.096) | 6.634 (2.054) | <.001 | -0.099 |
| Prospective Memory Score | 1.069 (0.433) | 1.068 (0.397) | 0.783 | 0.004 |

Table logic: Mean (SD) for each sample's variables. *P*-values are indicated for Welch two sample t-tests.

### MRI acquisition, diffusion pipeline and TBSS analysis

UKB MRI data acquisition procedures are described elsewhere^5, 90^. The brain scan protocol (https://biobank.ctsu.ox.ac.uk/crystal/refer.cgi?id=2367) was applied at each scanner site (see also documentation: https://biobank.ctsu.ox.ac.uk/crystal/refer.cgi?id=1977). Shortly, the diffusion protocol consists of 2 b-values (1000 and 2000 s/mm^2^) with 50 non-coplanar diffusion weighting gradients per each shell. For a susceptibility artefact correction, non-diffusion weighted images with an opposite gradient encoding direction were acquited as well.

Diffusion data preprocessing was conducted as described in Maximov et al.^71^, using an optimised pipeline which includes corrections for noise^72^, Gibbs ringing^73^, susceptibility-induced and motion distortions, and eddy current artefacts^74^. Isotropic Gaussian smoothing was carried out with the FSL^75^ function *fslmaths* with a Gaussian kernel of 1 mm^3^. After that DTI, DKI, and WMTI metrics were estimated using Matlab 2017b^76^. Employing the multi-shell data, DKI and WMTI metrics were estimated using Matlab code (https://github.com/NYU-DiffusionMRI/DESIGNER)^36^. SMT, and mcSMT metrics were estimated using original code (https://github.com/ekaden/smt)^37^, as well as Bayesian estimates / BRIA were estimated by the original Matlab code (https://bitbucket.org/reisert/baydiff/src/master/)^34^.

In total, we obtained 28 metrics from six diffusion approaches (DTI, DKI, WMTI, SMT, mcSMT, BRIA)^25, 38, 71, 77–79^. In order to normalise all metrics, we used tract-based spatial statistics (TBSS)^80^, as part of FSL^81^. In brief, initially all BET-extracted^82^ FA images were aligned to MNI space using non-linear transformation (FNIRT)^75^. Afterwards, the mean FA image and related mean FA skeleton were derived. Each diffusion scalar map was projected onto the mean FA skeleton using the TBSS procedure. In order to provide a quantitative description of diffusion metrics we evaluated averaged values over the skeleton and two white matter atlases, namely the JHU atlas^83^ and the JHU tractographic atlas^84^. Finally, we obtained 20 WM tracts and 48 regions of interest (ROIs) based on a probabilistic white matter atlas (JHU) (Hua et al., 2008) for each of the 28 metrics, including the mean skeleton values. Altogether, 1932 features per individual were derived (28 metrics * (48 ROIs + 1 skeleton mean + 20 tracts); see number of dMRI features in **Table 2**)). We included both whole- brain average metrics in addition to tracts and regional averages, as these provide spatially differential information (**SF16**), also expressed the metrics’ relationships with age^25, 33, 91–93^.

**Table 2:** Performance of Brain Age Prediction Models.

| Approach<sup>§</sup> | Number of MRI features | R<sup>2</sup> (SD) | RMSE (SD) | MAE (SD) | Prediction-Age Correlation* [95% CI] | Corrected Prediction-Age Correlation* [95% CI] |
| --- | --- | --- | --- | --- | --- | --- |
| BRIA | 690 | 0.550 (0.012) | 5.007 (0.057) | 4.002 (0.042) | 0.742 [0.737, 0.747] | 0.892<sup>+</sup> [0.889, 0.894] |
| DKI | 207 | 0.576 (0.015) | 4.958 (0.077) | 3.975 (0.068) | 0.754 [0.755, 0.764] | 0.903 [0.901, 0.905] |
| DTI | 276 | 0.571 (0.014) | 4.983 (0.072) | 3.984 (0.062) | 0.756 [0.751, 0.761] | 0.900 [0.897, 0.902] |
| SMT | 276 | 0.531 (0.010) | 5.214 (0.053) | 4.183 (0.036) | 0.729 [0.724, 0.734] | 0.899 [0.897, 0.901] |
| mcSMT | 276 | 0.519 (0.011) | 5.175 (0.045) | 4.153 (0.036) | 0.721 [0.716, 0.726] | 0.892<sup>+</sup> [0.889, 0.894] |
| WMTI | 207 | 0.585 (0.012) | 4.903 (0.065) | 3.928 (0.050) | 0.765 [0.761, 0.770] | 0.902 [0.900, 0.904] |
| Mean multimodal | 28 | 0.393 (0.012) | 5.932 (0.051) | 4.812 (0.046) | 0.627 [0.621, 0.634] | 0.905 [0.903, 0.907] |
| Full multimodal | 1932 | 0.645 (0.011) | 4.534 (0.041) | 3.624 (0.037) | 0.804 [0.800, 0.808] | 0.907 [0.905, 0.909] |

*Table logic:* R<sup>2</sup>, RMSE, MAE are displayed in the format Mean (Standard Deviation), Pearson's correlations are displayed in the format Correlation Score 95% Confidence Interval [Lower Bound, Upper Bound].
Mean multimodal refers to diffusion metrics averaged over the skeleton for all six diffusion approaches. Full multimodal refers to all diffusion data from the six diffusion approaches, i.e. mean multimodal data in addition to metrics averaged over the JHU atlas regions. R<sup>2</sup> = variance explained, RMSE = root mean squared error, MAE = mean absolute error.
§ For an overview of the metrics contained in each of the diffusion approaches see **ST10**.
+ Details on the smallest correlation: BRIA Corrected Prediction-Age Correlation $r = .89173$ , mcSMT Corrected Prediction-Age Correlation $r = .89176$
\* All correlation were significant at $p < .001$ .

### Brain Age Predictions

*First*, brain age predictions were performed using XGBoost^24^ in Python (v3.7.1). To evaluate how much data was needed for hyper-parameter tuning while accurately predicting brain age from all 1932 brain features, we divided the full dataset (N=35,749) into two equal parts: one validation set and one hyper-parameter tuning set for independent parameter-tuning. From the hyper-parameter tuning set, data was randomly sampled into sub-samples consisting of 358, 715, 1,073, 1,430, 1,788, 2,145, 2,503, 2,860, 3,218, 3,575, 7,150, 10,725, 14,300, or 17,875 participants, corresponding to 1%, 2%, 3%, 4%, 5%, 6%, 7%, 8%, 9%, 10%, 20%, 30%, 40% and 50% of the total subjects, respectively (**Fig.2**). Hyper-parameters were tuned on these sub-samples and then tested on the remaining half, i.e., the validation sample, using 10-fold cross validation showing model performance to not further improve past the 10% (tuning) data mark, informing our tuning- validation-split (**Fig.2, ST1**).

**Fig. 2:**
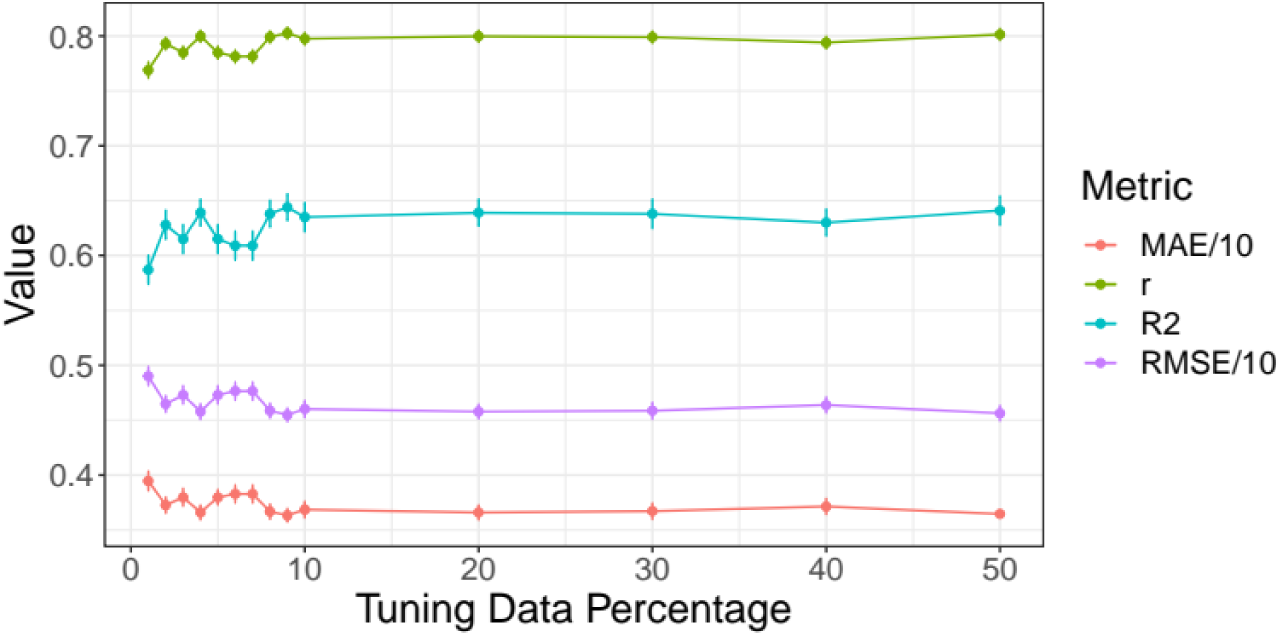
Model performance for different train-test splits.

Model metrics R^2^, RMSE, MAE and their standard deviations, as well as the Pearson’s correlations between predicted and chronological age and its 95% confidence interval are displayed for different training data percentages of the total data (x-axis). For visualisation purposes, RMSE and MAE were divided by 10. For exact values see Suppl. Table ST1. *Second*, in order to compare the different diffusion approaches, based on the previous steps, the training-test split was fixed at previously used 10% training data (N = 3,575) and 90% test data (N = 32,174) which indicated a best fit at a learning rate = 0.05, max layers/depth = 3 and number of trees = 750. These tuned parameters were used for 10-fold cross-validations brain age predictions on the test data of all six individual models, one multimodal model combining all metrics from all diffusion models, and one multimodal model using only mean values from all diffusion models (**Table 2**).

*Third*, uncorrected BAG was calculated as the difference between chronological age Ω and predicted age P:

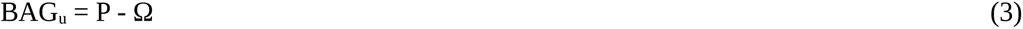

As a supplement, age-bias-corrected predicted age was calculated from the intercept and slope of age predictions as previously described^26, 85^:

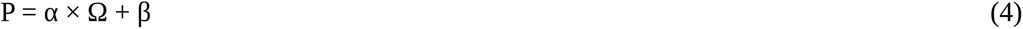

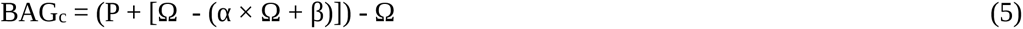

P represents predicted age modelled from chronological age Ω, with intercept β and slope α. This age-bias correction allowed for a bias-corrected BAG estimate (BAG_c_).

### Statistical Analyses

All statistical analyses were carried out using R (v3.6.0; www.r-project.org/). *P*-values were adjusted for multiple comparison using Holm correction^44^. Model performance for brain ages estimations across different diffusion approaches are presented in addition to top five features for each brain age model ranked based on their model contributions (variance explained, as determined by permutation feature importance testing). Then, the correlation structure of age, brain age, BAG, and brain features (identified as main contributors in the model and whole-brain-average scores) were examined across diffusion approaches. In detail: *First*, brain ages were correlated across diffusion-approach-specific brain ages. Then, the correlations between true and estimated age across diffuson approaches were compared. *Second*, BAGs were correlated across diffusion approaches.

*Third*, we present the correlation structure of fornix and age, and present brain-age crude and adjusted age-relationships for all included metrics (M).

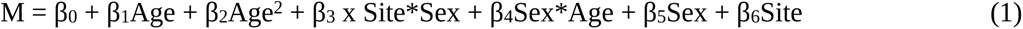

*Fourth*, we plot absolute/crude whole-brain and fornix diffusion metrics by age, and contrast these with diffusion metrics (M) adjusted for age, sex and site. To test the age-sensitivity of the metrics, we removed age from the model and compared the models using Likelihood Ratio tests.

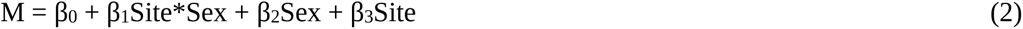

We also assess to which extent the regression lines can be called linear by comparing model fit of gerneralized additive models with simple linear regression models for fornix and whole brain features. Finally, we associate the first two principal components of all WM features with the different brain ages to assess the relationship between BAG and WM. For an overview of the analyses see **Fig.3**.

**Fig. 3:**
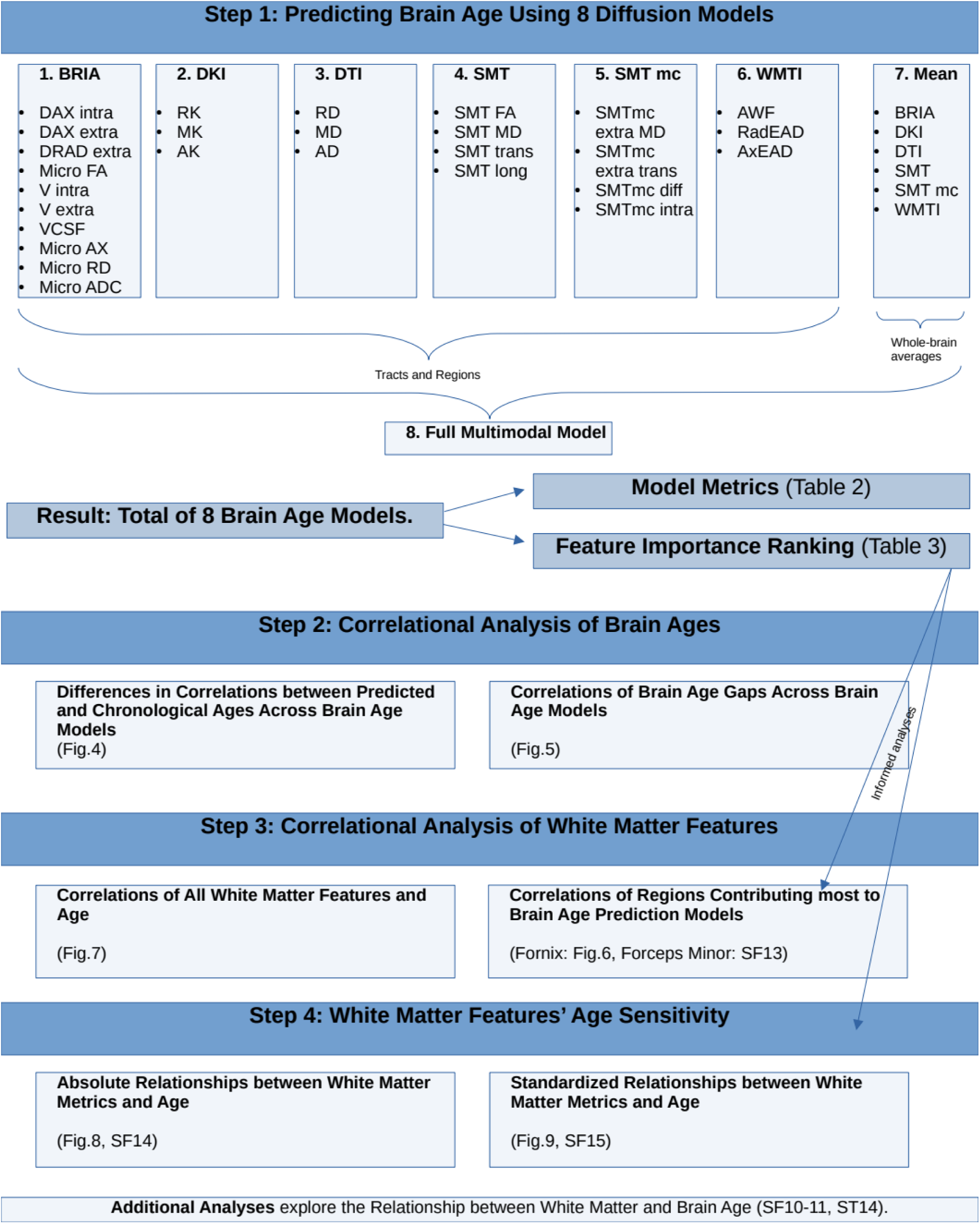
Overview of the Analysis Steps.

## Results

### Brain age predictions

**Table 2** presents a comparison between different diffusion approaches in predicting brain age for each diffusion approach. The strongest correlation between uncorrected age predictions and chronological age was observed for WMTI Pearson’s *r*=0.765, 95% CI [0.761, 0.770], *p*<.001, and the smallest for mcSMT Pearson’s *r*=0.721, 95% CI [0.716, 0.726], *p*<.001.

Hotelling’s^41^ *t*-tests were used to compare correlations between uncorrected predicted age and chronological age across diffusion models. Zou’s^42^ method was used to estimate the confidence intervals around the correlation differences (**Fig.4** and **ST3**; **SF8** and **ST2** for corrected prediction correlation comparisons). These differences were not significantly different from each other for model pairs DKI and DTI (*p≈*1). All other correlations were different from each other, Pearson’s *rs*_diff_≤0.15, *p*<.001, with the biggest difference observed between mean and full multimodal scores’ correlations (**ST2** for exact values).

**Fig. 4:**
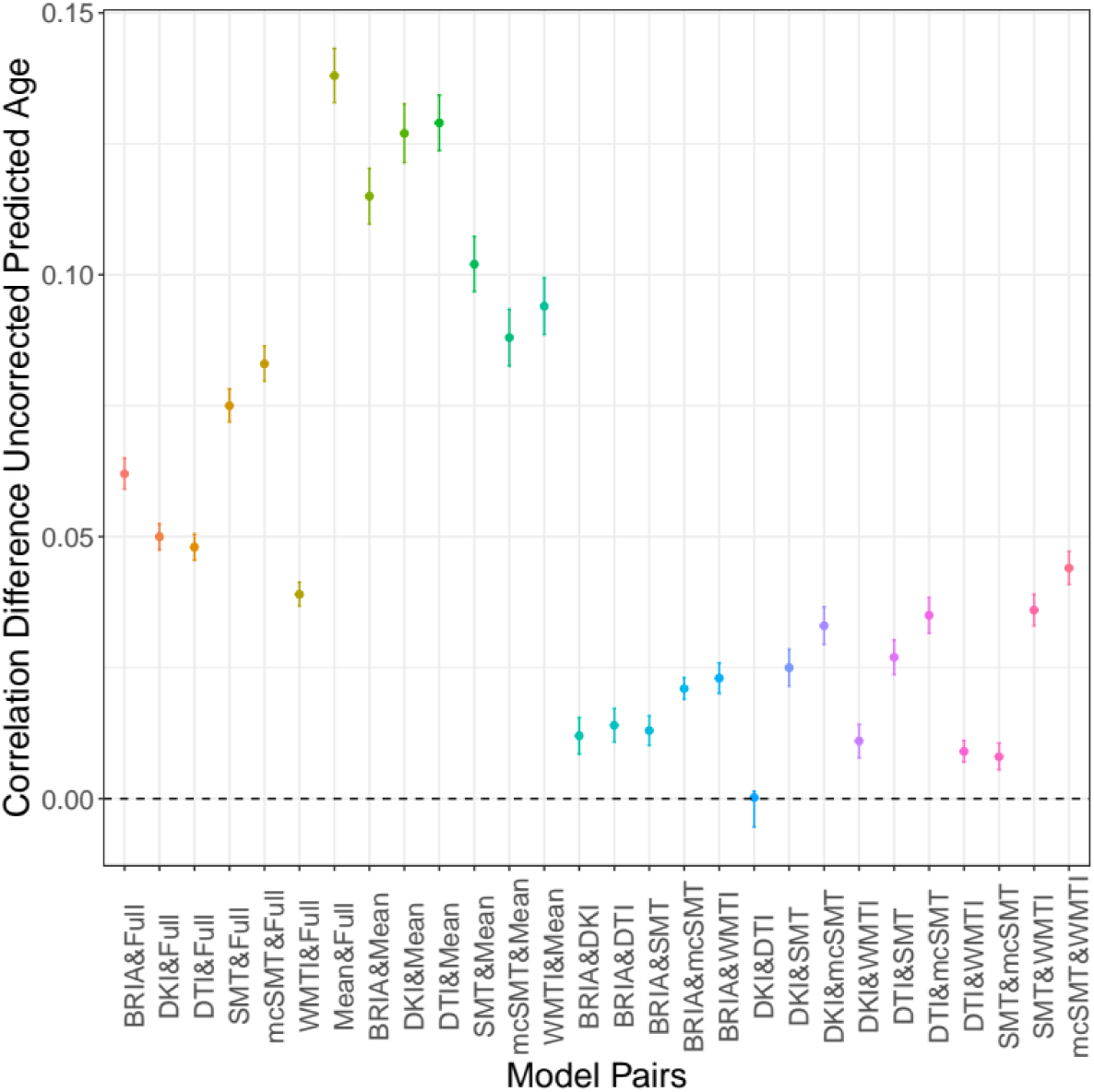
Differences between Pearson’s correlations of chronological and uncorrected predicted ages across diffusion approaches with 95% confidence interval. Differences between Pearson’s correlation coefficients of chronological and uncorrected predicted age by diffusion approach. See SF8 for correlational differences between approaches for corrected brain age predictions.

Permutation feature importance estimates across diffusion models showed that fornix contributed strongest to variance explained (**Table 3**), which was in correspondence with feature rankings by gain score^43^ (**ST15**). Follow-up models which had fornix features removed had lower model fit, explained less variance in age, and predicted-chronological-age correlations were smaller than for models containing fornix (*rsdiff*<-0.003, *ps*<.001; **ST16**). Another potentially important region was the forceps minor, also contributing significantly to age predictions (**Table 3**).

**Table 3:**
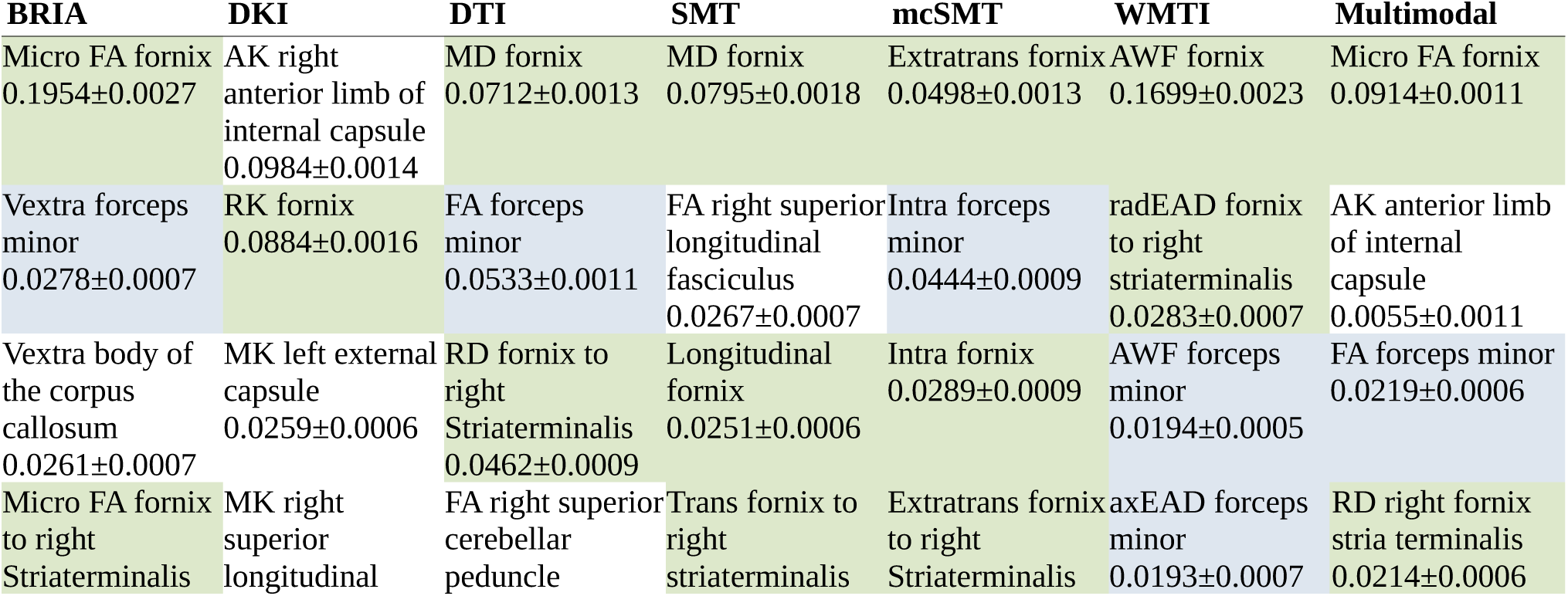
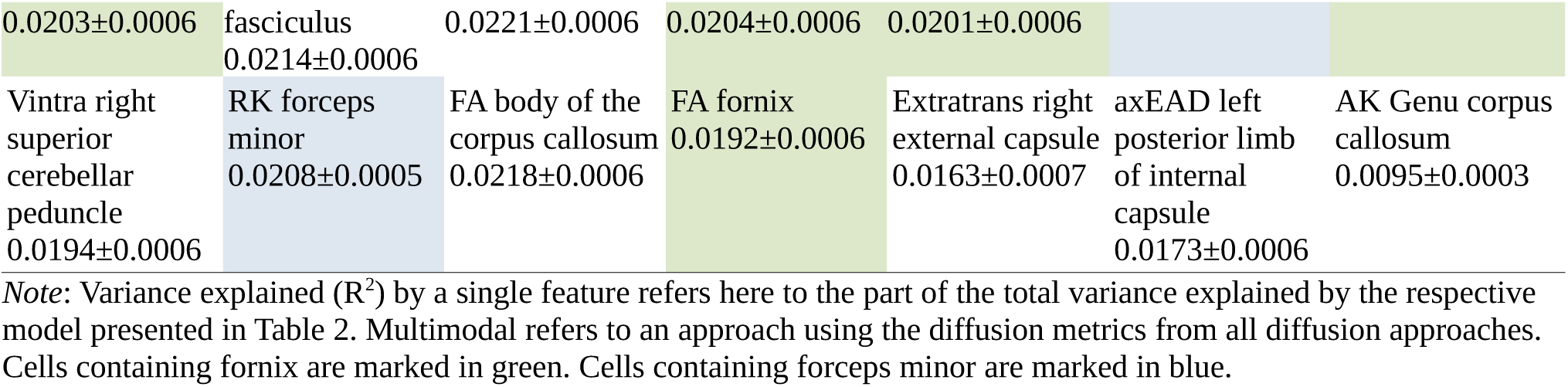
Top five diffusion metrics ranked by their contribution to variance explained (R^2^) in age.

### Brain age gap across diffusion approaches and age

In order to compare uncorrected BAG (BAG_u_) calculations across the used diffusion approaches, BAG_u_ was correlated from different diffusion approaches and with age. Correlations between the six diffusion approaches ranged between *r*=0.857 and *r*=0.966 (**Fig.9; SF1** for corrected BAG correlations). Overall, BAG_u_ scores from the different approaches were strongest related to WMTI BAG_c_ (range: *r* = 0.873 to 0.952), and weakest to mean multimodal BAG_u_ (range: *r*=0.779 to *r*=0.828), and could be observed in one cluster containing DKI, DTI, WMTI and multimodal BAG_u_ and a second cluster containing BRIA, SMT, and SMTmc. However, DKI, BAG_u_ was more strongly correlated with full multimodal BAG_c_ than with other well-performing approaches DTI (Pearson’s *r*_diff_=0.03, *p*<.001) and WMTI (*r*_diff_=0.03, *p*<.001). Vice versa, DTI BAG_c_ correlated strongest with WMTI BAG_c_ (*r*=0.905, *p*<.001).

### Associations between diffusion metrics and age

A correlational analysis was used to demonstrate associations among fornix diffusion metrics and age (**Fig.6**, including QC outliers: **SF4**). Association strengths ranged from to r=-0.997 (smtTrans and smtMCintra) to *r*=0.999 (smtTrans and smtMD). Correlations between fornix metrics and age ranged from *r*=-0.558 (smtMCintra) to *r*=0.570 (microRD), and between forceps minor metrics and age from *r* = -0.519 (FA) to *r* = 0.493 (RD, see **SF13**).

Correlations across all diffusion metrics and age (1933 x 1933 correlations), age-fornix associations were the strongest (**Fig.7, SF12**). Overall, the significant N = 1823 correlations (at *pHolm*<.001) ranged from |*r*|= 0.024 to |*r*| = 0.578 with |*r*|Mean = 0.245, |*r*|SD=0.122.

### Age Trajectories of Diffusion Features

In **Fig.8** we present absolute diffusion metrics for the whole brain (**Fig.8a**) and fornix (**Fig.8b**) across ages for the examined six diffusion approaches (for forceps see **SF14**; overview of metrics: **ST10**). Age-metric relationships for fornix were approximating linearity closer than more curvilinear global age-curves.

Several fornix-age relationships for BRIA extra-axonal and intra-axonal radial and axonal diffusivity opposed age relationships of whole-brain-averages, whereas forceps-age relationships closely resembled these whole-brain-average metrics’ age relationships.

Whole-brain (**Fig.9**), fornix (**SF9**), and forceps (**SF15**) diffusion metrics M were predicted from age, sex and scanner site to create age curves (**Fig.9A-B**) which can be compared to crude curves (**Fig.9C-D).** Highest SE, R^2^_adj_ and variability across metrics was observed when predicting BRIA metrics (R^2^_adj_ = .21), as well as lowest R^2^_adj_ ≈ 0 in BRIA Vextra, respectively. While DTI metrics could also be predicted well from the model, lowest variability in R^2^_adj_ was found in WMTI and DKI. For fornix metrics, SE and R^2^_adj_ was generally higher across diffusion approaches (**SF9**).

Likelihood Ratio tests indicated age dependence across global metrics (*p*_Holm_<.001), with the exception of WMTI axEAD (χ^2^=6.66, *p*_Holm_=.084; **ST11**), wheras all fornix (**ST4**) and forceps (**ST17**) features were age sensitive. While the regression lines show a slight curvature, model fit did not differ between linear and non-linear models for whole-brain (**ST12**), fornix metrics (**ST9**), and forceps minor metrics (**ST18**), indicating steady WM degeneration in mid-life to older ages.

### Associations between BAG and WM

Finally, principal components of regional and whole-brain WM metrics for each of the eight models (**Table 2**) were only weakly correlated with uncorrected BAG_u_, and similarly related to corrected BAG_c_, chronological and predicted ages (**SF10**). Furthermore, when predicting either WM components which explain most variability (**SF10**, **ST14**) or single regional or whole-brain metrics (**SF11**) from BAG_c_ and BAG_u_ and covariates, models predicted relatively small proportions of variance, with small contributions of BAG to the model (**SF10**-**11**).

## Discussion

We revealed that both conventional DTI and advanced diffusion approaches (WMTI, DKI, BRIA, SMT, mcSMT) perform consistently on brain age predictions, as indicated previously^25^. As a novel finding, our results show strong contributions of fornix and forceps minor microstructures to brain age prediction models. Additionally, among WM features, fornix shows strongest correlations with age. This suggest that the fornix and forceps minor are key WM region of cross-sectional brain age, with fornix and whole-brain dMRI metrics’ age trajectories following similar patterns such as steepening slopes at later ages. Furthermore, WM microstructure is expected to steadily degenerate in midlife to older ages, in particular, in extra axonal space.

### Limitations

There are multiple challenges related to fornix and forceps minor as drivers of brain age estimates, particularly multicollinearity, which might bias estimates of the importance of fornix and forceps minor (gain and permutation feature importance) for brain age predictions, and second, data processing artefacts. UKB offers diffusion data acquired with the most typical two-shell-diffusion protocol. Nevertheless, the standard diffusion model^66^ based on differentiation of intra- and extra- axonal water pools could not be solved using this measurement strategy^66^. As a result, the derived diffusion metrics have both numerical uncertainties and the variability introduced from non- biological parameters^66^. Quantitative metrics derived from the different diffusion approaches allow to investigate such non-biological variability and to grade the subject variability in terms of used covariances. Yet, the aforementioned technical limitation might play a decisive role in a clinical context^50, 66^.

Besides obstacles resulting from modelling assumptions, our sample is cross-sectional in design and limited to adults older than forty, which, in turn, influences predictions^45^. Additionally, the UKB imaging sub-sample shows better health than the non-imaging UKB subjects^67^. Another open question is the exact interpretation of BAG and its relationship with WM metrics. This BAG-WM relationship was found to be small for principal WM components (**SF10**) and single diffusion metrics (**SF11**). Previous research indicates no relationship between the rate of change in longitudinal regional and global T_1_-weighted-feature-retrieved BAG^18^. Yet, further investigation of longitudinal, in particular voxel-wise WM-derived BAG provides additional avenues to increase the interpretability of BAG.

Diffusion metrics were highly correlated within fornix (**Fig.6**) and forceps (**SF13**) across diffusion approaches, and show similar age trajectories (fornix: **SF9**, forceps: **SF15**). This provokes the question of redundancy of some of the metrics. The identification of redundant metrics and the combination of metrics across diffusion approaches is a matter of future research comparing diffusion approaches by probing them in practical settings such as in clinical samples^70^.

Only few studies^56, 57^ address the fornix across ages. A possible reason is fornix’ artefact- susceptibility induced from its proximity to the cerebrospinal-fluid, while being a small tubular region. Recent processing pipelines such as TBSS minimise such artefacts^80^. Yet, the influence of cerebrospinal-fluid artefacts in small tubular structures like the fornix remains unclear^68^. Fornix is a relatively small anatomical structure, and, for example, fornix BRIA cerebrospinal-fluid fraction is higher (vCSF>0.5) than global measures (vCSF>0.075), suggesting a presence of strong partial volume effect. In order to overcome such distorting effects, voxel-wise techniques are recommended, demanding the development of novel approaches incorporating techniques such as deep learning showing better performance than traditional ML, especially on large population samples^69^.

### Consistency across diffusion approaches

Overall, the results of brain age predictions are similar across diffusion approaches, with WMTI, DTI and DKI predicting age better than SMT, mcSMT and BRIA considering model fit and prediction-outcome correlations (**Table 2**). This finding could be explained in terms of diffusion approaches; i.e., the attempt to introduce more biophysically accurate parameters into the model might simultaneously reduce the general sensitivity of the used approaches to tissue changes.

Integrative approaches such as DTI or DKI are able to localise brain changes, however, without providing information about the underlying mechanisms. Our study supports a previous study with a smaller but more age-differentiated sample (*n=*702) of DTI and WMTI being superior to mcSMT at brain age predictions in terms of model performance^25^. When examining additional diffusion models on a larger sample, and also including JHU ROIs in addition to tract and whole-brain average scores, we find DKI metrics to have higher predictive power than in Beck and colleagues^25^. This effect might be partly due to added spatial detail from the added RIOs and their relationships to the tracts. Simultaneously, differences between diffusion approaches, and both variance explained and prediction error (RMSE, MAE) were smaller in this study. These differences are likely due to the narrower age range in our study^45^, whereas our significantly larger sample emphasises the reliability of our findings.

While brain age predictions from single diffusion approaches were grossly similar, predictions from combined approaches were most accurate (**Table 2**). Correlations between predicted and chronological age were consistent across diffusion approaches, as differences between correlations were small (**Fig.4**, **SF8**). This shows that addressing a wider range of WM characteristics improves predictive models compared to models with single diffusion approach metrics (e.g., only DTI), which would be intuitive when considering BAG as a general indicator of health^10, 13, 19, 20^.Vice versa, reducing spatial specificity by averaging diffusion metrics across all WM reduced prediction accuracy. Conventionally used DTI on its own is limited in its ability to present biophysically meaningful measures of the underlying microstructure. As a result, the advanced modelling is recalled including intra- and extra-axonal spaces and tissue peculiarities being influenced by individual differences in myelin and fibre architecture (crossing/bending fibres, and axonal characteristics)^25^. Hence, adding additional information to DTI better allow to infer the underlying neurobiology of tissue, for example, expressed in differential WM-age-dependences (**Fig.8-9, SF14-15**) or brain age predictions (**Table 2**)^25^.

We observed that BAG exhibits strong correlations across all diffusion approaches (**Fig.5, SF1**). Congruently with the correlational differences (**Fig.4, SF8**), BAG based on averaged skeleton values was least correlated to all other diffusion approaches (**Fig.5**), indicating inferiority of global compared to region-wide approaches. BAG obtained from WMTI, DTI and DKI were closest related to BAG from the multimodal approach (which predicted age best), both for age-bias corrected and uncorrected BAG (**Fig.5, SF1**). This is in agreement with the observed age-prediction model performance (**Table 2**). BAG correlations were observed in three clusters: 1) WMTI and DTI, 2) mcSMT, SMT, BRIA, and 3) DKI, indicative of similar measurements within these clusters (**Fig.5, SF1**). To a certain extent, these clusters reflect similarities in the underlying mathematics of the clustering diffusion approaches. For example, mcSMT and SMT are closely related models^37^, whereas DKI’s non-Gaussianity might reveal another quality of age-sensitive WM microstructures not captured by the other approaches^46^. Additionally, the cluster differences indicate that the observed diffusion approaches measure different age(ing)-sensitive characteristics, supporting the argument for a combination of diffusion approaches when assessing the ageing brain.

**Fig. 5:**
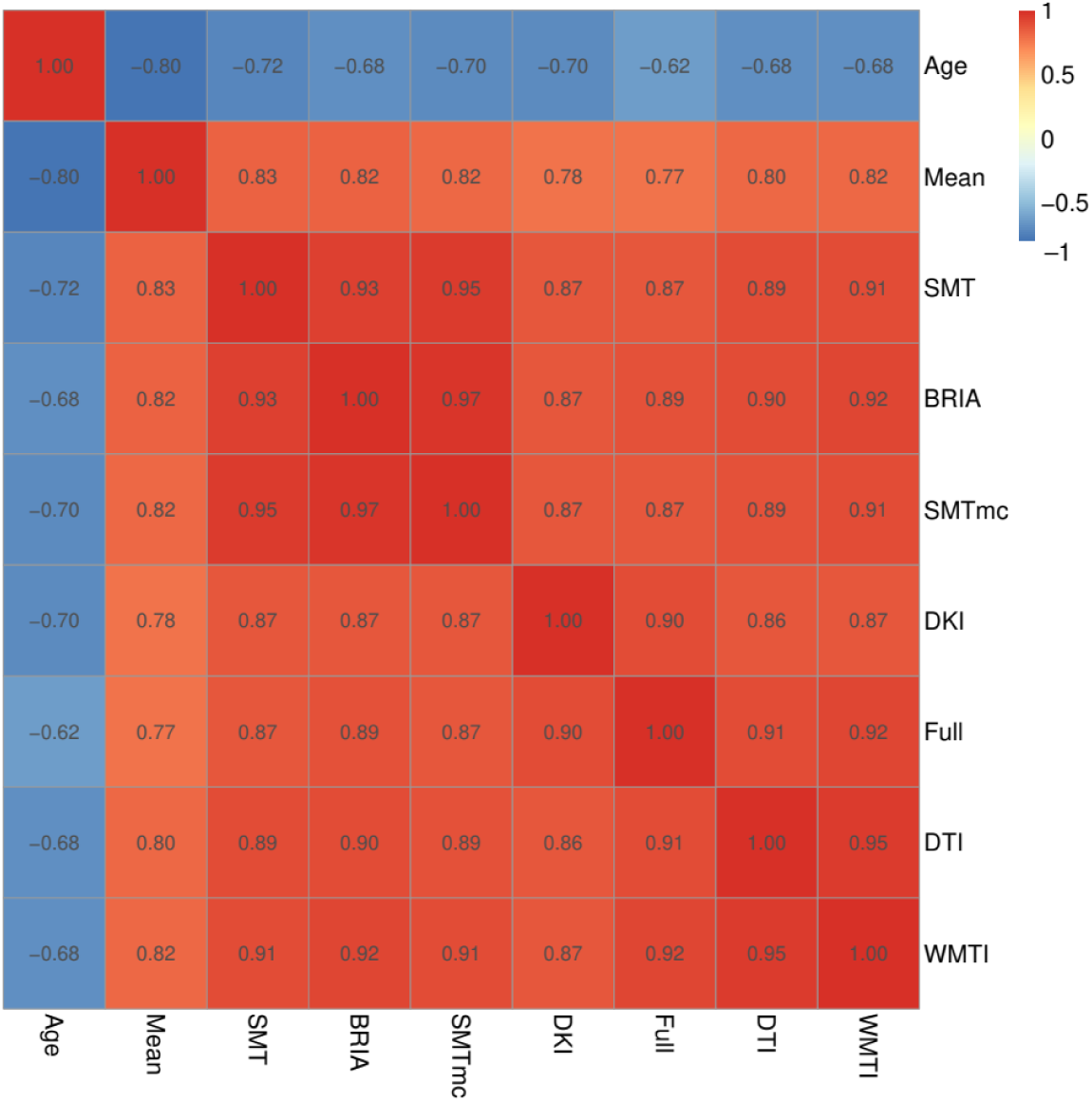
Correlations of uncorrected BAG and age across used diffusion approaches. Age-BAG correlations were significant at *pHolm* < .001. For the corrected BAG correlations across models see SF1.

### Age trajectories and fornix and forceps minor as a brain age feature

Based on the presented findings on fornix, we further investigate details of fornix, keeping discussed limitations to the generalizability of the findings in mind. Diffusion metrics describing fornix microstructure were consistently related to each other and age across all diffusion approaches in two clusters. Values were positively correlated within each cluster and negatively between clusters (see **Fig.6**). In the first cluster, different approaches’ FA, kurtosis metrics (MK, RK, AK), water fractions (vintra and vextra from BRIA and AWF from WMTI), and BRIA intra-axonal and extra-axonal radial and axial diffusivity were positively correlated. The second cluster, which was negatively related to the first cluster but positive to age, contained metrics of mean, axial and radial diffusivity, and cerebrospinal-fluid fraction of the different diffusion approaches, which were positively related to each other. Interestingly, both clusters consisted of unit-less values, for example water fractions, and diffusivities, which might have the same meaning as extra-axonal axial diffusivities from different diffusion approaches, for example BRIA vs SMTmc. Such consistencies of similar metrics across diffusion approaches were more apparent for the fornix when QC- identified outliers were removed (compare **Fig.6** and **SF4**), which supports the reliability of our findings of fornix-age-dependencies. Furthermore, fornix metrics were most strongly related to age across diffusion approaches (**Fig.7, SF11**), supporting the importance of fornix in reducing error of brain age predictions (**Table 3**). Correlations of diffusion metrics within the forceps minor were not as strong and consistent as in the fornix, and partly in the opposite direction as for the fornix (**SF13**). Not surprisingly, all fornix and forceps minor features were age-sensitive (**ST4, ST17**), and more age sensitive than whole-brain metrics (compare: **ST11**). Whole-brain trajectories are in agreement with previous results, showing-age sensitivity of various mean diffusion metrics^25^, and the same directionality of age trajectories of metrics for DTI^9, 33^, mcSMT, DKI, WMTI^25^.

**Fig. 6:**
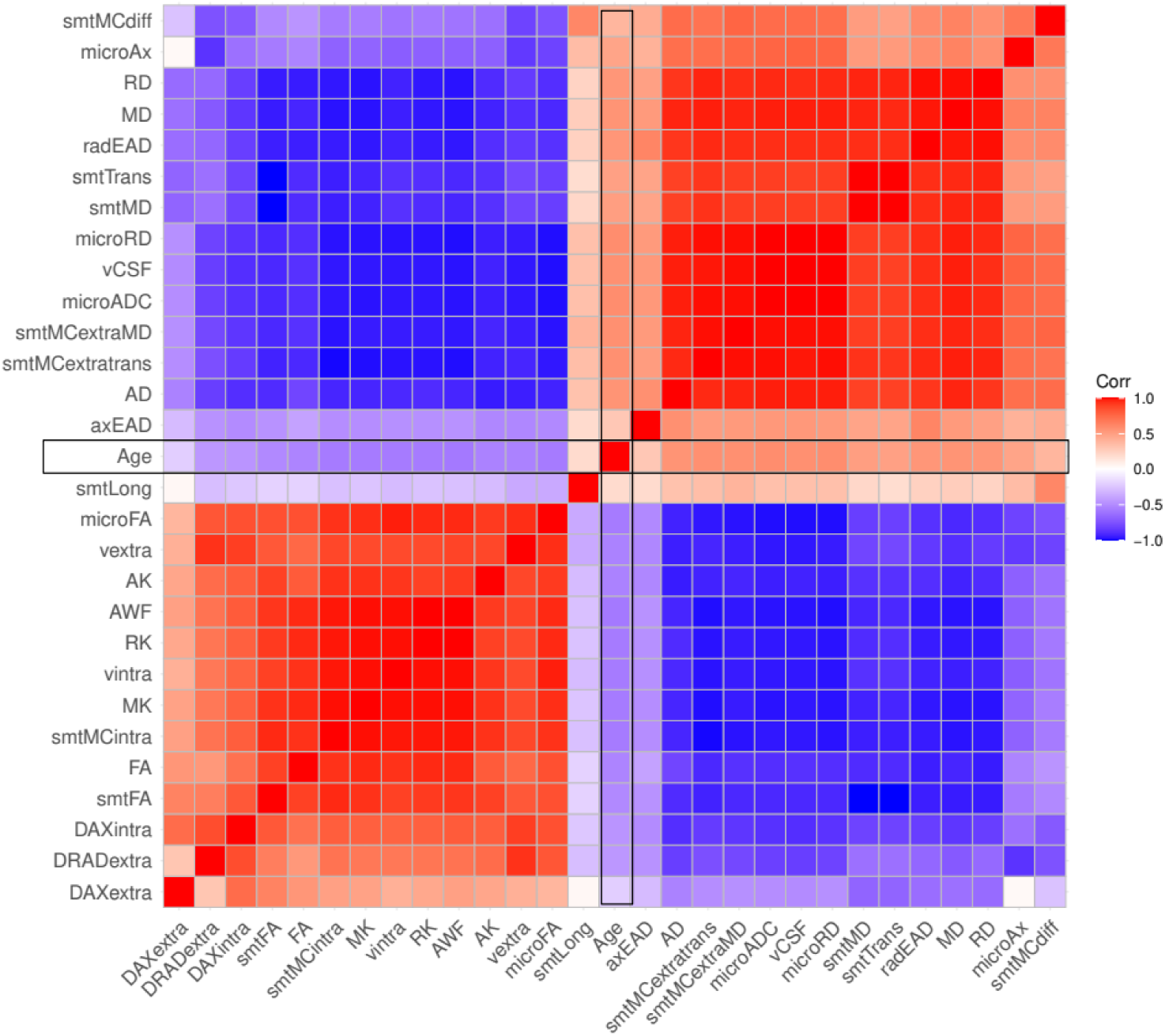
Correlation matrix for fornix diffusion metrics and chronological age. All correlations were significant at Holm-corrected *pHolm* < .05.

**Fig. 7:**
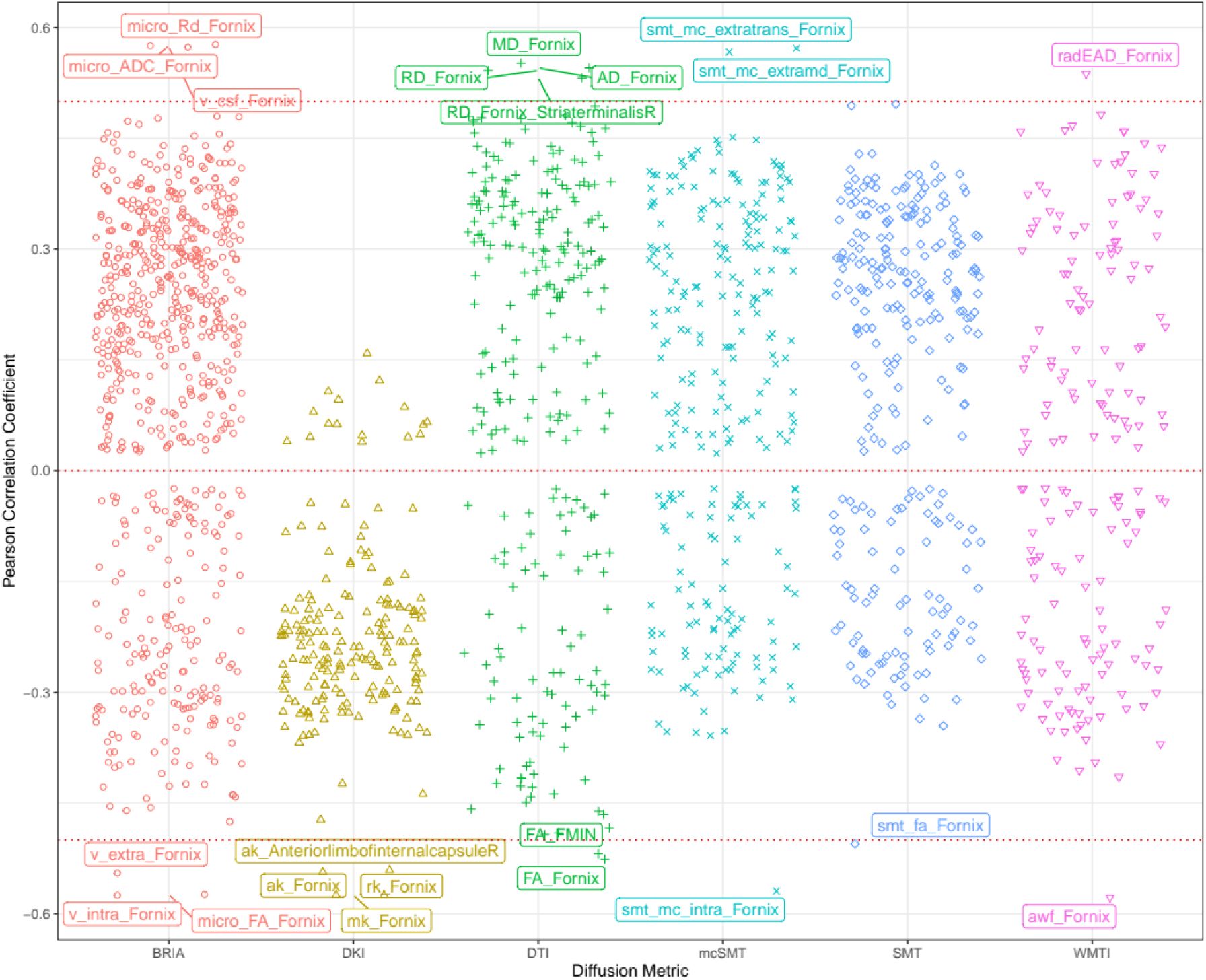
Correlations between diffusion metrics and age. *Note*: Each point indicates one correlation between a diffusion metric and chronological age. Names of diffusion metrics are displayed when correlations between the metric and age reached a Pearson correlation of |*r|*>0.5. Holm correction^44^ was used for Holm-correction, and all displayed values were significant at *p* < .001. For the distribution of the correlations see **SF12**.

We displayed differential behaviours of fornix microstructure measures across diffusion approaches (**Figs.8-9**). Focussing on absolute diffusion values (**Fig.8**), diffusion measures which are correlated (**Figs.5-6, SF13**) exhibit similar age dependences. Additionally, slopes of fornix compared to whole-brain diffusion metrics were generally steeper and closer approximating linearity, indicating stronger changes, such as quicker WM degeneration in the fornix compared to the whole-brain average (see **Fig.8**). Particularly BRIA metrics show visually detectable differences between the fornix and the whole brain (**Fig.8**, DAXextra, DAXintra, DRADextra, Vextra); as opposed to global age trends which are also strongly resembled by forceps minor (**SF14**), fornix intra and extra-axonal diffusion decreased, indicating fornix shrinkage with increasing age. Periventricular shrinkage is linked to enlarging ventricles^47^, which has been related to ageing and neurodegenerative disorder progression^48^. This effect was observed by a positive relationship between age and cerebrospinal fluid (CSF) fraction in BRIA. Another metric which revealed larger differences in the fornix than for the whole-brain average was intra-axonal water fractions, which can be treated as a proxy for the axonal density, decreased with increasing age (see **Fig.8**, BRIA:Vintra; SMTmc:intra; WMTI:AWF) while the CSF fraction (BRIA) increases. Such WM microstructure changes are not only directly linked to different neurobiological features but can be markers of clinical outcomes, such as dementia^49, 50^.

**Fig. 8:**
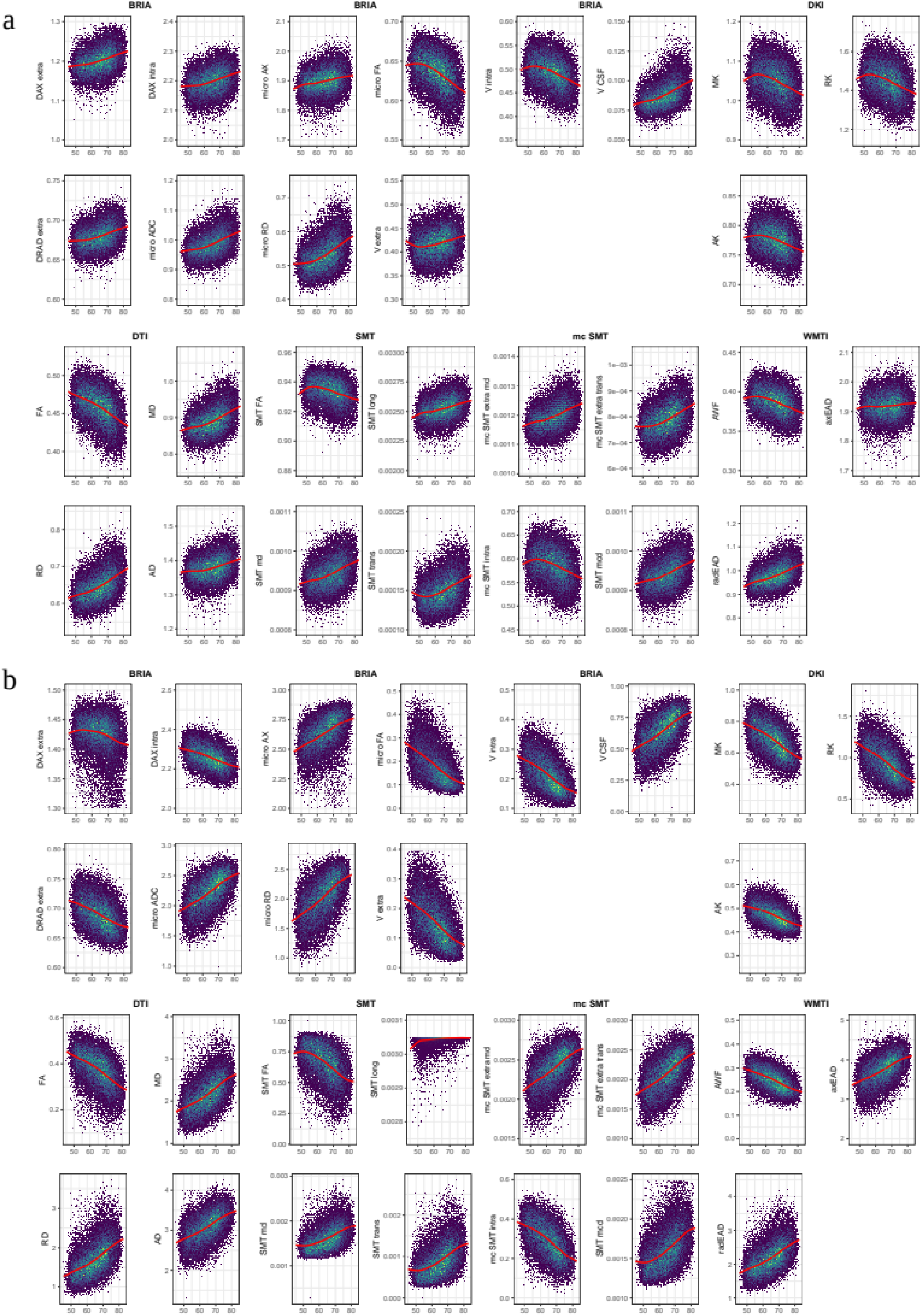
Whole-brain and fornix diffusion metrics across age. *Note*: The presented plots represent diffusion metrics for each of the six diffusion models from the full sample *N* = 35,749 for a) whole-brain diffusion metrics, b) fornix diffusion metrics. Brighter colours indicate higher density and red lines are fitted lines to the relationship between age and diffusion metric. Plots for forceps can be found in SF14.

A selection of metrics is comparable across diffusion approaches when taking DTI as reference point and focussing on similar age trends. DTI metrics AD, RD, and MD tend to increase over the lifespan and FA tends to decrease across brain regions (**Fig.8-9**)^25, 33, 51, 52^ as well as in fornix (**Fig.8b**, **SF9**), implying processes such as de-myelination, changes in axonal and general WM integrity.

Such DTI-age-dependences are reflected by according BRIA, SMT, and WMTI metrics, whereas DKI shows opposite age-relationships, as presented previously^25^. Deterioration effects, measured by the age-dependency of axonal water fractions, were generally stronger in fornix compared to whole- brain metrics (**Fig.8**). Interestingly, opposed to global metrics, radial diffusivity measures from DKI and BRIA (DRADextra) decreased in fornix (**Fig.8**), suggesting higher fornix than global plasticity, potentially being an antecedent of age-related hippocampal changes^55^.

Additional, unique information about age dynamics was presented by standardised scores corrected for age, sex and scanner site and crude standardised scores across ages (**Fig.9**, **SF9**). After corrections, most fornix metrics follow a tightly resembling near-linear trend either increasing or decreasing by age (**SF9A-B**), as opposed to forceps minor (**SF15**) and whole-brain metrics which follow a rather curvilinear line, as previously shown^25, 33, 52^. Diffusion metrics’ variance explained across models indicates fornix metrics to be more sensitive to a combination of covariates age, sex, and scanner site than whole-brain metrics (**Fig.9, SF9**). In the fornix, only BRIA extra-axonal axial diffusivity (DAX extra) and the SMT longitudinal diffusion coefficient (SMT long) showed non- linear trajectories, however, both measures are weakly correlated to other diffusion parameters (**Fig.9**). Yet, when comparing model metrics such as variance explained of linear and non-linear models predicting fornix, forceps minor, and whole-brain diffusion metrics from age, sex and scanner site and their interactions, there were no apparent differences between models (**ST9, ST12, ST15**). This implies that contrary to previous research observing the entire lifespan presenting curvilinear DTI age trajectories^25, 33^, or trends towards curvilinearity (with yet better linear fit for selected regions)^52^, we found that fornix and whole-brain age trajectories from age 40 can be described as linear when accounting for covariates sex, age, and scanner site. While the crossing of the x-axis at age 65 (**Fig.9**, **SF9, SF15**) is a reflection of the sample’s age distribution (**Fig.1**), in addition to the shapes of the different age-trajectories, it reveals that the different diffusion approaches are similarly age-sensitive or measure similar underlying ageing-related changes. For whole-brain metrics, changes become exacerbated from 65 onwards (**Fig.1**), with reasons potentially laying in an accelerated neurodegeneration also reflected in the exponentially increasing risk to develop neurodegenerative disorders from age 65 onwards^53^. For example, in the USA, 3% of 65-74 year olds, 17% of the 75-84 year olds, and 32% of those ag 85+ developed Alzheimer’s dementia^54^. Subclinical or preclinical states are, however, not captured by these approximations, and WM changes usually precede clinical detections. This makes WM monitoring a promising tool for early neurodegenerative disease detection.

**Fig. 9:**
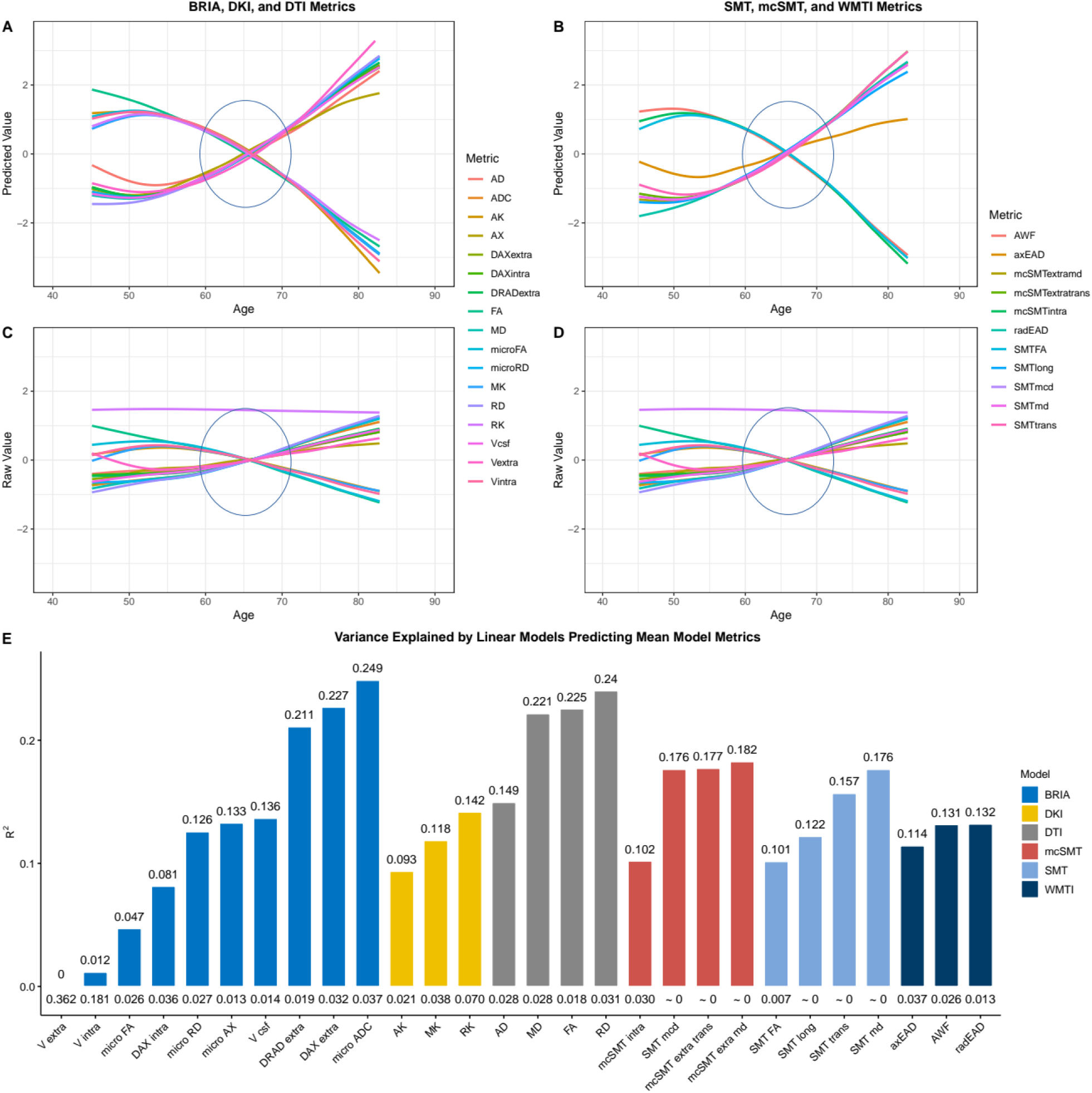
Raw and predicted whole-brain WM diffusion metrics by chronological age. Fig.9A-D shows age curves for each standardised (z-score) diffusion metric’s mean skeleton value (y-axis) plotted as a function of age (x-axis). Shaded areas represent 95% CI. Curves fitted to raw values (Fig.9 C-D) serve as a comparison to the lm-derived predicted values from Equation 1 (Fig.A-B). Fig.9E indicates the model fit for the linear models from Fig.9A-B, showing R^2^adj values on top and Standard Error (SE) on the bottom of the bars which each represent a Fornix skeleton value for one of the seven models. Lines crossing at age 65 are marked with ovals. Model summaries of all 28 mean models can be found in ST5. The same visualisation of fornix diffusion values can be found in SF9, and for the forceps minor in SF15.

Beyond WM, fornix changes seem to play an important role for GM changes, particularly in the hippocampus: for example, fornix glia damages lead to hippocampal GM atrophy^55^. This might be reflected by dis-connectivity of fornix with other brain regions as described by decreasing extra axonal space coefficients (**Fig.8b**), and following changes in fornix function. Potentially, the consequences of age-related fornix changes thereby affect functionality of a selection of brain regions, such as the hippocampus. While several studies have presented ageing-related fornix microstructure changes in humans^56, 57^ and monkeys^58^ in small samples, only one large-scale study revealed findings connected to the fornix, namely strongest default mode network GM volume covariation with fornix WM microstructure^59^. This suggests that fornix, a key connector of the limbic system with the cortex, might also be critical for default mode network functioning.

Moreover, memory and episodic recall have been related to fornix^60^. Hence, fornix changes might play an important role in known ageing-dependent temporal lobe changes, and specifically hippocampal changes for ageing-related pathological developments^61–64^. Previous studies presented age-related fornix DTI metric changes^55–57^ which potentially appear prior to hippocampal volume changes^55, 56^, and are related to declining episodic memory performance^55^. Hence, fornix changes potentially serve to predict future pathological development, suggesting fornix WM microstructure and changes in such as ageing biomarkers. This supports previous findings showing network re- activations, metabolic and GM changes after fornix deep-brain-stimulation, antagonising the progression of neurodegenerative disorders^65^.

Different studies showed age-related deterioation effects in the forceps minor^95–96^, a sub-region of the corpus callosum. Loss in WM integretiy have also been associated with various phenotypes, for example, behavioural impacts, such as mental slowing^97^, and various disorders, such as major depressive disorder^98^, schizophrenia^99^, dependencies on cocaine^100^ and alcohol^101^, with WM degeneracy explaining higher impulsivity in cocaine addiction^100^. Overall, the forceps are assumed to have an important role of connecting both hemispheres, which might be crucial for interhemispheric signal propagation^102^. Previous research shows also that WM changes in FA and MD relate to GM thinning with the forceps being particularly vulnerable to such changes^105^.

Moreover, cognitive test scores were related to forceps minor AD and MD scores in Alzheimer’s Disease patients^103^, and already at mild cognitive impaired forceps minor FA and MD scores were different from age-matched participants with subjective cognitive decline^104^. FA was also shown in this study as important brain age feature for both multimodal and DTI models (**Table 3**). This suggests forceps as an important region for brain age and ageing.

The current study gives for the first time a detailed account on region-wise-to-global WM-age relationships for multiple diffusion approaches in a representative sample, and highlights fornix and forceps minor as an important structures for age predictions across diffusion approaches. Brain age was estimated best when combining diffusion approaches, showing different aspects of WM to contribute to brain age with fornix and forceps minor being the central regions for these predictions.

## Data Availability

All raw data are available from the UKB^5^ (www.ukbiobank.ac.uk). Synthetic datasets with the synthpop^89^ R package based on the original data for all six diffusion approaches (resulting in six datasets) to run the code are openly available at the Open Science Framework: (https://osf.io/nv8ea/). Synthetic datasets are simulated datasets closely mimicking the statistical characteristics of the original data while protecting data privacy and anonymity.

## Code Availability

Code needed to run brain age predictions in Python, and for all analyses and visualisations in R is available at the Open Science Framework: (https://osf.io/nv8ea/).

## Acknowledgements

This research was funded by the Research Council of Norway (#223273). This study has been conducted using UKB data under Application 27412. UKB has received ethics approval from the National Health Service National Research Ethics Service (ref 11/NW/0382). The work was performed on the Service for Sensitive Data (TSD) platform, owned by the University of Oslo, operated and developed by the TSD service group at the University of Oslo IT-Department (USIT). Computations were performed using resources provided by UNINETT Sigma2 – the National Infrastructure for High Performance Computing and Data Storage in Norway. Finally, we want to thank all UKB participants and facilitators who made this research possible.

## Author contributions

M.K.: Study design, Software, Formal analysis, Visualisations, Project administration, Writing – original draft, Writing – review & editing

A.M.d.L.: Software, Writing – review & editing

D.v.d.M.: Software, Writing – review & editing

A.L.: Writing – review & editing, Funding acquisition

E.E.: Writing – review & editing

D.B.: Writing – review & editing

O.A.A.: Writing – review & editing, Funding acquisition

L.W.: Writing – review & editing, Funding acquisition

I.I.M. supervision, Study design, Data pre-processing and quality control, Writing – review & editing, Funding acquisition

## Conflicts of Interesting

The authors have no conflicts of interest to disclaim.

## Supplement

### Supplementary Figures

**SF1:**
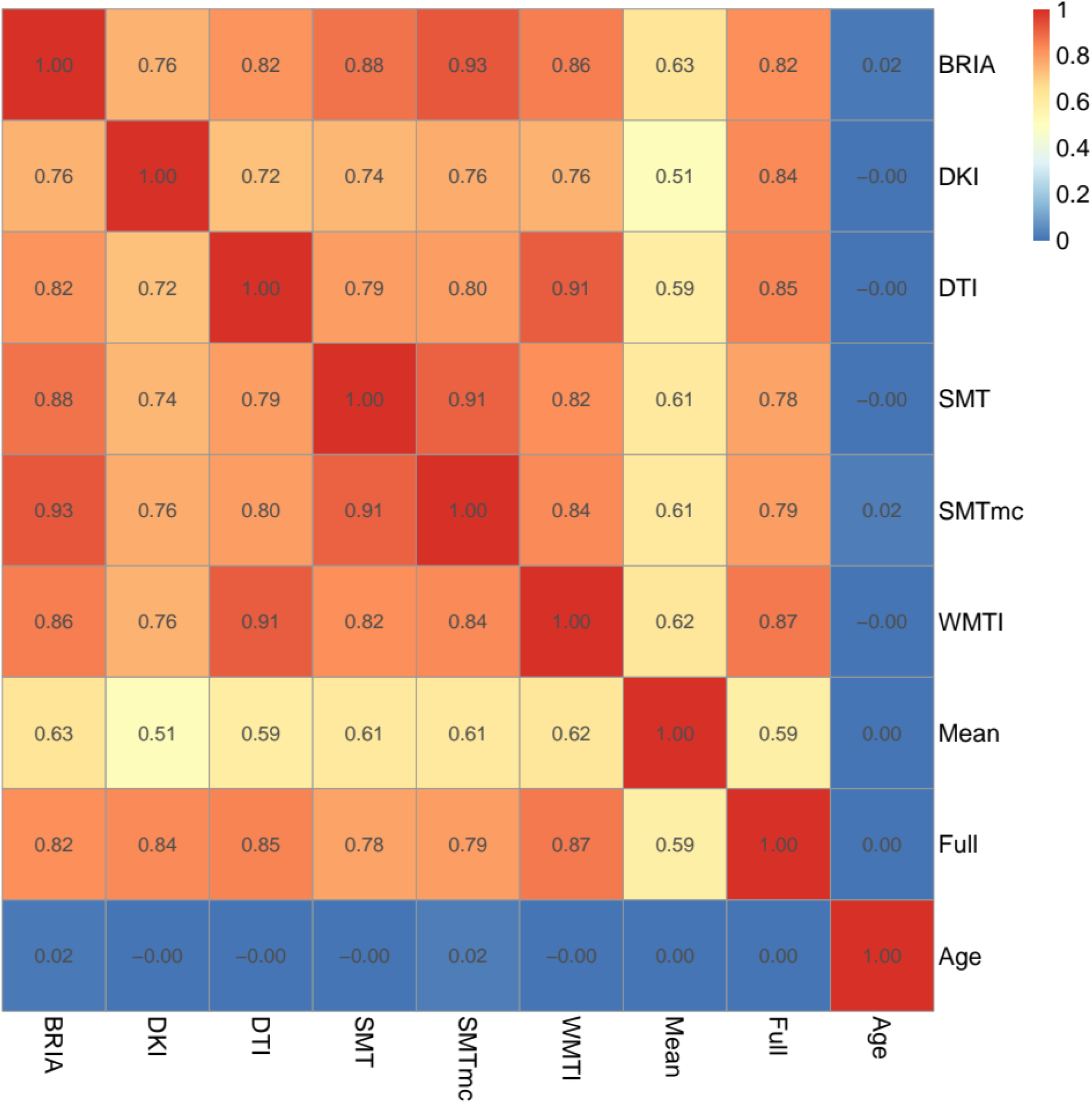
Correlations of corrected BAG and age across models. Mean = multimodal model including only mean metrics; Full = full multimodal model including all diffusion indices. Age-BAG correlations, approximating 0, were not significant at *pHolm* ≥ .05. All other correlations were significant at *pHolm* < .001.

**SF2:**
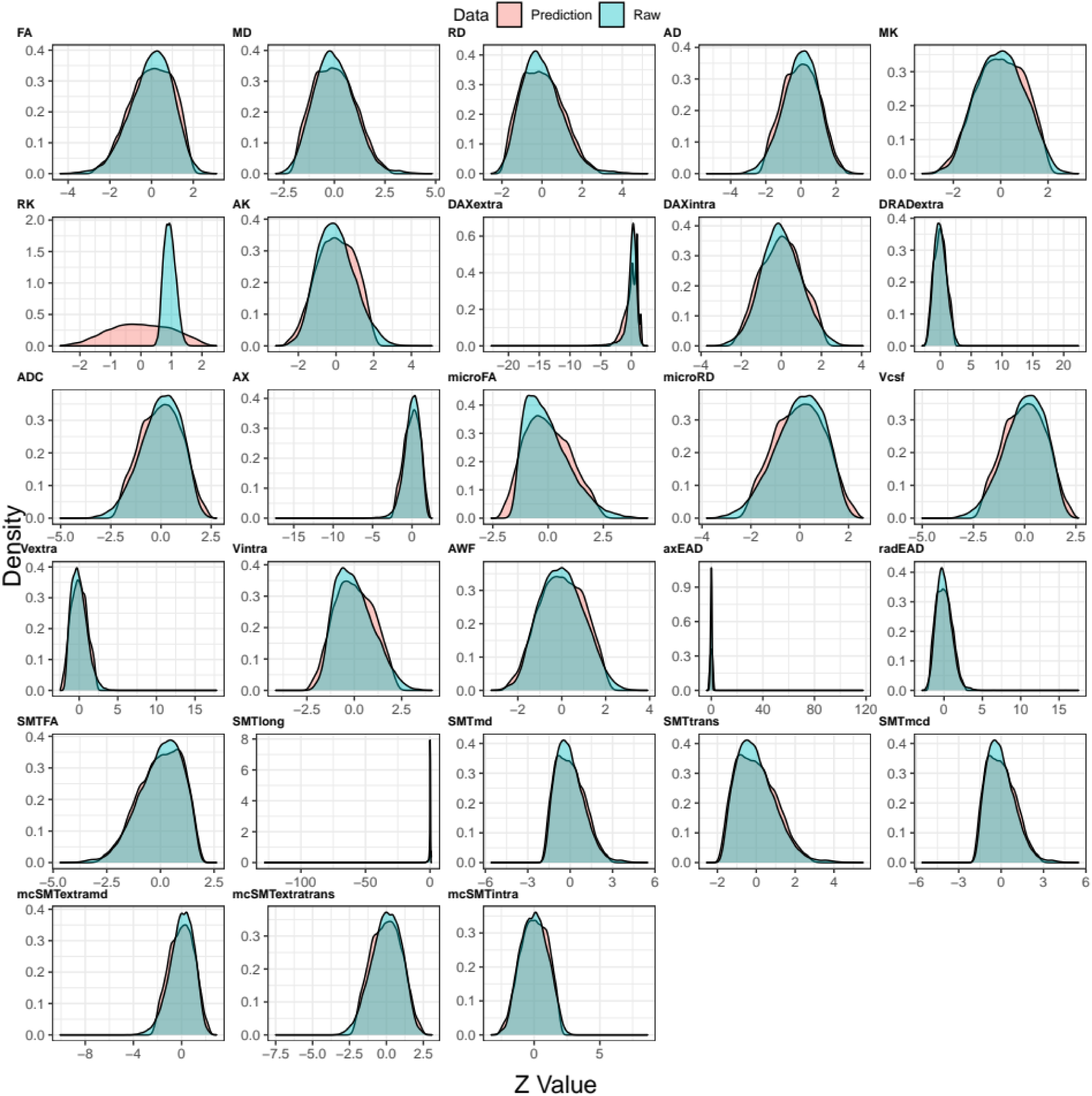
Comparison of predicted and raw fornix Z-scored diffusion metrics’ density. Density plots for each Z-scored (standardised) raw and predicted values for each fornix metric from the six observed diffusion models. Predictions were made from the linear model described in Equation 1. Find the same density plot for data including QC outliers in SF3.

Supplementing the density plots, two one-sided tests for equivalence testing (TOST)^87, 88^ were used to test whether mean differences between the model’s predictions (**SF9A-B**) and the raw scores (**SF9C-D**) are equal to zero with the assumptions that observed *Z*-score differences smaller |0.5| are equal to 0. Following this assumption, differences were equal to zero for all metrics, except the DKI metric RK: M_diff_ = 0.943, 95% CI [0.935, 0.951], *p* ≈ 1.

**SF3:**
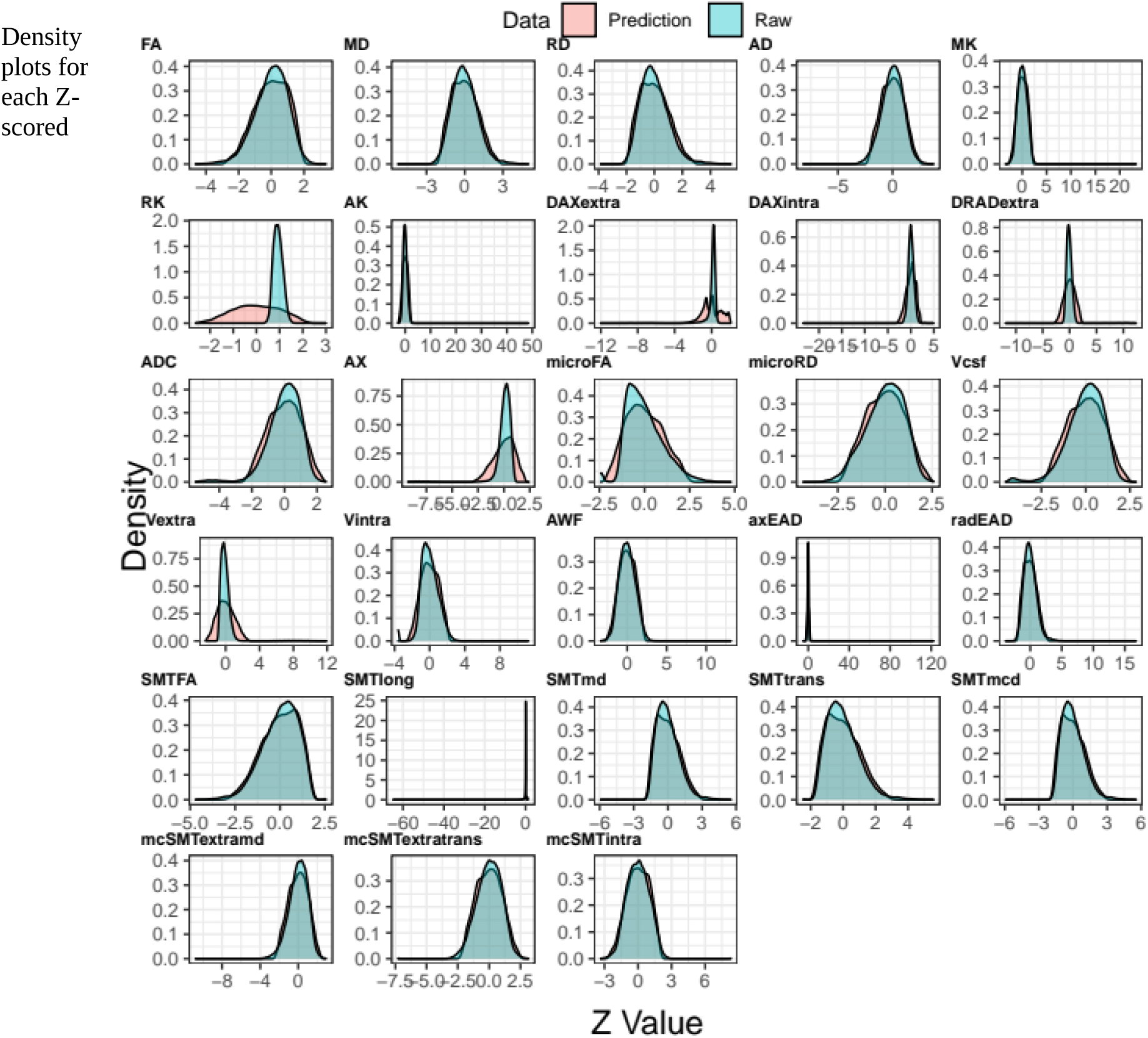
Comparison of predicted and raw Fornix Z-scored diffusion metrics’ density including QC outliers. Density plots for each Z-scored (standardised) raw and predicted values for each fornix metric from the six observed diffusion models on data *containing QC outliers*. Predictions were made from the linear model described in Equation 1. Outliers were defined by the YTTRIUM method^39^ including outlier removal based on density-based spatial clusterisation (k-means)The total data used here was Nfull+outliers = 38,687, including the full data Nfull = 35,749 used for all analyses and Noutliers = 2,938 datasets defined as outliers. This dataset does not include participants who withdrew their consent or participants with an ICD-10 diagnosis categories G or F or stroke, category I.

**SF4:**
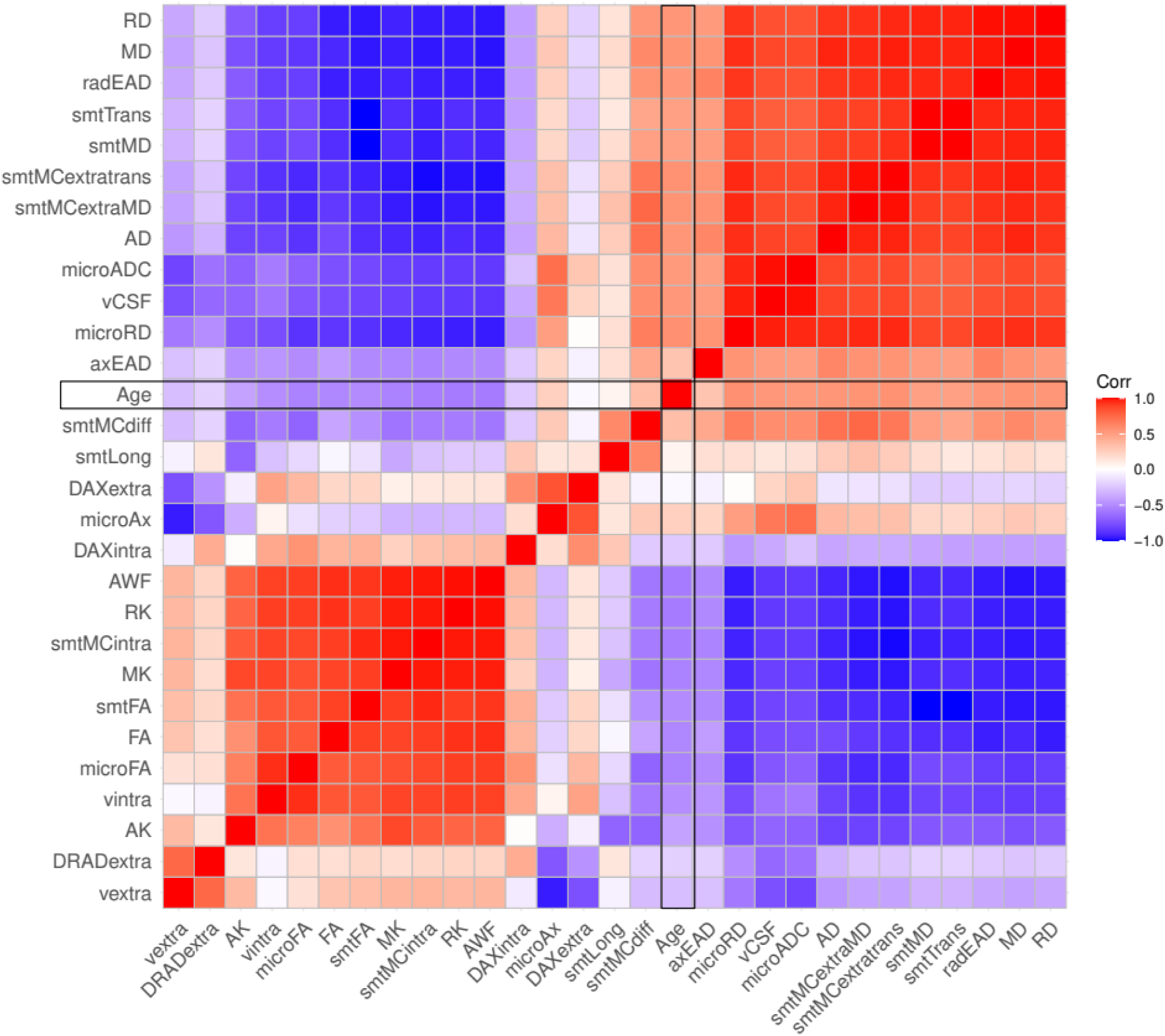
Correlations between Fornix diffusion metrics and chronological age for data including QC outliers. All correlations were significant at FWE-corrected *pHolm* < .05. Outliers were defined by the YTTRIUM method^38^ including outlier removal based on density-based spatial clusterisation (k-means). The total data used here was Nfull+outliers = 38,687, including the full data Nfull = 35,749 used for all analyses and Noutliers = 2,938 datasets defined as outliers. This dataset does not include participants who withdrew their consent or participants with an ICD-10 diagnosis categories G or F or stroke, category I.

**SF5:**
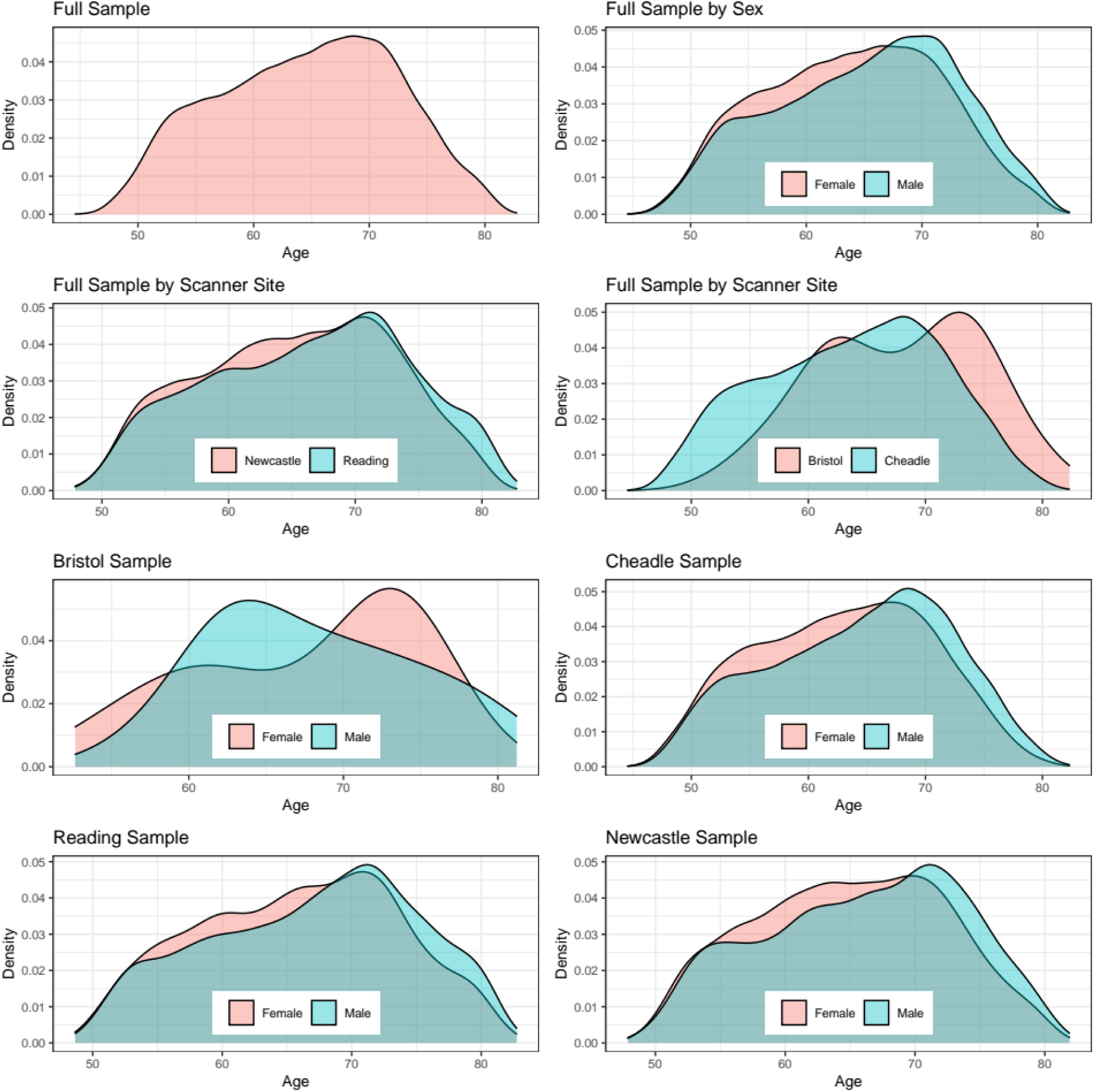
Density plots for the sample’s age by sex and scanner site for data including QC outliers. Outliers were defined by the YTTRIUM method^38^ including outlier removal based on density-based spatial clusterisation (k-means). The total data used here was Nfull+outliers = 38,687, including the full data Nfull = 35,749 used for all analyses and Noutliers = 2,938 datasets defined as outliers. This dataset does not include participants who withdrew their consent or participants with an ICD-10 diagnosis categories G or F or stroke, category I.

**SF6:**
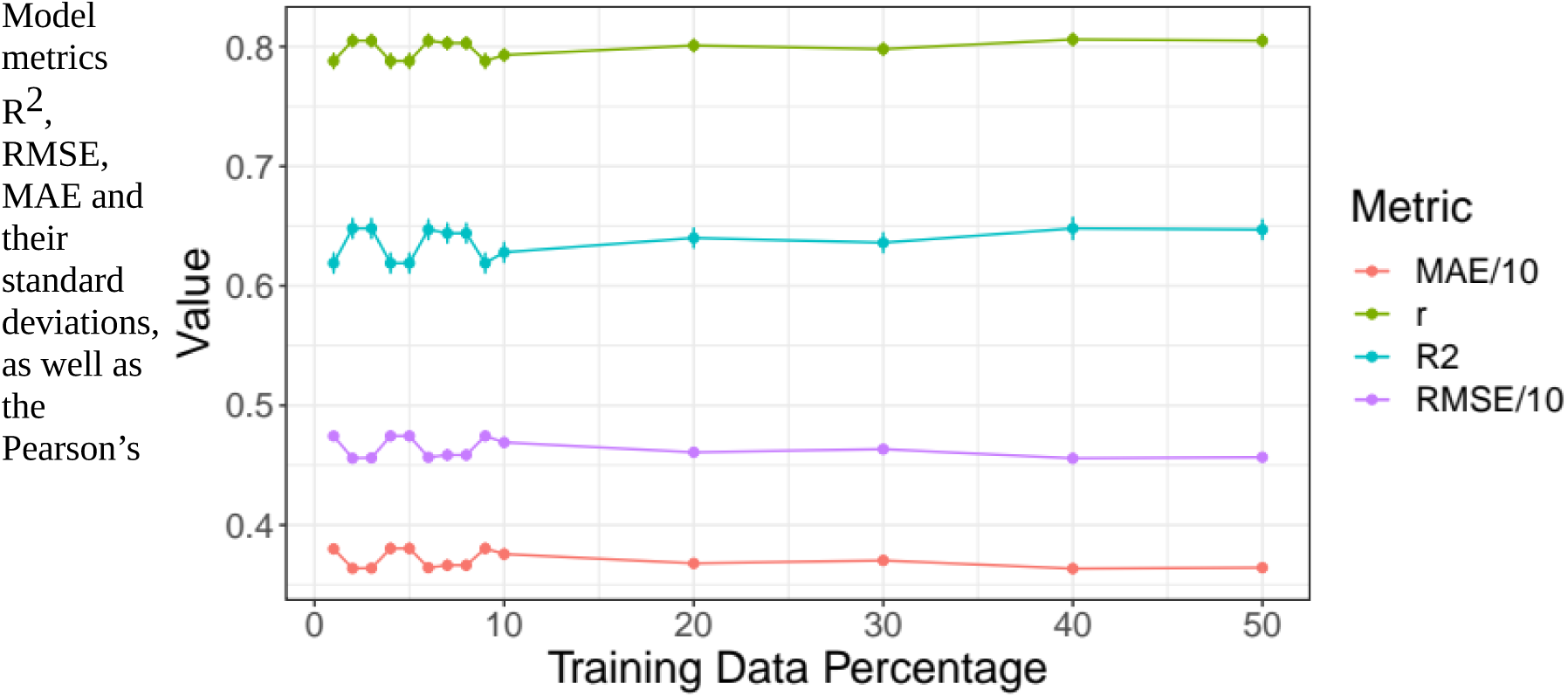
Model performance for different train-test splits for data *including QC outliers*. Model metrics R^2^, RMSE, MAE and their standard deviations, as well as the Pearson’s correlations between predicted and chronological age and its 95% confidence interval are displayed for different training data percentages of the total data (x-axis). For visualisation purposes, RMSE and MAE were divided by 10. For exact values see Suppl. Table ST8. Outliers were defined by the YTTRIUM method^38^ including outlier removal based on density-based spatial clusterisation (k-means). The total data used here was Nfull+outliers = 38,687, including the full data Nfull = 35,749 used for all analyses and Noutliers = 2,938 datasets defined as outliers. This dataset does not include participants who withdrew their consent or participants with an ICD-10 diagnosis categories G or F or stroke, category I.

**SF7:**
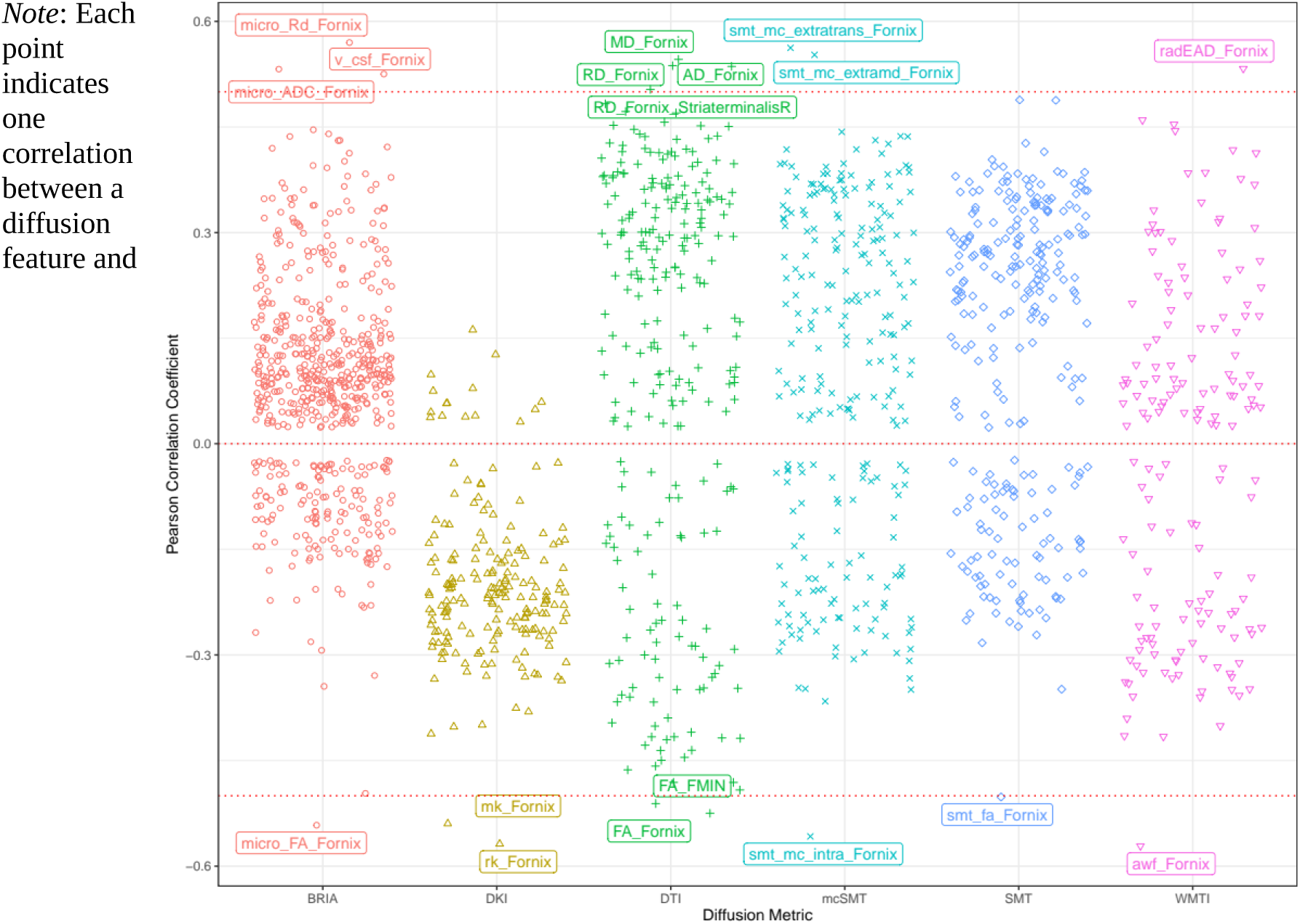
Correlations between diffusion metrics and chronological age for data including QC outliers. *Note*: Each point indicates one correlation between a diffusion feature and chronological age. Names of diffusion features are displayed when correlations between the feature and age reached a Pearson correlation of |*r|*>0.5. Holm correction was used for FDR-correction, and all displayed values were significant at *p* < .001. Results for the analysis run on data *not* including QC outliers (N = 35,749) can be found in Fig.9. Outliers were defined by the YTTRIUM method^38^ including outlier removal based on density-based spatial clusterisation (k-means). The total data used here was Nfull+outliers = 38,687, including the full data Nfull = 35,749 used for all analyses and Noutliers = 2,938 datasets defined as outliers. This dataset does not include participants who withdrew their consent or participants with an ICD-10 diagnosis categories G or F or stroke, category I.

**SF8:**
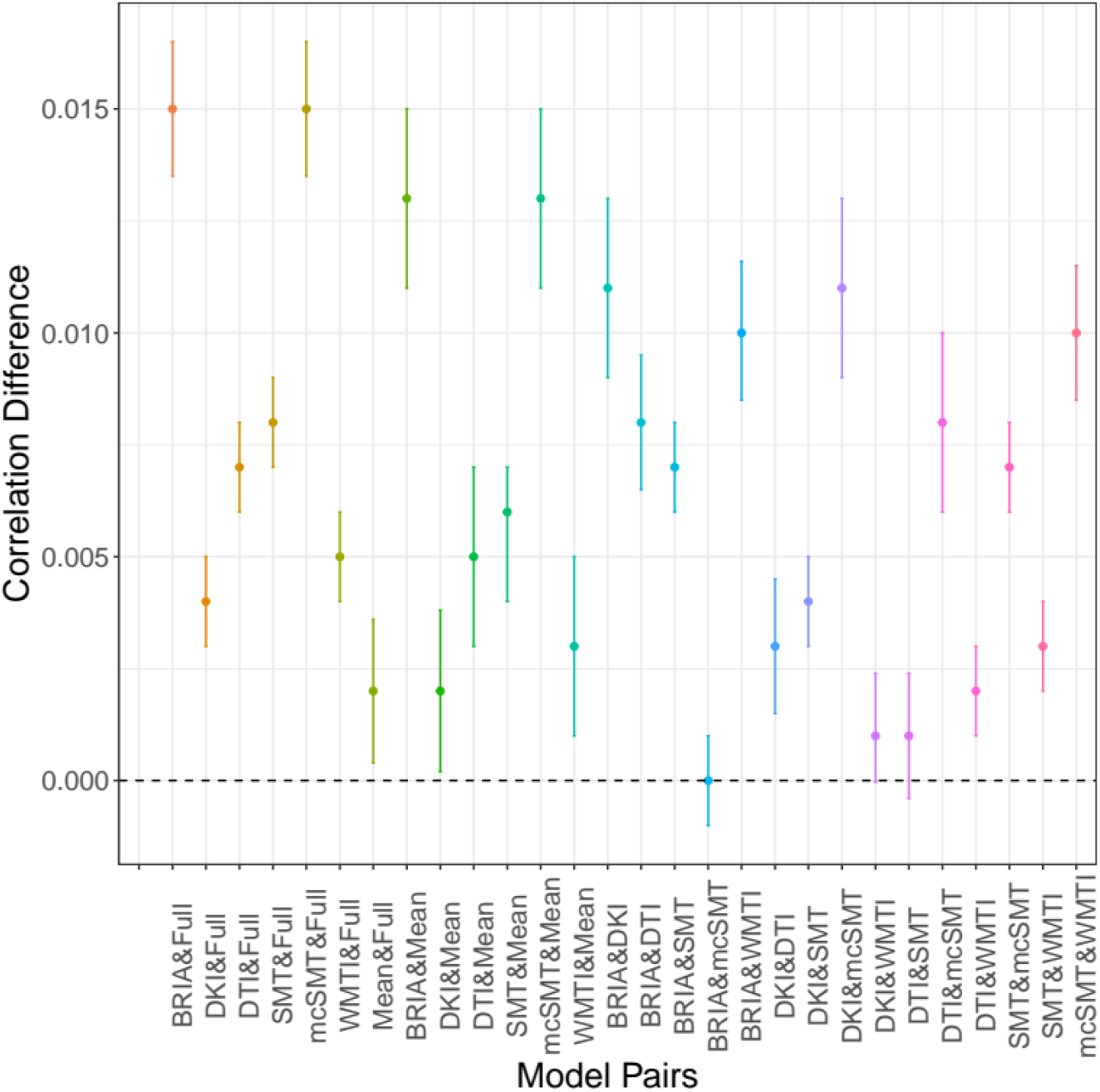
Differences between correlations of chronological and *corrected* predicted age across diffusion approaches with 95% confidence interval.

**SF9:**
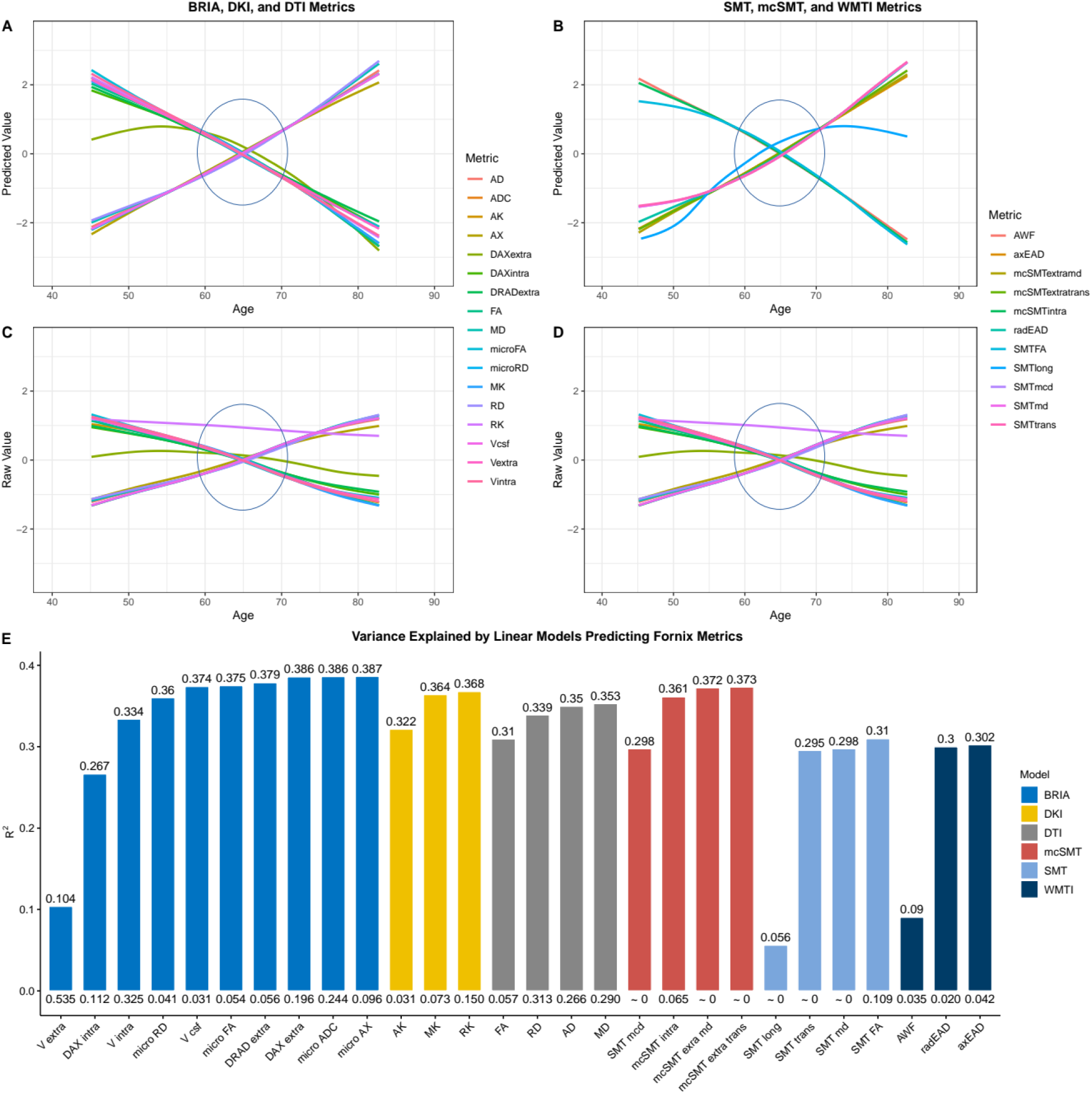
Raw and predicted fornix diffusion metrics by chronological age. SF9A-D shows age curves for each standardised (z-score) fornix diffusion skeleton value (y-axis) plotted as a function of age (x-axis). Shaded areas represent 95% CI. Curves fitted to raw values (SF9C-D) serve as a comparison to the lm-derived predicted values from Equation 1 (Fig.A-B). SF9E indicates the model fit for the linear models from SF9A-B, showing R^2^adj values on top and Standard Error (SE) on the bottom of the bars which each represent a Fornix skeleton value for one of the seven models. Lines crossing at age 65 are marked with circles. Model summaries of all 28 Fornix models can be found in ST5. The same visualisation of diffusion values averaged across the brain can be found in Fig.9. Model fit metrics R^2^adj and Standard Error (SE) for the models accounting for age, sex and scanner site (Equation 1) when predicting fornix metrics were calculated (**SF9E;** see Fig.9 for whole brain metrics). Highest R^2^adj and variability across metrics were observed when predicting BRIA fornix features, lowest R^2^adj when predicting SMT fornix metrics. DKI, DTI and mcSMT fornix diffusion metric predictions were most consistent, with BRIA, mcSMT and SMT having one outlier each, Vextra, SMTlong, and AWF, respectively, being less sensitive to age, sex and scanner site. Highest SE could be observed in the BRIA model and the lowest SE in SMT. To test age-sensitivity of the fornix features, likelihood ratio tests were conduced comparing models derived from Equation 1 against models derived from the same formula with age removed (Equation 2). All models showed significant age dependence, with BRIA microRD (χ^2^= 14,480.54, *p*Holm < .001), microADC (χ^2^= 14,384.87, *p*Holm < .001) and SMT vCSF (χ^2^= 14,311.47, *p*Holm < .001) being the most age-sensitive metrics, and mcSMT smtLong (χ^2^= 1,554.49, *p*Holm < .001), BRIA DAXextra (χ^2^= 1,824.54, *p*Holm < .001) and axEAD (χ^2^= 3,024.74, *p*Holm < .001) the least age-sensitive metrics (**ST4**).

**SF10.**
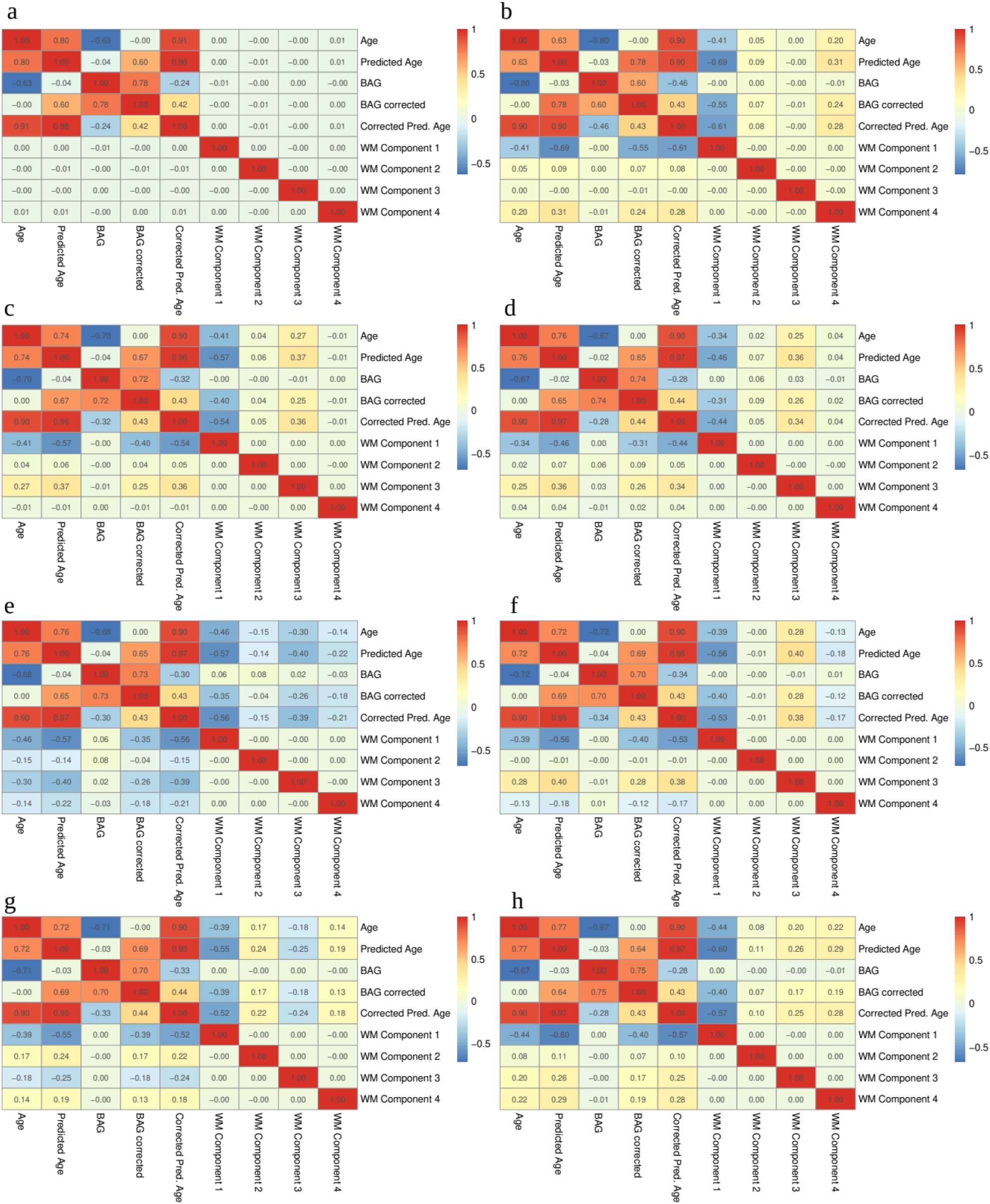
Pearson’s *r* for age, brain age and WM principal components’ relationships. All correlations with Pearson’s *r* > .01 were significant at p < .001 Read row-wise from top-left to right with matrixes indicating a) full multimodal data, b) mean/whole brain average data, c) BRIA, d) DKI, e) DTI, f) SMT, g) SMT mc, h) WMTI The first five principal components of the respective number of WM metrics for each of the eight principal components analyses were related to age, predicted (brain) age, corrected predicted age, uncorrected and corrected BAG (see **ST13** for overview of variance explained by principal components). Notably, BAG was not or only weakly related to WM components, and relationships of age, predicted age, corrected predicted age and corrected BAG with WM components followed the same pattern of direction and strength of associations, suggesting age-dependencies of these measures. When predicting the first 4 components retrieved from the respective models (as done for brain age predictions), using BAG, age, sex, site, as well as age-sex and sex-site interactions as predictors (as specified in Equation 1), different sized proportions of the variance in the components could be explained with corrected and uncorrected BAG models not differing in variance predicted and beta values. Average data BAG models explained most variance in its first component R^2^ = .505, with bBAG = -0.673, followed by WMTI R^2^ = .372, with bBAG = -0.847, and the DTI BAG model R^2^ = .358, with bBAG = -1.082. The second component was best predicted by a DTI BAG model R^2^ = .152, with bBAG = -0.059. The third component was best predicted by the DKI BAG model R^2^ = .256, bBAG = 0.170, followed by the DTI BAG model R^2^ = .250, bBAG = -0.210; and the SMT BAG model R^2^ = .247, bBAG = 0.291. Finally, the last component was best predicted by the full BAG model, R^2^ = .128, bBAG = 0.0002. For an overview of all BAG models’ performance see **ST14**. For a more nuanced follow-up analysis of global and regional individual diffusion metric predictions see **SF11**.

**SF11.**
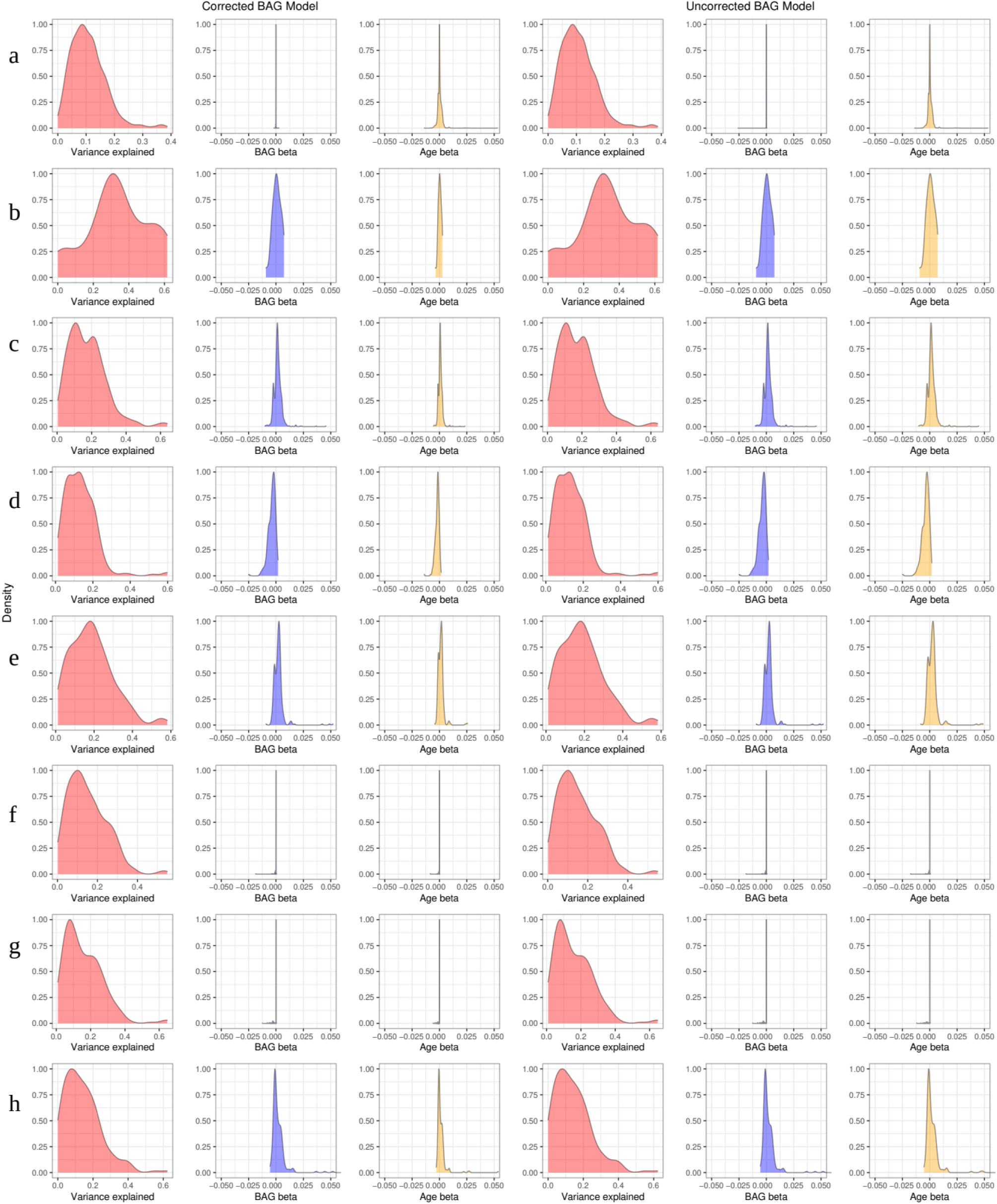
Predictions of individual global and regional diffusion metrics. Panels indicate used models: a) full multimodal model including all approaches global and local fatures, b) mean multimodal modal, including only global metrics of all diffusion approaches, c) BRIA, d) DKI, e) DTI, f) SMT, g) mcSMT, h) WMTI. We predicted the individual 1940 regional and global WM diffusion metrics from BAG, site, sex, age, as well as sex- age and sex-site interaction terms. While there were no differences in explaining variance between corrected and uncorrected BAG, models coefficients differed (see **SF11**). Variance explained across statistically significant models (at Bonferroni-corrected *p* < 0.05/1940) ranged from adjusted R^2^min = .001 to R^2^max = .387 (R^2^mean = .108, SD = 0.062), and beta values for BAG ranged from bBAG > -0.001 to bBAG< 0.001, with most variance explained in metrics Fornix v csf (Radj^2^ = .387, bBAG > -0.001, bage = 0.009), Fornix micro RD (Radj^2^ = .386, bBAG > -0.001, bage = 0.009), and Fornix micro ADC (Radj^2^ = .386, bBAG > - 0.001, bage = 0.019).

**SF12.**
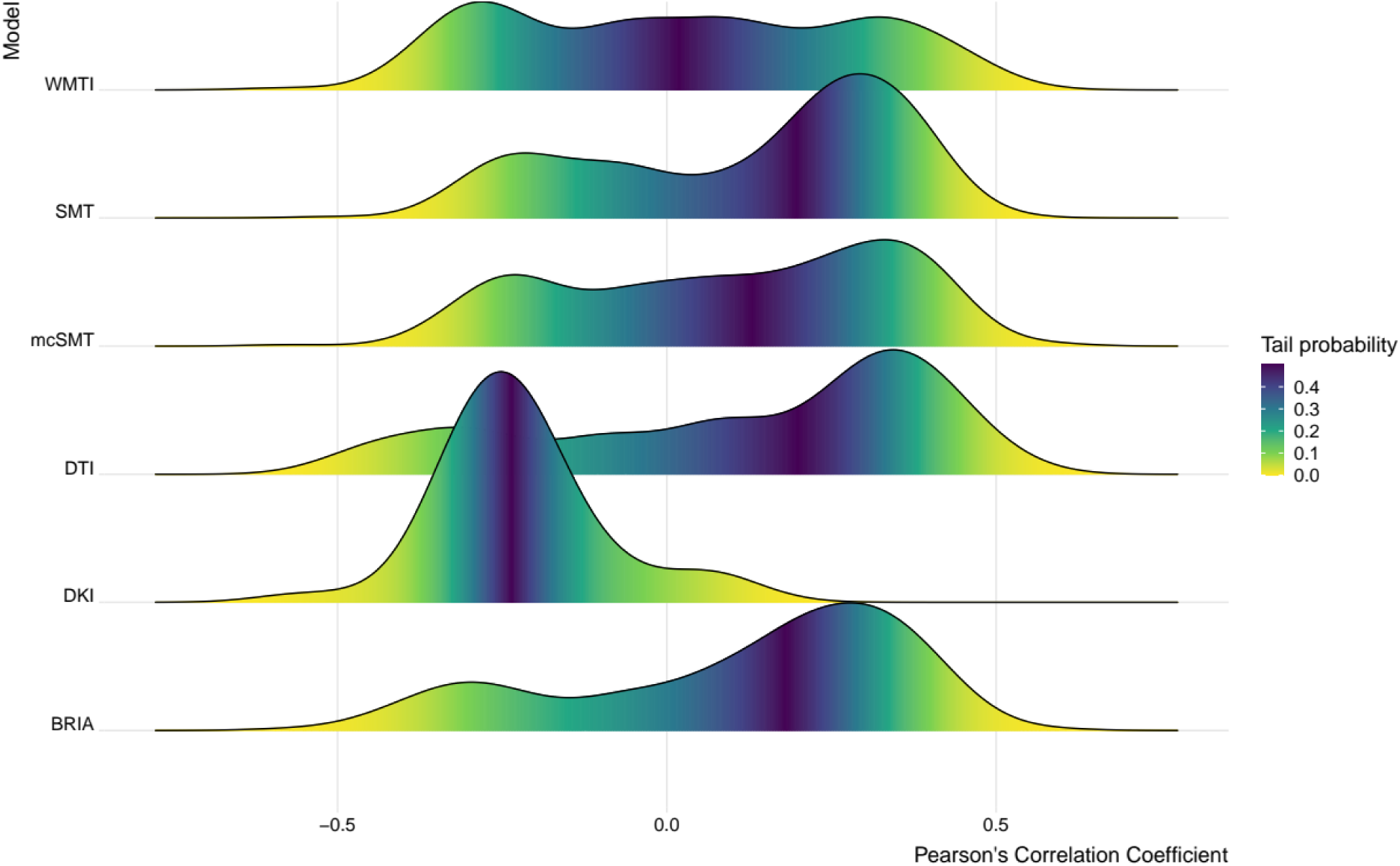
Density plots feature-age correlation across diffusion approaches with tail probabilities. This figure is a supplement to Figure 5, showing the distributions of the correlations between age and each models’ diffusion metrics.

**SF13:**
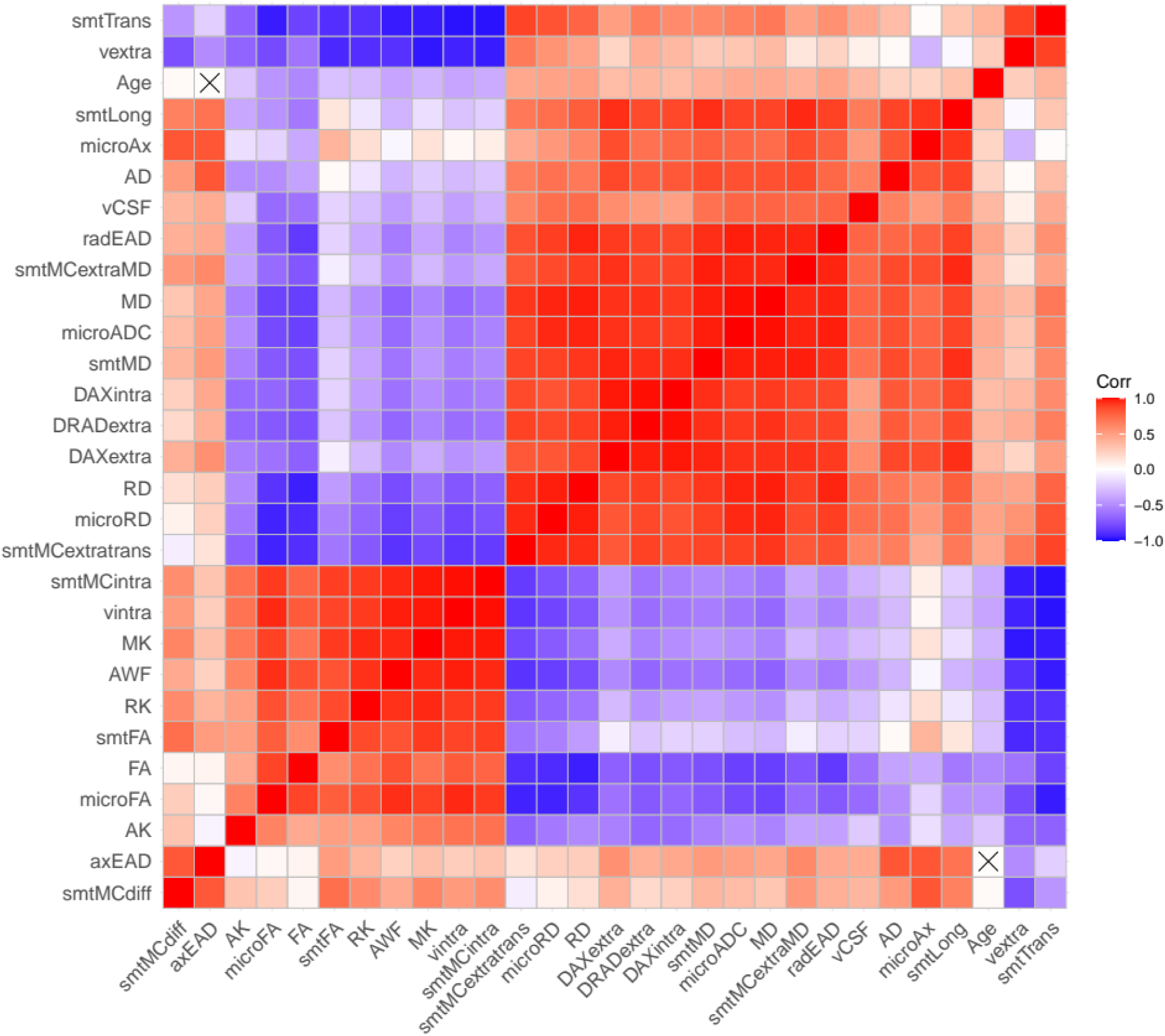
Correlations between forceps diffusion metrics and chronological age *Note*: Crossed out values were non-significant.

**SF14:**
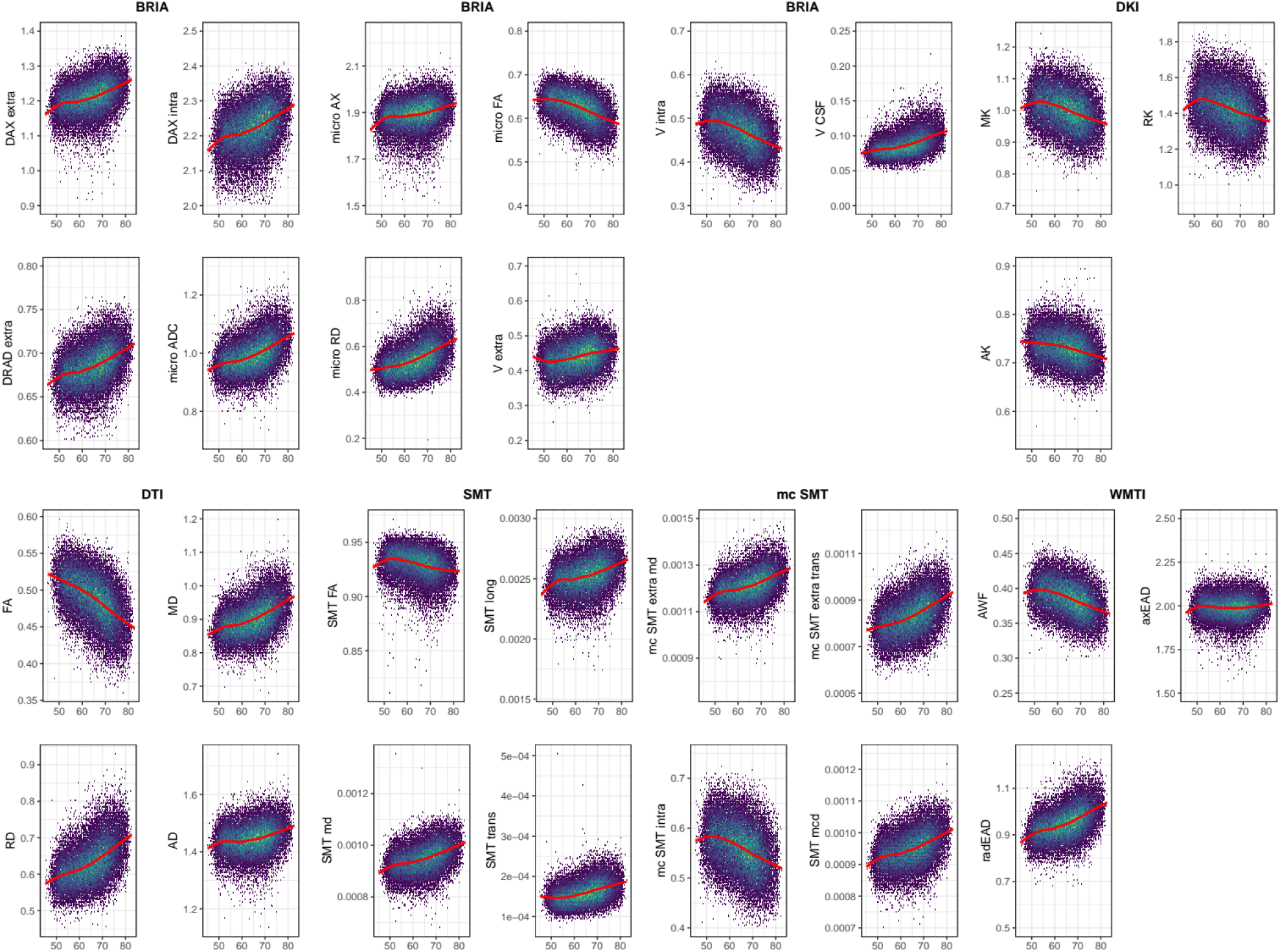
Absolute forceps diffusion metrics across age. *Note*: The presented plots represent diffusion metrics for each of the six diffusion models from the full sample *N* = 35,749 for forceps. Brighter colours indicate higher density and red lines are fitted lines to the relationship between age and diffusion metric.

**SF15:**
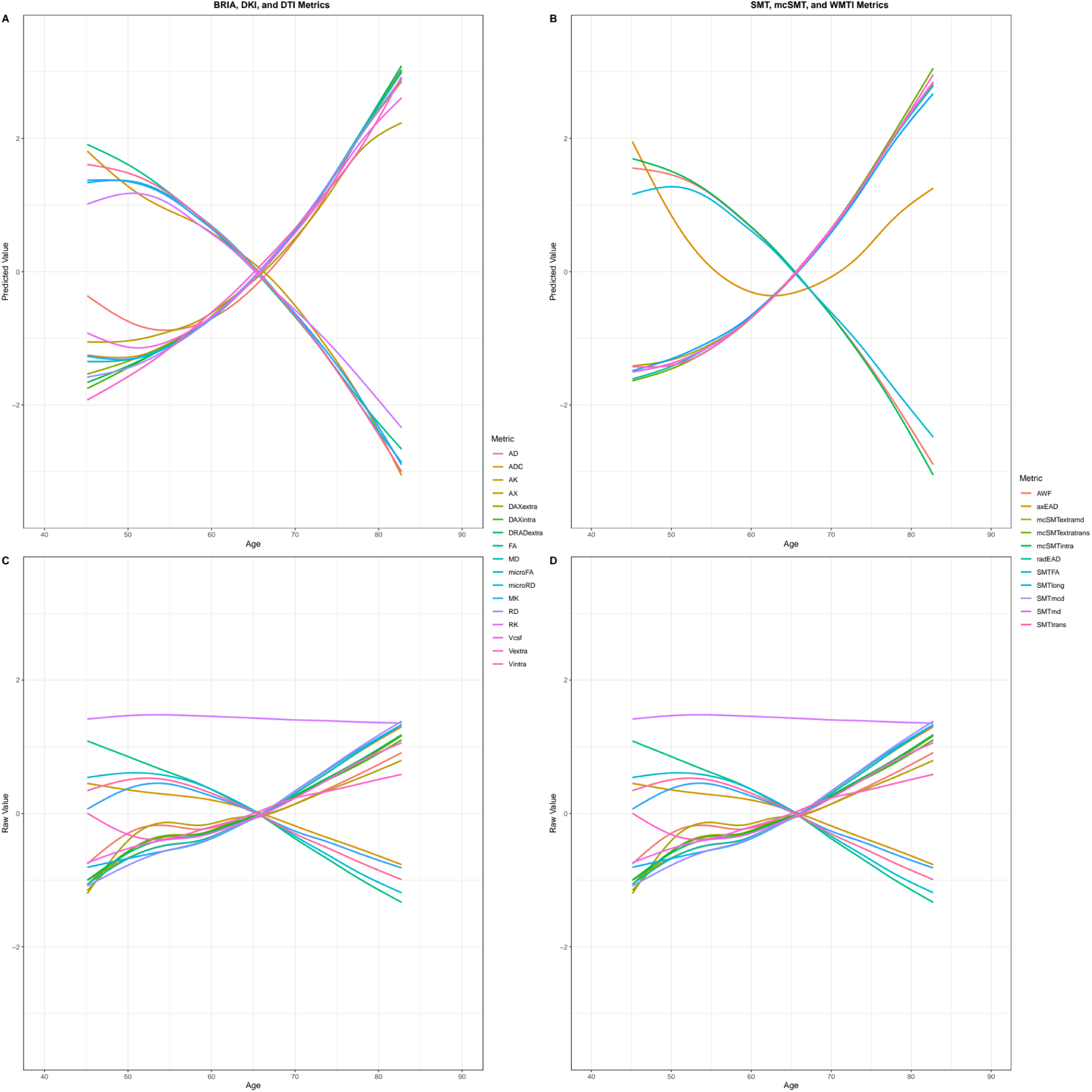
Raw and predicted forceps diffusion metrics by chronological age. SF14A-D shows age curves for each standardised (z-score) fornix diffusion skeleton value (y-axis) plotted as a function of age (x-axis). Shaded areas represent 95% CI. Curves fitted to raw values (SF9C-D) serve as a comparison to the lm- derived predicted values from Equation 1.

**SF16:**
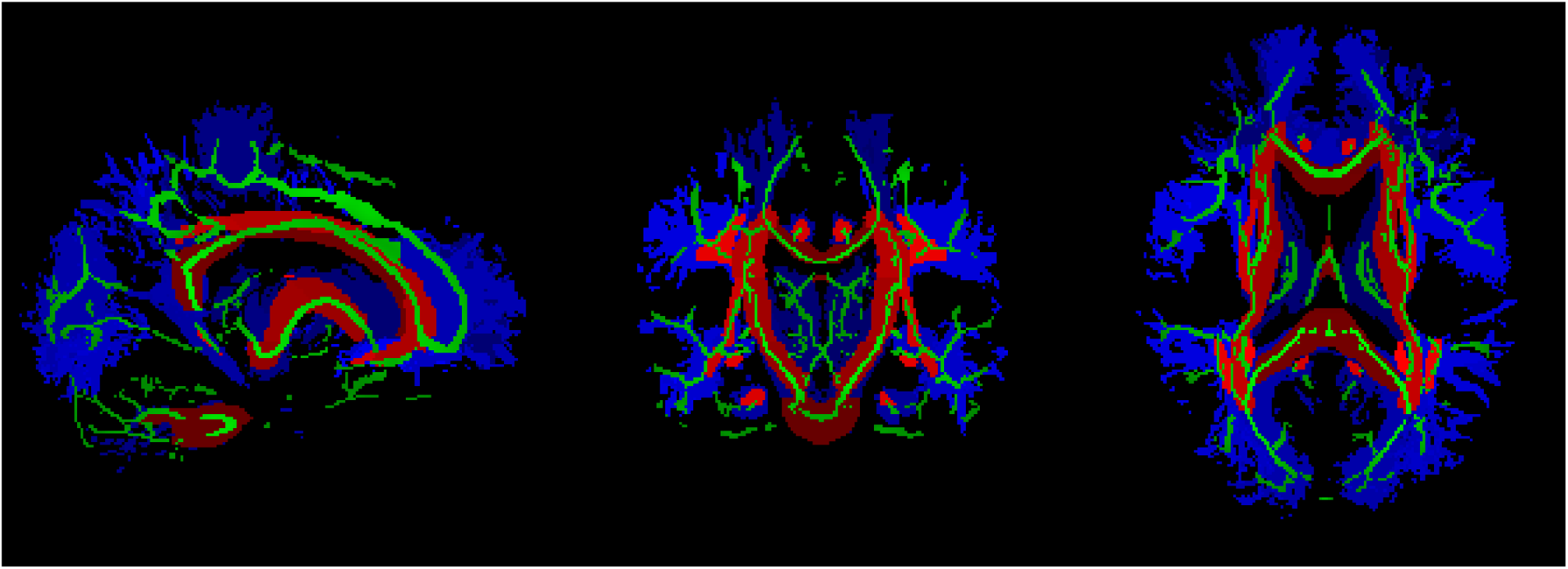
White Matter Tracts and Regions Used in this Study. Red indicates JHU labels atlas with main ROIs. Blue indicates the tractorgaphic atlas with finer skeleton structure. The green colour marks the FA skeleton.

### Supplementary Tables

**ST1:**
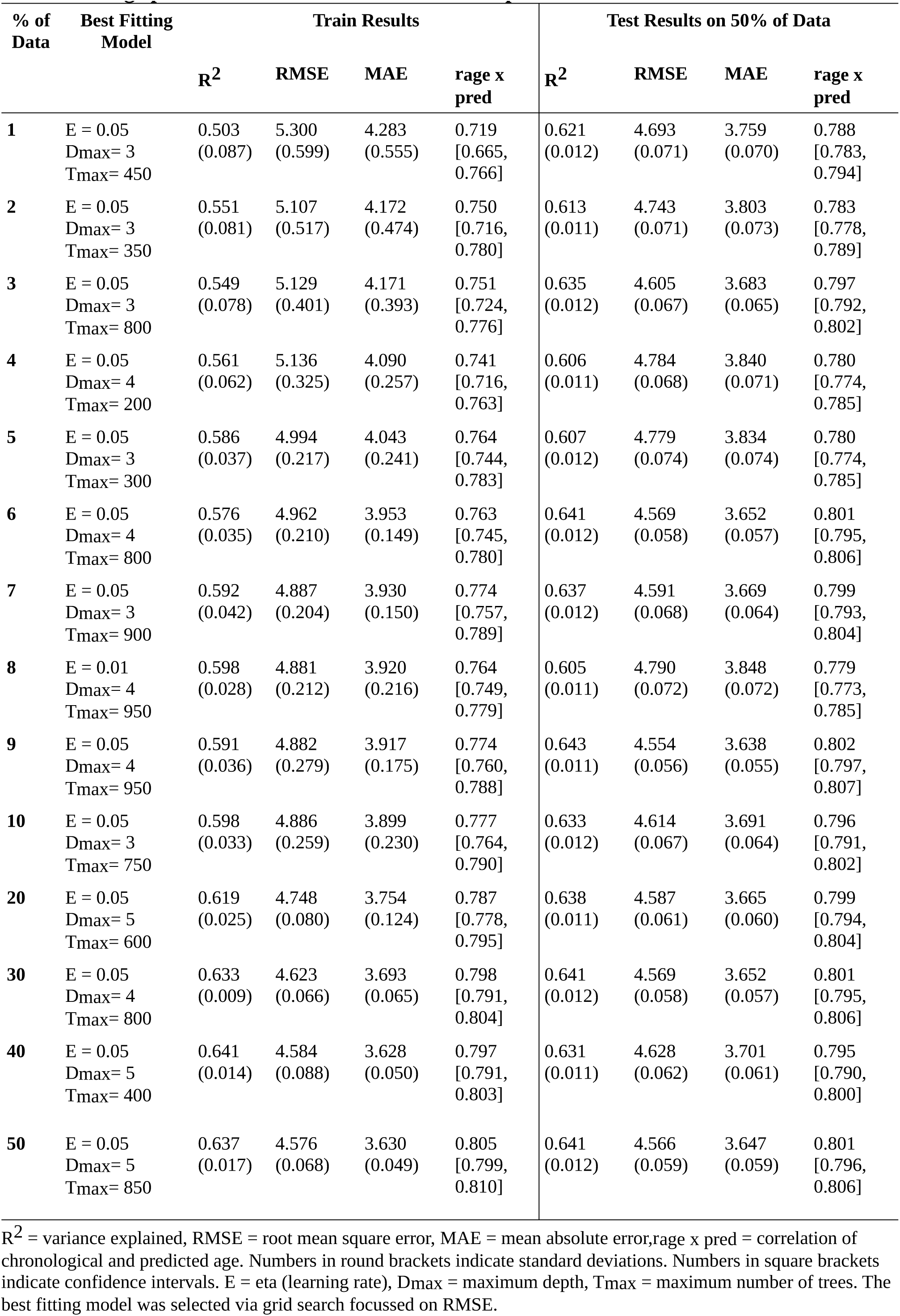
Brain age predictions from different train-test splits.

**ST2:** Differences between correlations of chronological and *corrected* predicted age across models with 95% confidence interval.

|  | BRIA | DKI | DTI | SMT | mcSMT | WMTI | Mean |
| --- | --- | --- | --- | --- | --- | --- | --- |
| **DKI** | 0.011<br>[0.009,<br>0.013] |  |  |  |  |  |  |
| **DTI** | 0.008<br>[0.007,<br>0.010] | 0.003<br>[0.002,<br>0.005] |  |  |  |  |  |
| **SMT** | 0.007<br>[0.006,<br>0.008] | 0.004<br>[0.003,<br>0.005] | 0.001 [-<br>0.0004,<br>0.0024]<sup>4</sup> |  |  |  |  |
| **mcSMT** | ≈0<br>[-0.001,<br>0.001]<sup>1</sup> | 0.011<br>[0.009,<br>0.013] | 0.008<br>[0.006,<br>0.010] | 0.007<br>[0.006,<br>0.008] |  |  |  |
| **WMTI** | 0.010<br>[0.009,<br>0.012] | 0.001<br>[-0.0004,<br>0.0024]<sup>2</sup> | 0.002<br>[0.001,<br>0.003] | 0.003<br>[0.002,<br>0.004] | 0.010<br>[0.009,<br>0.012] |  |  |
| **Mean** | 0.013<br>[0.011,<br>0.015] | 0.002<br>[0.0002,<br>0.038]<sup>3</sup> | 0.005<br>[0.003,<br>0.007] | 0.006<br>[0.004,<br>0.007] | 0.013<br>[0.011,<br>0.015] | 0.003<br>[0.001,<br>0.005] |  |
| **FULL** | 0.015<br>[0.014,<br>0.017] | 0.004<br>[0.003,<br>0.005] | 0.007<br>[0.006,<br>0.008] | 0.008<br>[0.007,<br>0.009] | 0.015<br>[0.014,<br>0.017] | 0.005<br>[0.004,<br>0.006] | 0.002<br>[0.0004,<br>0.0036] |

Confidence Intervals are based on Zou <sup>42</sup>. Unmarked differences were significant at $p < .001$ .
1 Hotelling's <sup>86</sup> $t(32171) \approx 0$ , $p \approx 1$ .
2 Hotelling's (1940) $t(32171) = 1.4232$ , $p = .1547$ .
3 Hotelling's (1940) $t(32171) = 2.2174$ , $p = .0266$ .
4 Hotelling's (1940) $t(32171) = 2.4176$ , $p = .0156$ .

**ST3:**
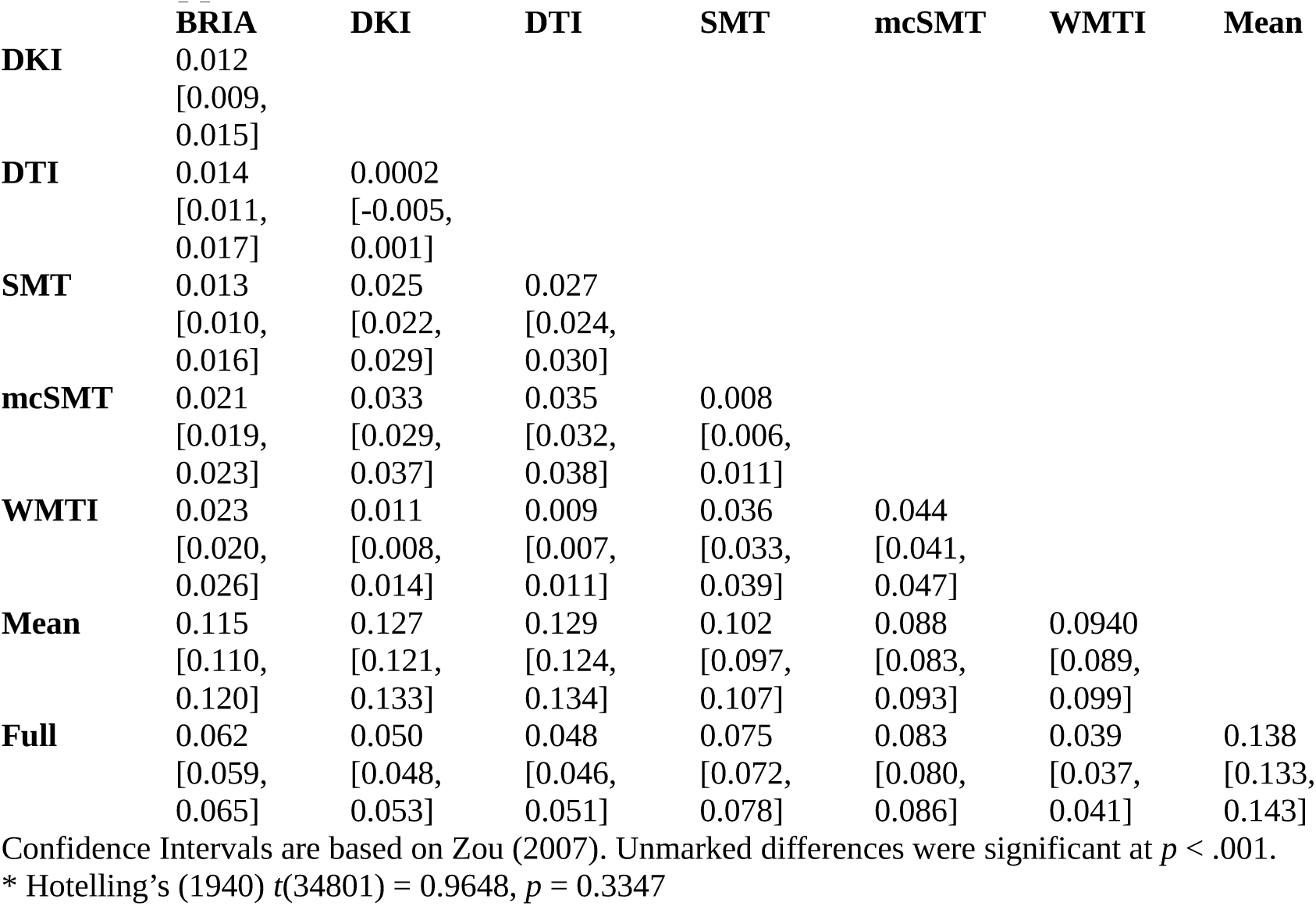
Differences between correlations of uncorrected predicted and chronological age across diffusion approaches with 95% confidence interval.

|  | BRIA | DKI | DTI | SMT | mcSMT | WMTI | Mean |
| --- | --- | --- | --- | --- | --- | --- | --- |
| **DKI** | 0.012<br>[0.009,<br>0.015] |  |  |  |  |  |  |
| **DTI** | 0.014<br>[0.011,<br>0.017] | 0.0002<br>[-0.005,<br>0.001] |  |  |  |  |  |
| **SMT** | 0.013<br>[0.010,<br>0.016] | 0.025<br>[0.022,<br>0.029] | 0.027<br>[0.024,<br>0.030] |  |  |  |  |
| **mcSMT** | 0.021<br>[0.019,<br>0.023] | 0.033<br>[0.029,<br>0.037] | 0.035<br>[0.032,<br>0.038] | 0.008<br>[0.006,<br>0.011] |  |  |  |
| **WMTI** | 0.023<br>[0.020,<br>0.026] | 0.011<br>[0.008,<br>0.014] | 0.009<br>[0.007,<br>0.011] | 0.036<br>[0.033,<br>0.039] | 0.044<br>[0.041,<br>0.047] |  |  |
| **Mean** | 0.115<br>[0.110,<br>0.120] | 0.127<br>[0.121,<br>0.133] | 0.129<br>[0.124,<br>0.134] | 0.102<br>[0.097,<br>0.107] | 0.088<br>[0.083,<br>0.093] | 0.0940<br>[0.089,<br>0.099] |  |
| **Full** | 0.062<br>[0.059,<br>0.065] | 0.050<br>[0.048,<br>0.053] | 0.048<br>[0.046,<br>0.051] | 0.075<br>[0.072,<br>0.078] | 0.083<br>[0.080,<br>0.086] | 0.039<br>[0.037,<br>0.041] | 0.138<br>[0.133,<br>0.143] |

Confidence Intervals are based on Zou (2007). Unmarked differences were significant at $p < .001$ .
\* Hotelling's (1940) $t(34801) = 0.9648$ , $p = 0.3347$

**ST4:** Fornix metrics’ age sensitivity: comparing diffusion metric prediction models with and without age.

| Model | Metric | Full<sup>1</sup> | Reduced<sup>2</sup> | $\chi^2$ | *p* | *p*Holm |
| --- | --- | --- | --- | --- | --- | --- |
| BRIA | vintra | 63395.90 | 56372.64 | 14046.51 | <.001 | <.001 |
| BRIA | vextra | 53488.40 | 47076.64 | 12823.52 | <.001 | <.001 |
| BRIA | vCSF | 32902.83 | 25747.10 | 14311.47 | <.001 | <.001 |
| BRIA | microRD | -304.68 | -7544.95 | 14480.54 | <.001 | <.001 |
| BRIA | microFA | 52365.03 | 45215.37 | 14299.32 | <.001 | <.001 |
| BRIA | microAx | 27419.04 | 23070.45 | 8697.18 | <.001 | <.001 |
| BRIA | microADC | 7489.46 | 297.02 | 14384.87 | <.001 | <.001 |
| BRIA | DRADextra | 88839.19 | 84573.36 | 8531.64 | <.001 | <.001 |
| BRIA | DAXintra | 62832.56 | 58452.18 | 8760.75 | <.001 | <.001 |
| BRIA | DAXextra | 69448.84 | 68536.57 | 1824.54 | <.001 | <.001 |
| DKI | RK | 17066.24 | 10008.43 | 14115.62 | <.001 | <.001 |
| DKI | AK | 73031.59 | 67042.47 | 11978.25 | <.001 | <.001 |
| DKI | MK | 42812.92 | 35761.79 | 14102.27 | <.001 | <.001 |
| DTI | FA | 51988.61 | 46349.96 | 11277.31 | <.001 | <.001 |
| DTI | MD | -6425.12 | -12850.70 | 12851.17 | <.001 | <.001 |
| DTI | RD | -9157.86 | -15294.22 | 12272.70 | <.001 | <.001 |
| DTI | AD | -3330.43 | -9581.75 | 12502.64 | <.001 | <.001 |
| SMT | smtFA | 28610.35 | 23053.56 | 11113.58 | <.001 | <.001 |
| SMT | smtLong | 343020.49 | 342243.25 | 1554.49 | <.001 | <.001 |
| SMT | smtMD | 253964.29 | 248676.32 | 10575.95 | <.001 | <.001 |
| SMT | smtTrans | 239584.11 | 234337.78 | 10492.66 | <.001 | <.001 |
| mcSMT | smtMCintra | 46725.61 | 39797.67 | 13855.88 | <.001 | <.001 |
| mcSMT | smtMCextraMD | 260580.57 | 253725.97 | 13709.21 | <.001 | <.001 |
| mcSMT | smtMCextratrans | 249976.89 | 242973.81 | 14006.16 | <.001 | <.001 |
| mcSMT | smtMCD | 287624.52 | 284705.87 | 5837.30 | <.001 | <.001 |
| WMTI | AWF | 73001.96 | 65850.35 | 14303.22 | <.001 | <.001 |
| WMTI | axEAD | -28343.53 | -29855.90 | 3024.74 | <.001 | <.001 |
| WMTI | radEAD | -10582.93 | -16576.91 | 11987.96 | <.001 | <.001 |

1 Full = full model log likelihood
2 Reduced = reduced model log likelihood

**ST5:**
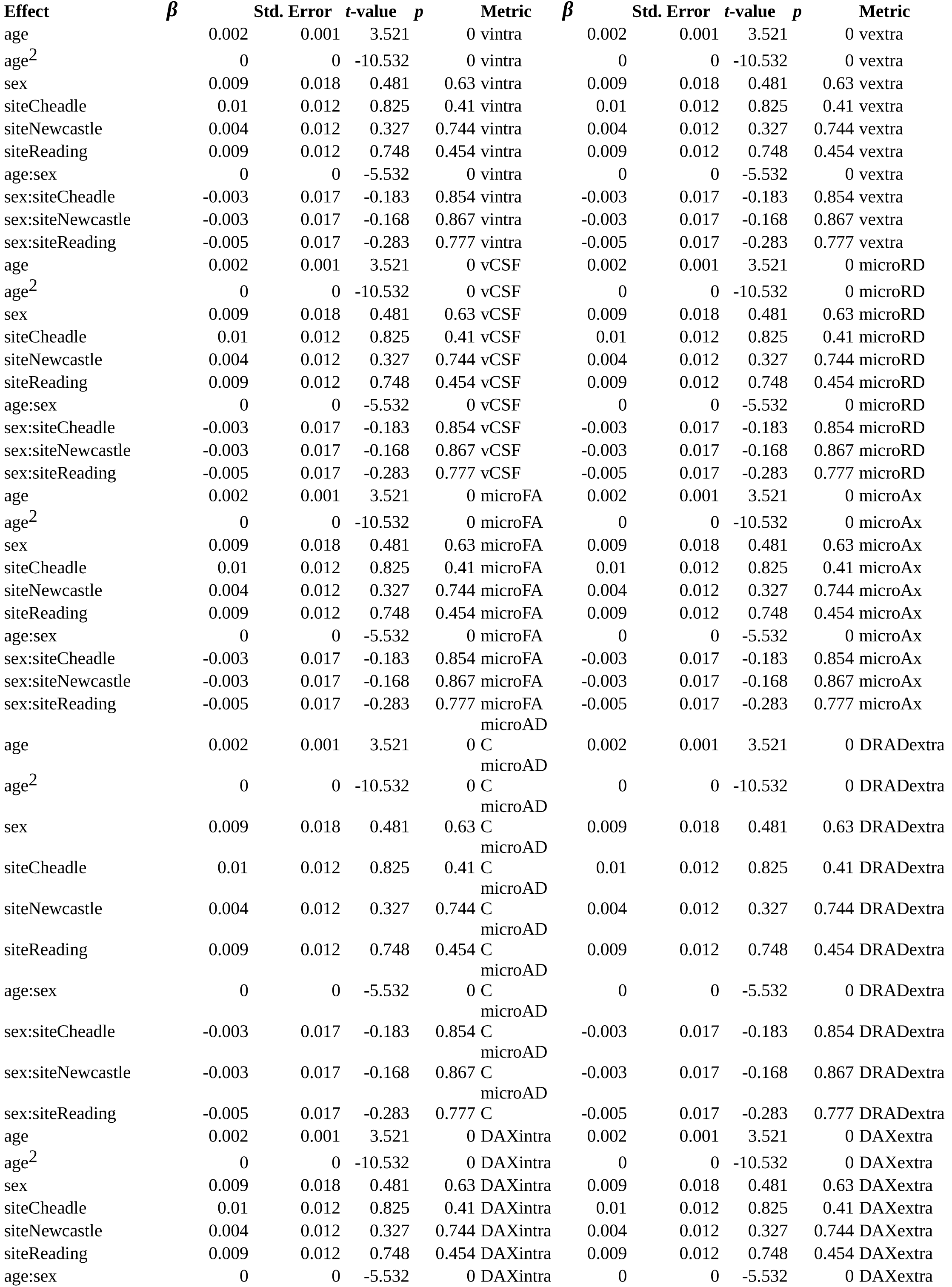
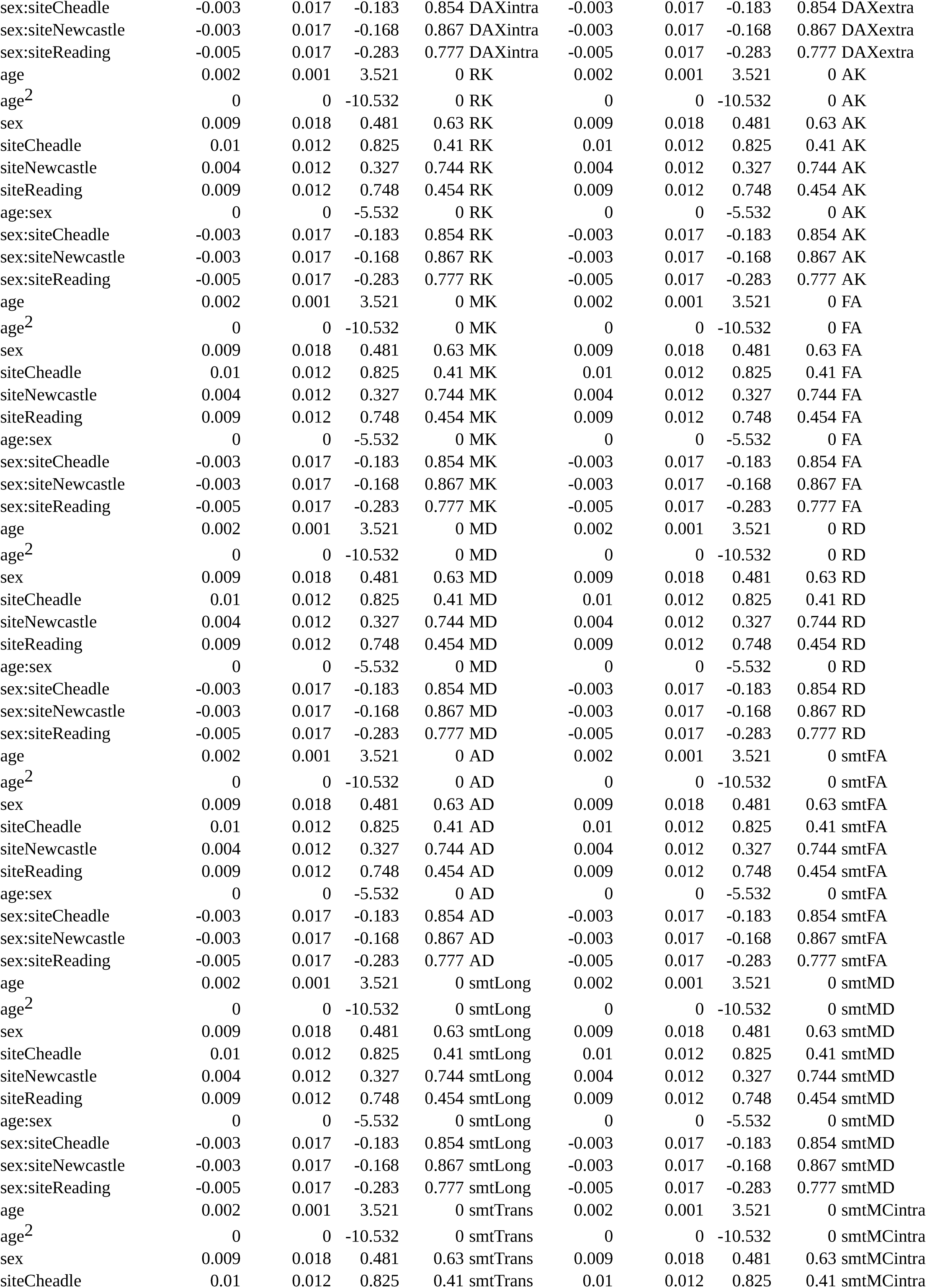
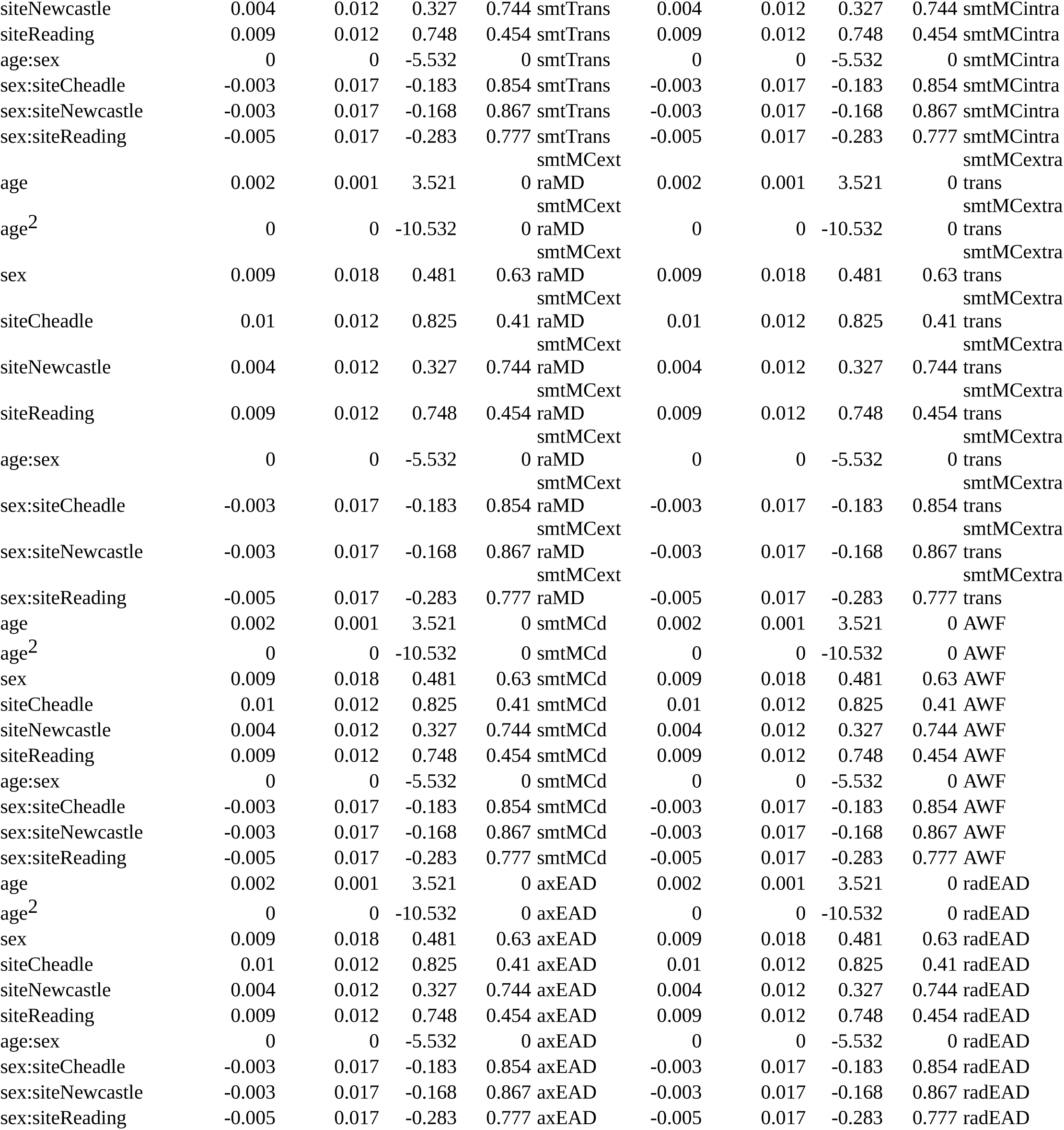
Model summaries for all 28 Fornix models.

**ST6:** Brain age prediction model performance *for data including QC outliers*.

| Name | MRI features | R<sup>2</sup> | RMSE | MAE | Prediction-Age Correlation* | Corrected Prediction-Age Correlation* |
| --- | --- | --- | --- | --- | --- | --- |
| BRIA | 700 | 0.538<br>(0.009) | 5.103<br>(0.044) | 4.096<br>(0.038) | 0.734<br>[0.729, 0.739] | 0.902<br>[0.900, 0.904] |
| DKI | 210 | 0.561<br>(0.009) | 5.078<br>(0.053) | 4.073<br>(0.047) | 0.750<br>[0.745, 0.754] | 0.876<br>[0.874, 0.879] |
| DTI | 280 | 0.565<br>(0.009) | 5.052<br>(0.039) | 4.041<br>(0.038) | 0.752<br>[0.748, 0.757] | 0.874<br>[0.872, 0.877] |
| SMT | 280 | 0.522<br>(0.009) | 5.297<br>(0.035) | 4.254<br>(0.031) | 0.723<br>[0.718, 0.728] | 0.870<br>[0.868, 0.873] |
| mcSMT | 280 | 0.508<br>(0.008) | 5.263<br>(0.040) | 4.227<br>(0.034) | 0.714<br>[0.708, 0.719] | 0.901<br>[0.899, 0.903] |
| WMTI | 210 | 0.574<br>(0.009) | 4.999<br>(0.036) | 4.003<br>(0.034) | 0.758<br>[0.754, 0.763] | 0.875<br>[0.873, 0.877] |
| Mean scores multimodal | 28 | 0.400<br>(0.068) | 5.945<br>(0.082) | 4.820<br>(0.068) | 0.633<br>[0.627, 0.639] | 0.875<br>[0.873, 0.878] |
| Full model multimodal | 1932 | 0.648<br>(0.009) | 4.557<br>(0.077) | 3.637<br>(0.066) | 0.805<br>[0.801, 0.808] | 0.877<br>[0.875, 0.880] |

R<sup>2</sup>:variance explained, RMSE: root mean squared error, MAE: mean absolute error
*Note:* R<sup>2</sup>, RMSE, MAE are displayed in the format Mean (Standard Deviation), Pearson's correlations are displayed in the format Correlation Score 95% Confidence Interval [Lower Bound, Upper Bound].
\*All correlation were significant at $p < .001$ .

**ST7:** Top five diffusion metrics ranked by gain in age prediction accuracy for data including QC outliers.

| BRIA | DKI | DTI | SMT | mcSMT | WMTI | Full |
| --- | --- | --- | --- | --- | --- | --- |
| Micro RD<br>ATRR (67436) | MK Fornix<br>(99885) | MD Fornix<br>(79767) | MD Fornix<br>(72129) | Extratrans<br>Fornix<br>(55429) | AWF Fornix<br>(69033) | AWF fornix<br>(278977) |
| Micro FA Fornix<br>(63804) | RK Fornix<br>(28196) | RD left anterior<br>corona radiata<br>(51592) | MD FMIN<br>(48161) | Intra Fornix<br>(46212) | RadEAD<br>anterior right<br>corona radiata<br>(34381) | Micro RD fornix<br>(71175) |
| Micro RD right<br>external capsule<br>(27069) | AK anterior<br>right limb of<br>internal capsule<br>(22725) | RD FMIN<br>(25201) | MD right<br>anterior corona<br>radiata<br>(44331) | Extratrans right<br>external capsule<br>(16611) | RadEAD IFOFR<br>(22512) | Micro FA<br>fornix(35049) |
| Micro RD<br>Fornix right<br>striaterminalis<br>(17090) | AK Fornix<br>(14401) | RD Fornix right<br>stria terminalis<br>(22951) | FA Fornix<br>(19697) | ExtraMD Fornix<br>(10267) | RadEAD FMIN<br>(20286) | MD right<br>tapetum<br>(34008) |
| Micro FA FMIN<br>(14335) | AK superior<br>frontooccipital<br>left fasciculus<br>(7978) | MD anterior<br>limb of internal<br>left capsule<br>(15589) | Long left<br>tapetum<br>(14596) | ExtraMD<br>anterior left limb<br>of internal<br>capsule<br>(7830) | RadEAD ATRL<br>(16666) | RadEAD<br>anterior right<br>corona radiata<br>(27907) |

Table values can be read as feature name (gain). Mean refers to the multimodal model containing only mean scores and full to the full model containing all features. Cells including Fornix are marked in green. laLC = left anterior limb of internal capsule; raLC = right anterior limb of internal capsule; lST = lSTria terminalis; rST = rSTria terminalis; lsfoF = left superior frontal occipital fasciculus; laCR = left anterior corona radiata; raCR = right anterior corona radiata; rEC = right external capsule

**ST8:**
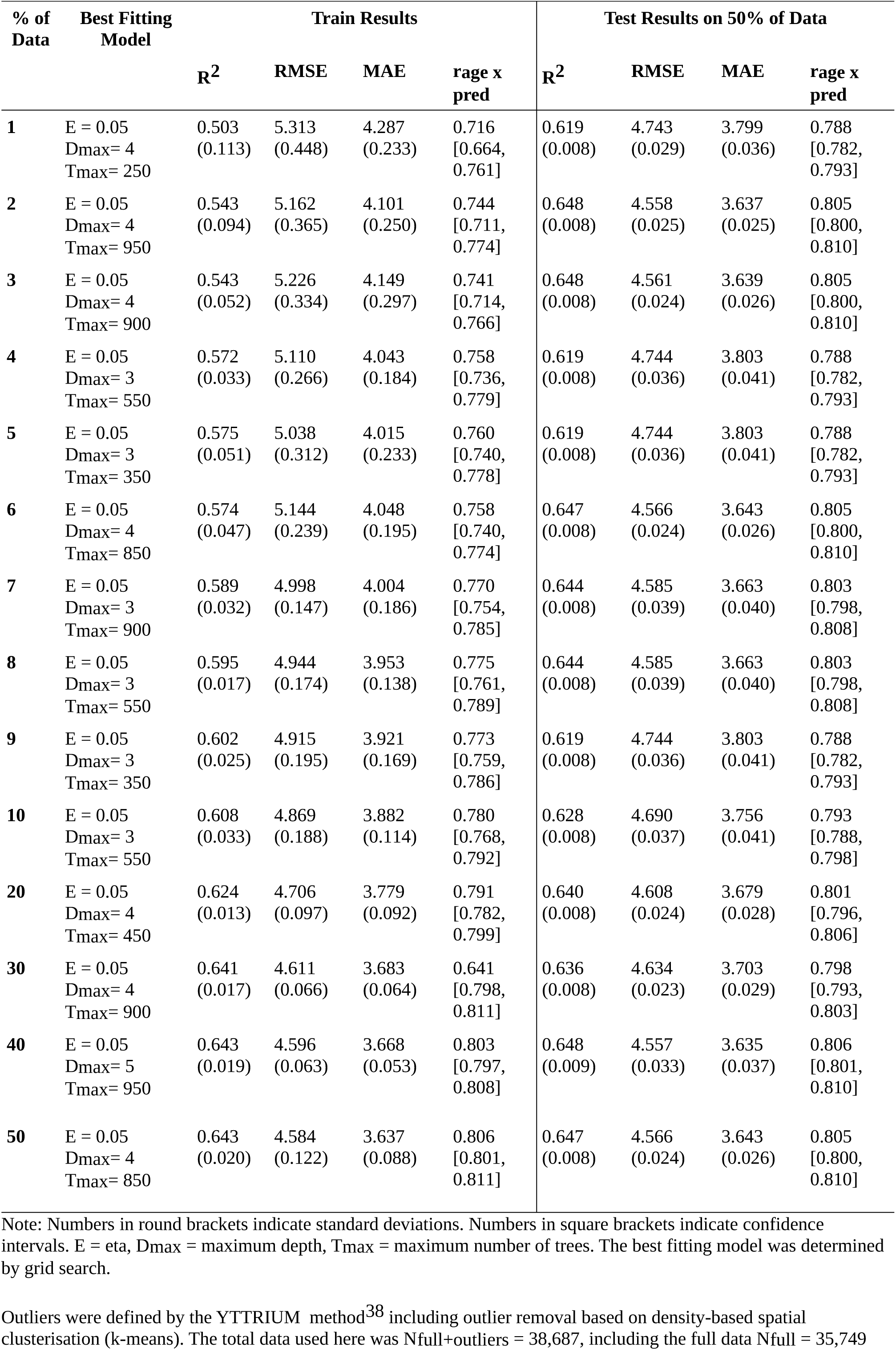
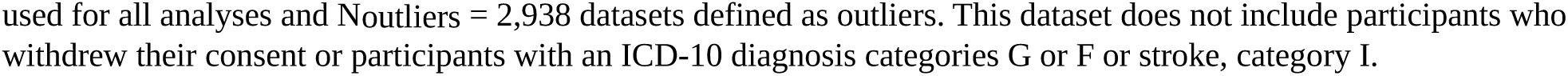
Brain age predictions from different train-test splits *for data including QC outliers*.

| % of Data | Best Fitting Model | Train Results |  |  |  | Test Results on 50% of Data |  |  |  |
| --- | --- | --- | --- | --- | --- | --- | --- | --- | --- |
|  |  | R<sup>2</sup> | RMSE | MAE | rage x pred | R<sup>2</sup> | RMSE | MAE | rage x pred |
| 1 | E = 0.05<br>Dmax= 4<br>Tmax= 250 | 0.503<br>(0.113) | 5.313<br>(0.448) | 4.287<br>(0.233) | 0.716<br>[0.664,<br>0.761] | 0.619<br>(0.008) | 4.743<br>(0.029) | 3.799<br>(0.036) | 0.788<br>[0.782,<br>0.793] |
| 2 | E = 0.05<br>Dmax= 4<br>Tmax= 950 | 0.543<br>(0.094) | 5.162<br>(0.365) | 4.101<br>(0.250) | 0.744<br>[0.711,<br>0.774] | 0.648<br>(0.008) | 4.558<br>(0.025) | 3.637<br>(0.025) | 0.805<br>[0.800,<br>0.810] |
| 3 | E = 0.05<br>Dmax= 4<br>Tmax= 900 | 0.543<br>(0.052) | 5.226<br>(0.334) | 4.149<br>(0.297) | 0.741<br>[0.714,<br>0.766] | 0.648<br>(0.008) | 4.561<br>(0.024) | 3.639<br>(0.026) | 0.805<br>[0.800,<br>0.810] |
| 4 | E = 0.05<br>Dmax= 3<br>Tmax= 550 | 0.572<br>(0.033) | 5.110<br>(0.266) | 4.043<br>(0.184) | 0.758<br>[0.736,<br>0.779] | 0.619<br>(0.008) | 4.744<br>(0.036) | 3.803<br>(0.041) | 0.788<br>[0.782,<br>0.793] |
| 5 | E = 0.05<br>Dmax= 3<br>Tmax= 350 | 0.575<br>(0.051) | 5.038<br>(0.312) | 4.015<br>(0.233) | 0.760<br>[0.740,<br>0.778] | 0.619<br>(0.008) | 4.744<br>(0.036) | 3.803<br>(0.041) | 0.788<br>[0.782,<br>0.793] |
| 6 | E = 0.05<br>Dmax= 4<br>Tmax= 850 | 0.574<br>(0.047) | 5.144<br>(0.239) | 4.048<br>(0.195) | 0.758<br>[0.740,<br>0.774] | 0.647<br>(0.008) | 4.566<br>(0.024) | 3.643<br>(0.026) | 0.805<br>[0.800,<br>0.810] |
| 7 | E = 0.05<br>Dmax= 3<br>Tmax= 900 | 0.589<br>(0.032) | 4.998<br>(0.147) | 4.004<br>(0.186) | 0.770<br>[0.754,<br>0.785] | 0.644<br>(0.008) | 4.585<br>(0.039) | 3.663<br>(0.040) | 0.803<br>[0.798,<br>0.808] |
| 8 | E = 0.05<br>Dmax= 3<br>Tmax= 550 | 0.595<br>(0.017) | 4.944<br>(0.174) | 3.953<br>(0.138) | 0.775<br>[0.761,<br>0.789] | 0.644<br>(0.008) | 4.585<br>(0.039) | 3.663<br>(0.040) | 0.803<br>[0.798,<br>0.808] |
| 9 | E = 0.05<br>Dmax= 3<br>Tmax= 350 | 0.602<br>(0.025) | 4.915<br>(0.195) | 3.921<br>(0.169) | 0.773<br>[0.759,<br>0.786] | 0.619<br>(0.008) | 4.744<br>(0.036) | 3.803<br>(0.041) | 0.788<br>[0.782,<br>0.793] |
| 10 | E = 0.05<br>Dmax= 3<br>Tmax= 550 | 0.608<br>(0.033) | 4.869<br>(0.188) | 3.882<br>(0.114) | 0.780<br>[0.768,<br>0.792] | 0.628<br>(0.008) | 4.690<br>(0.037) | 3.756<br>(0.041) | 0.793<br>[0.788,<br>0.798] |
| 20 | E = 0.05<br>Dmax= 4<br>Tmax= 450 | 0.624<br>(0.013) | 4.706<br>(0.097) | 3.779<br>(0.092) | 0.791<br>[0.782,<br>0.799] | 0.640<br>(0.008) | 4.608<br>(0.024) | 3.679<br>(0.028) | 0.801<br>[0.796,<br>0.806] |
| 30 | E = 0.05<br>Dmax= 4<br>Tmax= 900 | 0.641<br>(0.017) | 4.611<br>(0.066) | 3.683<br>(0.064) | 0.641<br>[0.798,<br>0.811] | 0.636<br>(0.008) | 4.634<br>(0.023) | 3.703<br>(0.029) | 0.798<br>[0.793,<br>0.803] |
| 40 | E = 0.05<br>Dmax= 5<br>Tmax= 950 | 0.643<br>(0.019) | 4.596<br>(0.063) | 3.668<br>(0.053) | 0.803<br>[0.797,<br>0.808] | 0.648<br>(0.009) | 4.557<br>(0.033) | 3.635<br>(0.037) | 0.806<br>[0.801,<br>0.810] |
| 50 | E = 0.05<br>Dmax= 4<br>Tmax= 850 | 0.643<br>(0.020) | 4.584<br>(0.122) | 3.637<br>(0.088) | 0.806<br>[0.801,<br>0.811] | 0.647<br>(0.008) | 4.566<br>(0.024) | 3.643<br>(0.026) | 0.805<br>[0.800,<br>0.810] |

Note: Numbers in round brackets indicate standard deviations. Numbers in square brackets indicate confidence intervals. E = eta, Dmax = maximum depth, Tmax = maximum number of trees. The best fitting model was determined by grid search.
Outliers were defined by the YTTRIUM method<sup>38</sup> including outlier removal based on density-based spatial clusterisation (k-means). The total data used here was Nfull+outliers = 38,687, including the full data Nfull = 35,749
used for all analyses and Noutliers = 2,938 datasets defined as outliers. This dataset does not include participants who withdrew their consent or participants with an ICD-10 diagnosis categories G or F or stroke, category I.

**ST9:** Comparisons of linear and generalized additive models predicting fornix diffusion metrics.

| Metric | LM AIC | GAM AIC | LM BIC | GAM BIC | LM R<sup>2</sup><sub>adj</sub> | GAM R<sup>2</sup><sub>adj</sub> |
| --- | --- | --- | --- | --- | --- | --- |
| vintra | -126767.79 | -126755.60 | -126665.98 | -126662.27 | 0.36 | 0.36 |
| vextra | -106952.81 | -106929.99 | -106851.00 | -106836.66 | 0.38 | 0.37 |
| vCSF | -65781.67 | -65773.53 | -65679.86 | -65680.20 | 0.39 | 0.39 |
| microRD | 633.36 | 643.11 | 735.18 | 736.43 | 0.39 | 0.39 |
| microFA | -104706.06 | -104634.30 | -104604.25 | -104540.97 | 0.38 | 0.38 |
| microAx | -54814.08 | -54790.59 | -54712.27 | -54697.26 | 0.27 | 0.27 |
| microADC | -14954.91 | -14946.59 | -14853.10 | -14853.26 | 0.39 | 0.39 |
| DRADextra | -177654.37 | -177648.58 | -177552.56 | -177555.25 | 0.30 | 0.30 |
| DAXintra | -125641.12 | -125639.53 | -125539.31 | -125546.20 | 0.30 | 0.30 |
| DAXextra | -138873.67 | -138808.20 | -138771.86 | -138714.87 | 0.09 | 0.09 |
| RK | -34108.48 | -34109.41 | -34006.67 | -34016.08 | 0.37 | 0.37 |
| AK | -146039.18 | -146041.08 | -145937.37 | -145947.75 | 0.32 | 0.32 |
| MK | -85601.85 | -85603.57 | -85500.04 | -85510.24 | 0.36 | 0.36 |
| FA | -103953.22 | -103924.62 | -103851.41 | -103831.30 | 0.31 | 0.31 |
| MD | 12874.23 | 12878.93 | 12976.04 | 12972.25 | 0.35 | 0.35 |
| RD | 18339.73 | 18359.69 | 18441.54 | 18453.02 | 0.34 | 0.34 |
| AD | 6684.86 | 6689.75 | 6786.67 | 6783.08 | 0.35 | 0.35 |
| smtFA | -57196.69 | -57161.31 | -57094.88 | -57067.98 | 0.31 | 0.31 |
| smtLong | -686016.98 | -685869.48 | -685915.17 | -685776.15 | 0.06 | 0.05 |
| smtMD | -507904.58 | -507869.45 | -507802.77 | -507776.13 | 0.30 | 0.30 |
| smtTrans | -479144.23 | -479104.60 | -479042.42 | -479011.28 | 0.30 | 0.29 |
| smtMCintra | -93427.21 | -93427.11 | -93325.40 | -93333.78 | 0.36 | 0.36 |
| smtMCextraMD | -521137.14 | -521125.59 | -521035.33 | -521032.26 | 0.37 | 0.37 |
| smtMCextratrans | -499929.78 | -499929.66 | -499827.97 | -499836.34 | 0.37 | 0.37 |
| smtMCD | -575225.03 | -574996.85 | -575123.22 | -574903.53 | 0.22 | 0.21 |
| AWF | -145979.93 | -145981.86 | -145878.11 | -145888.53 | 0.37 | 0.37 |
| axEAD | 56711.05 | 56715.91 | 56812.86 | 56809.24 | 0.10 | 0.10 |
| radEAD | 21189.85 | 21203.80 | 21291.67 | 21297.12 | 0.33 | 0.33 |

LM = linear model, GAM = generalized additive model, AIC = Akaike information criterion, BIC = Bayesian information criterion. The numbers are derived from the six diffusion approaches' 28 metrics following Equation 1 for linear models and all variables of the equation allowing splines for non-linear models.

**ST10:** Overview of diffusion metrics by diffusion approach.

ST10: Overview of diffusion metrics by diffusion approach
| Diffusion Approach | Metrics |
| --- | --- |
| Bayesian Rotationally Invariant Approach (BRIA) | intra-axonal axial diffusivity (DAX intra)<br>extra-axonal radial diffusivity (DRAD extra)<br>microscopic fractional anisotropy (micro FA)<br>extra-axonal axial diffusivity (DAX extra)<br>intra-axonal water fraction (V intra)<br>extra-axonal water fraction (V extra)<br>cerebrospinal fluid fraction (vCSF)<br>microscopical axial diffusivity (micro AX)<br>microscopic radial diffusivity (micro RD)<br>microscopical apparent diffusion coefficient (micro ADC) |
| Diffusion Kurtosis Imaging (DKI) | mean kurtosis (MK)<br>radial kurtosis (RK)<br>axial kurtosis (AK) |
| Diffusion Tensor Imaging (DTI) | fractional anisotropy (FA)<br>axial diffusivity (AD)<br>mean diffusivity (MD)<br>radial diffusivity (RD) |
| Spherical Mean Technique (SMT) | fractional anisotropy (SMT FA)<br>mean diffusivity (SMT md)<br>transverse diffusion coefficient (SMT trans)<br>longitudinal diffusion coefficient (SMT long) |
| Multi-compartment Spherical Mean Technique (mcSMT) | extra-neurite microscopic mean diffusivity (mcSMT extra md)<br>extra-neurite transverse microscopic diffusivity (mcSMT extra trans)<br>mc SMTdiffusion coefficient (SMT mcd)<br>intra-neurite volume fraction (mcSMT intra) |
| White Matter Tract Integrity (WMTI) | axonal water fraction (AWF)<br>radial extra-axonal diffusivity (radEAD)<br>axial extra-axonal diffusivity (axEAD) |

**ST11.** Whole-brain metrics’ age sensitivity: comparing diffusion metric prediction models with and without age.

| Metric | Full<sup>1</sup> | Reduced<sup>2</sup> | $\chi^2$ | *p* | *p*Holm |
| --- | --- | --- | --- | --- | --- |
| vintra | 77831.60 | 75474.15 | 4714.90 | <.001 | <.001 |
| vextra | 80010.15 | 79172.34 | 1675.61 | <.001 | <.001 |
| vCSF | 104258.51 | 102045.76 | 4425.50 | <.001 | <.001 |
| microRD | 67462.88 | 62714.78 | 9496.19 | <.001 | <.001 |
| microFA | 91462.73 | 87450.56 | 8024.34 | <.001 | <.001 |
| microAx | 68450.93 | 67363.35 | 2175.17 | <.001 | <.001 |
| microADC | 72076.62 | 67934.52 | 8284.22 | <.001 | <.001 |
| DRADextra | 105039.68 | 102688.39 | 4702.58 | <.001 | <.001 |
| DAXintra | 66812.02 | 64826.61 | 3970.82 | <.001 | <.001 |
| DAXextra | 80174.07 | 77914.51 | 4519.12 | <.001 | <.001 |
| RK | 44127.87 | 41757.04 | 4741.66 | <.001 | <.001 |
| AK | 87172.98 | 85483.34 | 3379.29 | <.001 | <.001 |
| MK | 66245.64 | 64166.15 | 4158.99 | <.001 | <.001 |
| FA | 93186.73 | 88785.16 | 8803.13 | <.001 | <.001 |
| MD | 76490.81 | 72273.83 | 8433.95 | <.001 | <.001 |
| RD | 73140.15 | 68320.02 | 9640.26 | <.001 | <.001 |
| AD | 76516.84 | 74699.38 | 3634.93 | <.001 | <.001 |
| smtFA | 125878.97 | 124092.64 | 3572.65 | <.001 | <.001 |
| smtLong | 287398.36 | 285277.70 | 4241.31 | <.001 | <.001 |
| smtMD | 320273.27 | 317056.32 | 6433.91 | <.001 | <.001 |
| smtTrans | 342834.04 | 339946.36 | 5775.35 | <.001 | <.001 |
| smtMCintra | 74365.62 | 72478.15 | 3774.94 | <.001 | <.001 |
| smtMCextraMD | 314618.75 | 311214.26 | 6809.00 | <.001 | <.001 |
| smtMCextratrans | 303909.44 | 300508.10 | 6802.69 | <.001 | <.001 |
| smtMCd | 290508.70 | 290389.47 | 238.47 | <.001 | <.001 |
| AWF | 102689.44 | 100136.63 | 5105.61 | <.001 | <.001 |
| axEAD | -14423.53 | -14426.86 | 6.66 | 0.08 | 0.08 |
| radEAD | 10331.86 | 10122.73 | 418.26 | <.001 | <.001 |

1 Full = full model log likelihood 2 Reduced = reduced model log likelihood

**ST12.** Comparisons of linear and generalized additive models predicting whole-brain diffusion metrics.

| Metric | LM AIC | GAM AIC | LM BIC | GAM BIC | LM R<sup>2</sup><sub>adj</sub> | GAM R<sup>2</sup><sub>adj</sub> |
| --- | --- | --- | --- | --- | --- | --- |
| vintra | -155639.20 | -155638.58 | -155537.39 | -155545.25 | 0.13 | 0.13 |
| vextra | -159996.29 | -159998.05 | -159894.48 | -159904.72 | 0.05 | 0.05 |
| vCSF | -208493.03 | -208487.27 | -208391.21 | -208393.94 | 0.13 | 0.13 |
| microRD | -134901.75 | -134894.29 | -134799.94 | -134800.96 | 0.25 | 0.25 |
| microFA | -182901.46 | -182896.42 | -182799.64 | -182803.09 | 0.21 | 0.21 |
| microAx | -136877.86 | -136869.60 | -136776.05 | -136776.28 | 0.08 | 0.08 |
| microADC | -144129.25 | -144118.74 | -144027.44 | -144025.42 | 0.23 | 0.23 |
| DRADextra | -210055.37 | -210046.77 | -209953.56 | -209953.44 | 0.13 | 0.13 |
| DAXintra | -133600.03 | -133591.04 | -133498.22 | -133497.71 | 0.11 | 0.11 |
| DAXextra | -160324.15 | -160312.75 | -160222.34 | -160219.42 | 0.13 | 0.13 |
| RK | -88231.74 | -88223.96 | -88129.93 | -88130.63 | 0.14 | 0.14 |
| AK | -174321.97 | -174323.26 | -174220.16 | -174229.93 | 0.09 | 0.09 |
| MK | -132467.28 | -132465.03 | -132365.47 | -132371.70 | 0.12 | 0.12 |
| FA | -186349.46 | -186338.20 | -186247.65 | -186244.88 | 0.23 | 0.23 |
| MD | -152957.62 | -152950.05 | -152855.81 | -152856.72 | 0.22 | 0.22 |
| RD | -146256.29 | -146248.25 | -146154.48 | -146154.92 | 0.24 | 0.24 |
| AD | -153009.69 | -153006.22 | -152907.88 | -152912.90 | 0.15 | 0.15 |
| smtFA | -251733.94 | -251735.36 | -251632.13 | -251642.03 | 0.10 | 0.10 |
| smtLong | -574772.72 | -574753.50 | -574670.90 | -574660.18 | 0.12 | 0.12 |
| smtMD | -640522.55 | -640508.56 | -640420.74 | -640415.23 | 0.18 | 0.18 |
| smtTrans | -685644.08 | -685645.94 | -685542.27 | -685552.61 | 0.16 | 0.16 |
| smtMCintra | -148707.24 | -148707.45 | -148605.43 | -148614.13 | 0.10 | 0.10 |
| smtMCextraMD | -629213.51 | -629198.88 | -629111.69 | -629105.56 | 0.18 | 0.18 |
| smtMCextratrans | -607794.88 | -607787.65 | -607693.07 | -607694.32 | 0.18 | 0.18 |
| smtMCd | -580993.40 | -580991.12 | -580891.59 | -580897.79 | 0.01 | 0.01 |
| AWF | -205354.88 | -205349.62 | -205253.07 | -205256.29 | 0.14 | 0.14 |
| axEAD | 28871.07 | 28869.86 | 28972.88 | 28963.19 | 0.00 | 0.00 |
| radEAD | -20639.72 | -20639.85 | -20537.91 | -20546.52 | 0.01 | 0.01 |

LM = linear model, GAM = generalized additive model, AIC = Akaike information criterion, BIC = Bayesian information criterion. The numbers are derived from the six diffusion approaches' 28 metrics following Equation 1 for linear models and all variables of the equation allowing splines for non-linear models.

**ST13.**
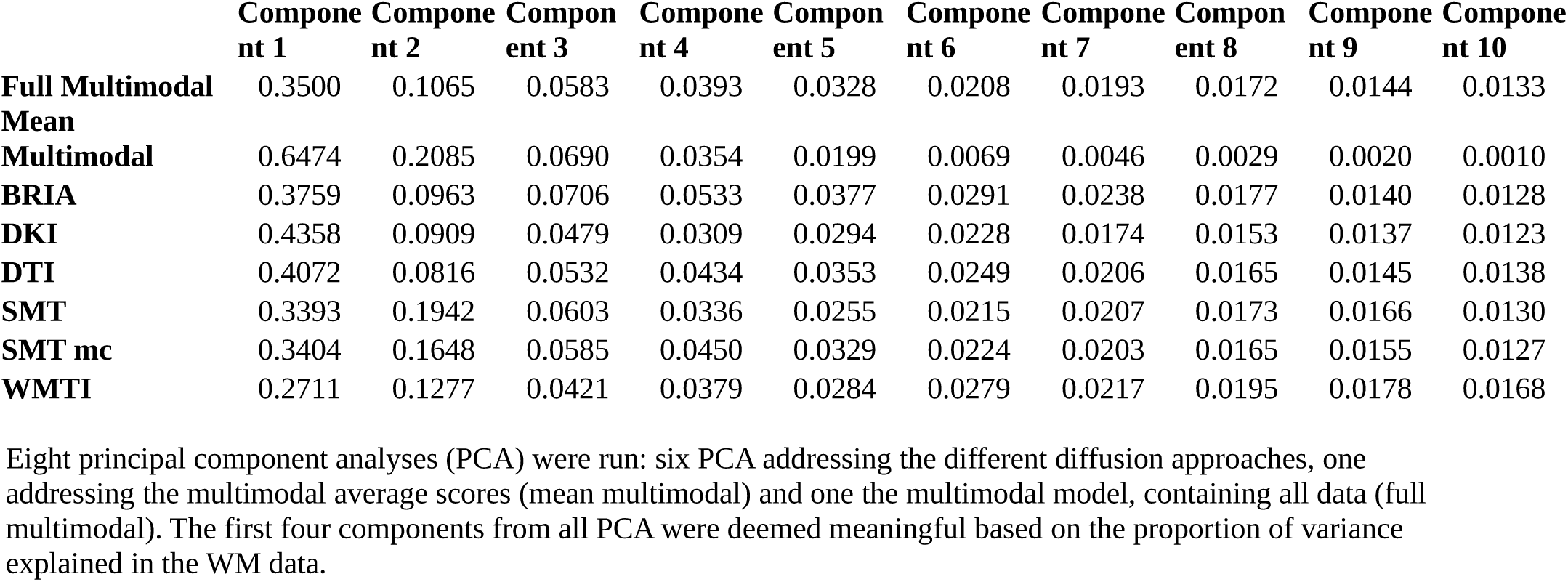
Variance explained by principal components of white matter metrics.

**ST14.** Model performance and BAG beta values for multimodal and diffusion-approach specific principal component predictions from multimodal and diffusion approach-specific BAG and covariates

| Predicted Component | Approach | $R^2$ | $R_{adj}^2$ | $T$ | $p$ | $bBAG$ |
| --- | --- | --- | --- | --- | --- | --- |
| 1 | Full Multimodal | 0.0141 | 0.0138 | 46.0160 | <0.0001 | -0.0151 |
| 1 | Mean Multimodal | 0.5054 | 0.5052 | 3651.2999 | <0.0001 | -0.6726 |
| 1 | BRIA | 0.3439 | 0.3437 | 1685.8645 | <0.0001 | -1.8093 |
| 1 | DKI | 0.2263 | 0.2261 | 940.7306 | <0.0001 | -0.8142 |
| 1 | DTI | 0.3575 | 0.3573 | 1789.7236 | <0.0001 | -1.0817 |
| 1 | SMT | 0.3462 | 0.3460 | 1841.6494 | <0.0001 | -1.0953 |
| 1 | SMT mc | 0.3152 | 0.3150 | 1480.6879 | <0.0001 | -1.0782 |
| 1 | WMTI | 0.3720 | 0.3718 | 1905.4699 | <0.0001 | -0.8474 |
| 2 | Full Multimodal | 0.0247 | 0.0244 | 81.5815 | <0.0001 | -0.0332 |
| 2 | Mean Multimodal | 0.0179 | 0.0177 | 65.2632 | <0.0001 | 0.0522 |
| 2 | BRIA | 0.0400 | 0.0397 | 134.1182 | <0.0001 | 0.1030 |
| 2 | DKI | 0.0191 | 0.0188 | 62.7254 | <0.0001 | 0.1090 |
| 2 | DTI | 0.1517 | 0.1514 | 575.2169 | <0.0001 | -0.0587 |
| 2 | SMT | 0.0022 | 0.0019 | 7.6338 | <0.0001 | -0.0159 |
| 2 | SMT mc | 0.0784 | 0.0781 | 273.5101 | <0.0001 | 0.2988 |
| 2 | WMTI | 0.0563 | 0.0560 | 191.8700 | <0.0001 | 0.0876 |
| 3 | Full Multimodal | 0.1148 | 0.1145 | 417.0057 | <0.0001 | -0.0086 |
| 3 | Mean Multimodal | 0.0003 | 0.0001 | 1.1795 | 0.2991 | -0.0023 |
| 3 | BRIA | 0.1842 | 0.1839 | 726.2248 | <0.0001 | 0.4627 |
| 3 | DKI | 0.2564 | 0.2562 | 1109.0911 | <0.0001 | 0.1705 |
| 3 | DTI | 0.2500 | 0.2498 | 1072.1366 | <0.0001 | -0.2096 |
| 3 | SMT | 0.2465 | 0.2463 | 1137.8183 | <0.0001 | 0.2908 |
| 3 | SMT mc | 0.1844 | 0.1841 | 726.9496 | <0.0001 | -0.1839 |
| 3 | WMTI | 0.1751 | 0.1749 | 682.7749 | <0.0001 | 0.1266 |
| 4 | Full Multimodal | 0.1279 | 0.1276 | 471.5012 | <0.0001 | 0.0002 |
| 4 | Mean Multimodal | 0.0999 | 0.0997 | 396.8655 | <0.0001 | 0.0672 |
| 4 | BRIA | 0.0880 | 0.0877 | 310.4268 | <0.0001 | -0.0049 |
| 4 | DKI | 0.0688 | 0.0685 | 237.5608 | <0.0001 | 0.0302 |
| 4 | DTI | 0.0696 | 0.0693 | 240.5501 | <0.0001 | -0.1682 |
| 4 | SMT | 0.1040 | 0.1038 | 403.8253 | <0.0001 | -0.0603 |
| 4 | SMT mc | 0.1639 | 0.1637 | 630.6907 | <0.0001 | 0.0803 |
| 4 | WMTI | 0.2239 | 0.2236 | 927.6834 | <0.0001 | 0.1036 |

**ST15.**
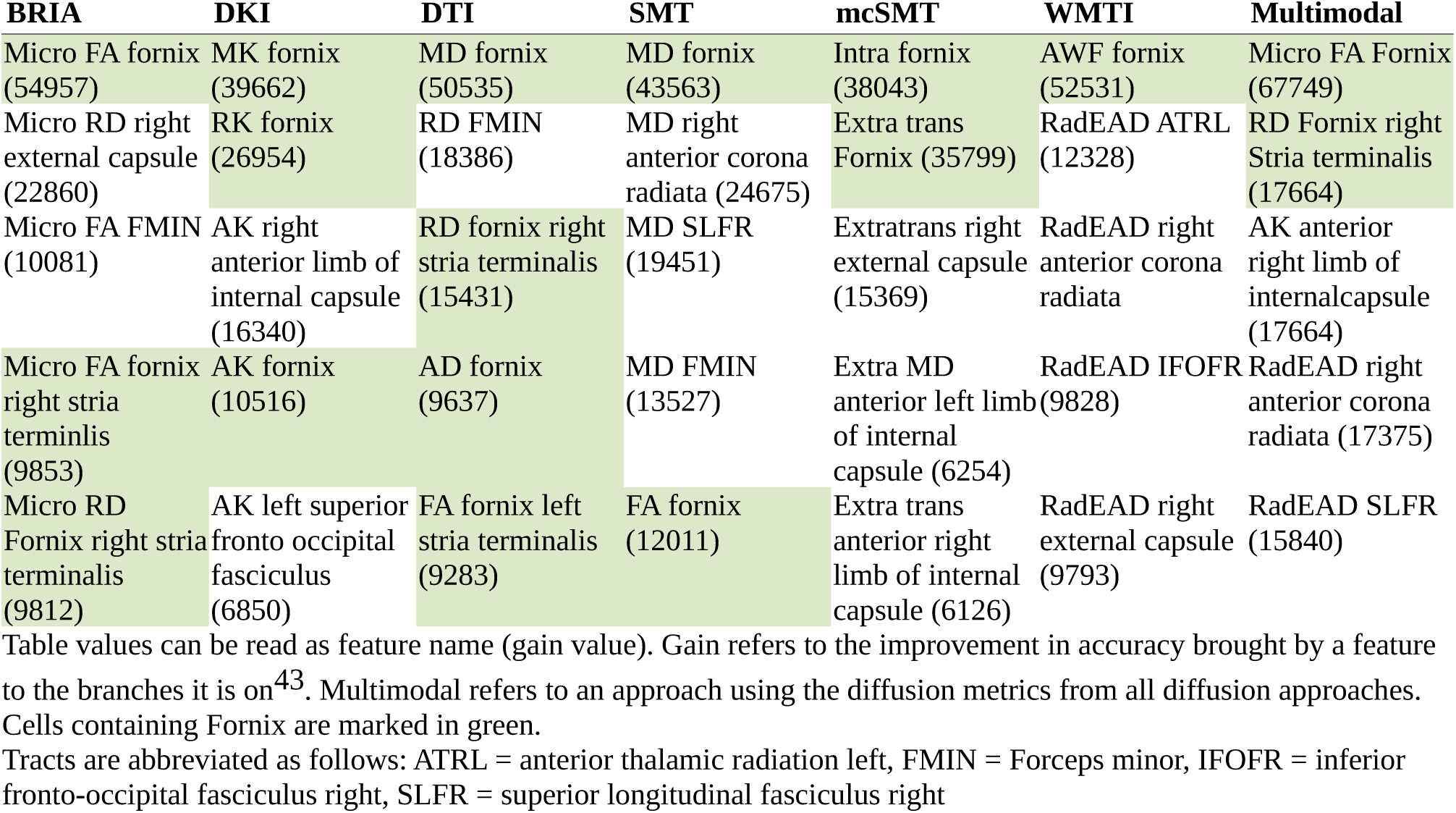
Top five diffusion metrics ranked by gain in age prediction accuracy.

| **BRIA** | **DKI** | **DTI** | **SMT** | **mcSMT** | **WMTI** | **Multimodal** |
| --- | --- | --- | --- | --- | --- | --- |
| Micro FA fornix (54957) | MK fornix (39662) | MD fornix (50535) | MD fornix (43563) | Intra fornix (38043) | AWF fornix (52531) | Micro FA Fornix (67749) |
| Micro RD right external capsule (22860) | RK fornix (26954) | RD FMIN (18386) | MD right anterior corona radiata (24675) | Extra trans Fornix (35799) | RadEAD ATRL (12328) | RD Fornix right Stria terminalis (17664) |
| Micro FA FMIN (10081) | AK right anterior limb of internal capsule (16340) | RD fornix right stria terminalis (15431) | MD SLFR (19451) | Extratrans right external capsule (15369) | RadEAD right anterior corona radiata | AK anterior right limb of internalcapsule (17664) |
| Micro FA fornix right stria terminlis (9853) | AK fornix (10516) | AD fornix (9637) | MD FMIN (13527) | Extra MD anterior left limb of internal capsule (6254) | RadEAD IFOFR (9828) | RadEAD right anterior corona radiata (17375) |
| Micro RD Fornix right stria terminalis (9812) | AK left superior fronto occipital fasciculus (6850) | FA fornix left stria terminalis (9283) | FA fornix (12011) | Extra trans anterior right limb of internal capsule (6126) | RadEAD right external capsule (9793) | RadEAD SLFR (15840) |

Table values can be read as feature name (gain value). Gain refers to the improvement in accuracy brought by a feature to the branches it is on<sup>43</sup>. Multimodal refers to an approach using the diffusion metrics from all diffusion approaches. Cells containing Fornix are marked in green. Tracts are abbreviated as follows: ATRL = anterior thalamic radiation left, FMIN = Forceps minor, IFOFR = inferior fronto-occipital fasciculus right, SLFR = superior longitudinal fasciculus right

**ST16.** Brain age prediction model performance e*xcluding fornix features* and uncorrected brain age – chronological age correlations comparison.

| Name | MRI features | R<sup>2</sup> | RMSE | MAE | Prediction-Age Correlation | Uncorrected Brain Age Correlation Difference to All Data |
| --- | --- | --- | --- | --- | --- | --- |
| BRIA | 670 | 0.527<br>(0.010) | 5.131<br>(0.042) | 4.129<br>(0.033) | 0.727<br>[0.722, 0.732] | -0.007*<br>[-0.009, -0.004] |
| DKI | 201 | 0.550<br>(0.015) | 5.108<br>(0.070) | 4.105<br>(0.065) | 0.742<br>[0.737, 0.747] | -0.006*<br>[-0.008, -0.003] |
| DTI | 269 | 0.555<br>(0.013) | 5.078<br>(0.066) | 4.079<br>(0.061) | 0.745<br>[0.745, 0.750] | -0.005*<br>[-0.007, -0.003] |
| SMT | 269 | 0.507<br>(0.008) | 5.347<br>(0.042) | 4.309<br>(0.028) | 0.713<br>[0.707, 0.718] | -0.009*<br>[-0.011, -0.006] |
| mcSMT | 269 | 0.488<br>(0.011) | 5.342<br>(0.045) | 4.303<br>(0.036) | 0.699<br>[0.693, 0.705] | -0.015*<br>[-0.018, -0.012] |
| WMTI | 201 | 0.566<br>(0.012) | 5.018<br>(0.062) | 4.031<br>(0.052) | 0.753<br>[0.748, 0.757] | -0.003*<br>[-0.006, -0.001] |
| Mean scores multimodal | 28 | 0.393<br>(0.012) | 5.932<br>(0.051) | 4.812<br>(0.046) | 0.627<br>[0.621, 0.634] | 0<br>[-0.0001, 0.0001] |
| Full model multimodal | 1876 | 0.636<br>(0.012) | 4.591<br>(0.077) | 3.677<br>(0.039) | 0.798<br>[0.794, 0.802] | -0.006*<br>[-0.007, -0.004] |

Multimodal mean scores remain unaffected as averaged across the brain.
\* p<.001

**ST17.**
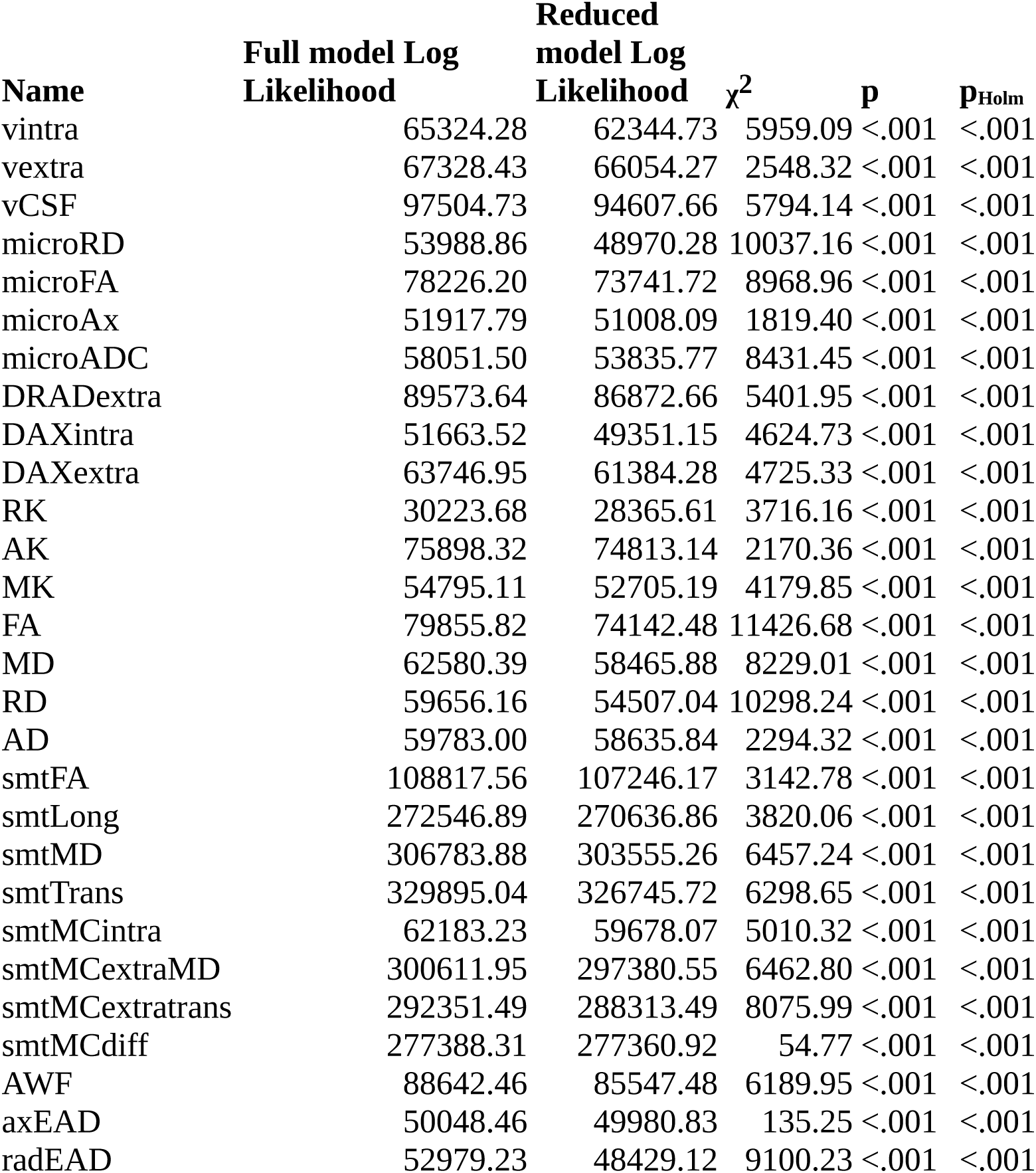
Forceps age-sensitivity.

**ST18:** Comparisons of linear and generalized additive models predicting forceps diffusion metrics.

| Metric | LM AIC | GAM AIC | LM BIC | GAM BIC | LM<br>R<sup>2</sup><sub>adj</sub> | GAM<br>R<sup>2</sup><sub>adj</sub> |
| --- | --- | --- | --- | --- | --- | --- |
| vintra | -130624.56 | -130624.65 | -130522.74 | -130531.33 | 0.16 | 0.16 |
| vextra | -134632.86 | -134634.69 | -134531.05 | -134541.36 | 0.08 | 0.08 |
| vCSF | -194985.46 | -194983.16 | -194883.65 | -194889.84 | 0.17 | 0.17 |
| microRD | -107953.72 | -107948.75 | -107851.91 | -107855.43 | 0.26 | 0.26 |
| microFA | -156428.39 | -156426.02 | -156326.58 | -156332.69 | 0.23 | 0.23 |
| microAx | -103811.58 | -103813.40 | -103709.77 | -103720.08 | 0.08 | 0.08 |
| microADC | -116078.99 | -116076.52 | -115977.18 | -115983.19 | 0.23 | 0.23 |
| DRADextra | -179123.28 | -179125.07 | -179021.47 | -179031.74 | 0.15 | 0.15 |
| DAXintra | -103303.03 | -103305.01 | -103201.22 | -103211.69 | 0.14 | 0.14 |
| DAXextra | -127469.89 | -127471.55 | -127368.08 | -127378.22 | 0.14 | 0.14 |
| RK | -60423.37 | -60425.09 | -60321.56 | -60331.76 | 0.13 | 0.13 |
| AK | -151772.64 | -151769.08 | -151670.82 | -151675.76 | 0.09 | 0.09 |
| MK | -109566.23 | -109567.55 | -109464.41 | -109474.23 | 0.12 | 0.12 |
| FA | -159687.65 | -159688.21 | -159585.83 | -159594.89 | 0.28 | 0.28 |
| MD | -125136.77 | -125133.28 | -125034.96 | -125039.96 | 0.22 | 0.22 |
| RD | -119288.32 | -119288.44 | -119186.51 | -119195.11 | 0.26 | 0.26 |
| AD | -119542.00 | -119530.17 | -119440.19 | -119436.84 | 0.09 | 0.09 |
| smtFA | -217611.12 | -217610.05 | -217509.31 | -217516.72 | 0.10 | 0.10 |
| smtLong | -545069.78 | -545071.74 | -544967.97 | -544978.42 | 0.13 | 0.12 |
| smtMD | -613543.77 | -613545.40 | -613441.96 | -613452.07 | 0.18 | 0.18 |
| smtTrans | -659766.09 | -659763.06 | -659664.28 | -659669.73 | 0.17 | 0.17 |
| smtMCintra | -124342.46 | -124343.29 | -124240.65 | -124249.96 | 0.13 | 0.13 |
| smtMCextraMD | -601199.89 | -601200.16 | -601098.08 | -601106.83 | 0.19 | 0.19 |
| smtMCextratrans | -584678.97 | -584678.64 | -584577.16 | -584585.31 | 0.21 | 0.21 |
| smtMCD | -554752.62 | -554754.61 | -554650.81 | -554661.29 | 0.03 | 0.03 |
| AWF | -177260.92 | -177262.82 | -177159.11 | -177169.50 | 0.17 | 0.17 |
| axEAD | -100072.91 | -100063.70 | -99971.10 | -99970.37 | 0.03 | 0.03 |
| radEAD | -105934.46 | -105934.45 | -105832.65 | -105841.12 | 0.25 | 0.24 |

LM = linear model, GAM = generalized additive model, AIC = Akaike information criterion, BIC = Bayesian information criterion. The numbers are derived from the six diffusion approaches' 28 metrics following Equation 1 for linear models and all variables of the equation allowing splines for non-linear models.

